# Models for Eco-evolutionary Extinction Vortices and their Detection

**DOI:** 10.1101/2020.02.28.969402

**Authors:** Peter Nabutanyi, Meike J. Wittmann

## Abstract

The smaller a population is, the faster it looses genetic variation due to genetic drift. Loss of genetic variation can reduce population growth rate, making populations even smaller and more vulnerable to loss of genetic variation, and so on. Ultimately, the population can be driven to extinction by this “eco-evolutionary extinction vortex”. So far, extinction vortices due to loss of genetic variation have been mainly described verbally. However, quantitative models are needed to better understand when such vortices arise and to develop methods for detecting them. Here we propose quantitative eco-evolutionary models, both individual-based simulations and analytic approximations, that link loss of genetic variation and population decline. Our models assume stochastic population dynamics and multi-locus genetics with different forms of balancing selection. Using mathematical analysis and simulations, we identify parameter combinations that exhibit strong interactions between population size and genetic variation as populations decline to extinction and match our definition of an eco-evolutionary vortex, i.e. the per-capita population decline rates and per-locus fixation rates increase with decreasing population size and number of polymorphic loci. We further highlight cues and early warning signals that may be useful in identifying populations undergoing an eco-evolutionary extinction vortex.

## 1 Introduction

The fate of small populations is often strongly affected by demographic stochasticity and genetic factors. Small populations can rapidly loose genetic variation due to genetic drift. Additionally, inbreeding, i.e. mating between relatives, is more likely in small than large populations. In captive and natural populations, inbreeding and reduced heterozygosity are often associated with a reduction in population fitness or in a fitness component (Saccheri et al. 1998; Reed and Frankham 2003). For instance, reduced heterozygosity is associated with reduced hatching success and increased mortality rates of chicks in a metapopulation of southern dunlins (Blomqvist et al. 2010). Inbreeding is also associated with higher mortality rates in captive ungulates (Ralls et al. 1979; Ballou and Ralls 1982), lower breeding recruitment rate in wolves (Bensch et al. 2006), reduced litter size in the Iberian Lynx (Palomares et al. 2012), and reduced disease resistance in *Drosophila melanogaster* and the Tasmanian devil (Spielman et al. 2004; Miller et al. 2011).

In summary, small populations experience genetic problems such as loss of variation and inbreeding depression, and these genetic problems can reduce population fitness. The reduction in population fitness components can cause a further decline in population size, thereby exposing the population to even more severe genetic problems. Shrinking population size and genetic problems can thus form an extinction vortex (Gilpin and Soulé 1986), a positive feedback loop that can ultimately drive the population to extinction. Gilpin and Soulé (1986) proposed four types of extinction vortices depending on the major factors driving the feedback loop. The R vortex is driven by the feedback between decreasing population size and increasing variance in growth rate, e.g. because of more variable sex ratio and resulting mate-finding problems. For the D vortex, the feedback is mediated by increasing fragmentation of the species distribution at small population size. The F and A vortices are mainly driven by genetic factors such as genetic drift, loss of heterozygosity and generally loss of genetic variation. In particular, the F vortex is driven by inbreeding depression and loss of heterozygosity and less dependent on the environment while the A vortex involves reduction of a population’s adaptive potential in new environments. Our study is closely related to the F and A vortices and to distinguish them from other vortices, e.g. those caused by anthropogenic Allee effect (Courchamp et al. 2006), we refer to extinction due to feedback loops between genetic deterioration and declining population sizes as an “eco-evolutionary vortex”.

Genetic problems in small populations can be divided into three categories: inbreeding depression, mutation accumulation and mutational meltdown, and loss of genetic variation and evolutionary potential (Frankham 2005). It is possible in principle that each of them alone or any combination of them can give rise to an eco-evolutionary vortex. Such extinction vortices are often described verbally in the literature. However, to be able to understand the conditions under which populations can enter an extinction vortex and develop methods to detect populations in an extinction vortex, we need quantitative models. So far, progress on such quantitative models has mostly been made for the two of the three types of genetic problem: inbreeding depression and mutation accumulation. For example, inbreeding depression can cause an “inbreeding vortex” (Tanaka 1997, 1998, 2000) which occurs when a large population with its relatively high frequency of recessive deleterious alleles is suddenly reduced in size, leading to more mating between relatives. Inbreeding depression can also give rise to a genetic Allee effect, with a rapid change in extinction probability around some critical population size (Wittmann et al. 2018). Furthermore, inbreeding depression is often included in population viability analyses, e.g. in a recent study on mountain lions by Benson et al. (2019) and many studies using the software VORTEX (Lacy 1993). For mutation accumulation, eco-evolutionary models have been developed to show how time to extinction depends on the population’s carrying capacity (Lynch and Gabriel 1990) and to quantify the strength of the mutational meltdown compared to a scenario without eco-evolutionary feedbacks (Coron et al. 2013). However, for the third type of genetic problem in small populations, loss of genetic variation and evolutionary potential, to our knowledge, there is so far no quantitative model for the potentially resulting extinction vortex. In this paper, we thus aim to develop a quantitative model for an extinction vortex driven by the positive feedback between loss of genetic variation and reduction in population size.

Because of a lack of appropriate detection methods, the contribution of genetic problems to extinction of endangered populations may often be underrated or remain unnoticed. Using our model, we want to identify key features and cues that can be used to identify natural populations caught in an extinction vortex driven by loss of genetic variation. Regarding the detection of extinction vortices in general, a retrospective analysis of demographic data from 10 already extinct wildlife populations showed that both year-to-year rates of population decline and variance in size increased as extinction was approached (Fagan and Holmes 2006). For extinction vortices due to loss of genetic variation, it would be useful to have measures of relevant levels of genetic variation. However, we generally do not know which loci contribute to fitness and how many such loci there are. Therefore, we also evaluate whether extinction vortices can be detected by some general early-warning signals for extinction in changing environments (Drake and Griffen 2010; Dakos and Bascompte 2014; Jarvis et al. 2016; Sommer et al. 2017; de Silva and Leimgruber 2019). As the environment gradually deteriorates, the commonly used early-warning statistical measures such as autocorrelation, standard deviation, coefficient of variation and kurtosis are expected to increase near bifurcation points where the population will shift to a new equilibrium, while skewness either increases if the new equilibrium point is higher or decreases otherwise (see Dakos et al. (2012) for an overview). In our case, the magnitude and stability of equilibria is not affected by a deteriorating external environmental factor, but genetic variation as an internal factor.

Here we develop stochastic individual-based eco-evolutionary models and analytic approximations for the feedback between loss of of genetic variation and population decline. We assume multi-locus genetics and focus on scenarios where genetic variation can be maintained in large populations due to some form of balancing selection, but is at risk of being lost due to genetic drift in small populations. First, we partition the parameter space into regions with qualitatively different eco-evolutionary behaviour. We then pick exemplary cases from each region to check for the presence of an eco-evolutionary vortex, which we define by two key features: (i) the per-capita rate of population decline increases with both decreasing genetic variation and declining population size and (ii) the per-locus rate of loss of genetic variation increases with declining population size as well as decreasing genetic variation. In addition, we use our model to test for three hypothesized cues of extinction vortices: (i) there exists a critical population size and (or) critical level of genetic variation below which population decline and (or) loss of genetic variation suddenly increases substantially, (ii) loss of genetic variation and population decline and extinction occur on the same time scale and, (iii) early-warning signals manifest as population size decreases.

## 2 Methods

In this section, we describe two different approaches to modeling the interplay between loss of genetic variation and population decline: an individual-based model (IBM) and an analytic approximation. Individuals in our model are diploid, hermaphroditic and have *n* unlinked loci, each with two alleles, *a* and *A*.

### 2.1 Individual-based Model

We start with a population size drawn from a Poisson distribution with mean *K*, the carrying capacity. The alleles are initially drawn with equal probability across all loci of each individual. For each individual at generation *t*, the actual number of offspring is independently drawn from a Poisson distribution with mean

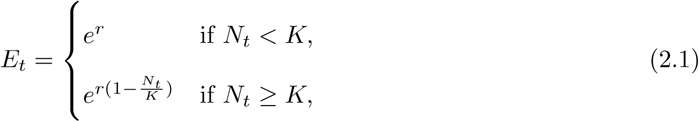

where *r* and *N*_*t*_ are the maximum intrinsic growth rate and population size at generation *t* respectively. In this model, the population grows geometrically when below *K*, but at *K* and larger sizes follows the Ricker model to prevent population explosion. The assumption that small populations do not experience any density dependence simplifies the mathematical analysis and should be a good approximation for many small endangered populations. However, we also consider a fully density-dependent model, where we only use the second part of Equation (2.1), i.e. the Ricker model, at all population sizes.

Each offspring produced by the focal parent randomly chooses the second parent from all the available parents in the population, including the focal parent. For each locus, the offspring inherits one randomly chosen allele from each parent.

We consider two selection mechanisms, both of which give rise to balancing selection such that in a large population both alleles can be maintained. The first mechanism is heterozygote advantage, where heterozygotes have higher fitness than either homozygote. In this paper, both homozygotes have the same fitness (*w*_*AA*_ = *w*_*aa*_ = 1 − *s*, where 0 < *s* < 1 is the selection coefficient) while heterozygotes have maximum fitness (*w*_*Aa*_ = 1). The second mechanism is fluctuating selection with reversal of dominance (Wittmann et al. 2017; Bertram and Masel 2019; Connallon and Chenoweth 2019). Recent studies have suggested dominance reversals, e.g. in *Drosophila melonagaster* (Chen et al. 2015) and beetles (Grieshop and Arnqvist 2018). Specifically, we assume that the fitness of homozygote genotypes fluctuates temporally. At each generation, the fitness of heterozygotes is intermediate between that of the two homozygotes, but closer to the currently fitter homozygote (Figure 1), i.e. there is beneficial reversal of dominance (Curtsinger et al. 1994). This causes the heterozygote genotype to have a higher geometric mean fitness than either homozygote, and balancing selection emerges. We achieve this by setting: *w*_*AA,t*_ = (1−*s*_*A,t*_)/(1+*s*), *w*_*aa,t*_ = (1−*s*_*a,t*_)/(1+*s*) and *w*_*Aa,t*_ = (1 − *h*_*t*_ · *s*_*A,t*_ − (1 − *h*_*t*_) · *s*_*a,t*_)/(1 + *s*), where *s*_*A,t*_ = *s* · sin(2*π* · *t/κ*) and *s*_*a,t*_ = *s* · sin(*π* + 2*π* · *t/κ*) are temporally fluctuating selection coefficients and *h*_*t*_ = 0.5 − *c* · sin(2*π* · *t/κ*) is the temporally fluctuating dominance coefficient. In this model, *s* determines the amplitude of fluctuating selection, *κ* is the number of generations in a complete cycle (we use *κ* = 50 throughout the paper), and 0 ≤ *c* ≤ 0.5 determines the magnitude of dominance changes. The division by 1 + *s* ensures that all fitness values are between 0 and 1. We also run a neutral control scenario where all the three genotypes have maximum fitness of 1 (i.e. selection coefficient *s* = *s*_*a,t*_ = *s*_*A,t*_ = 0) at all generations.

**Figure 1:**
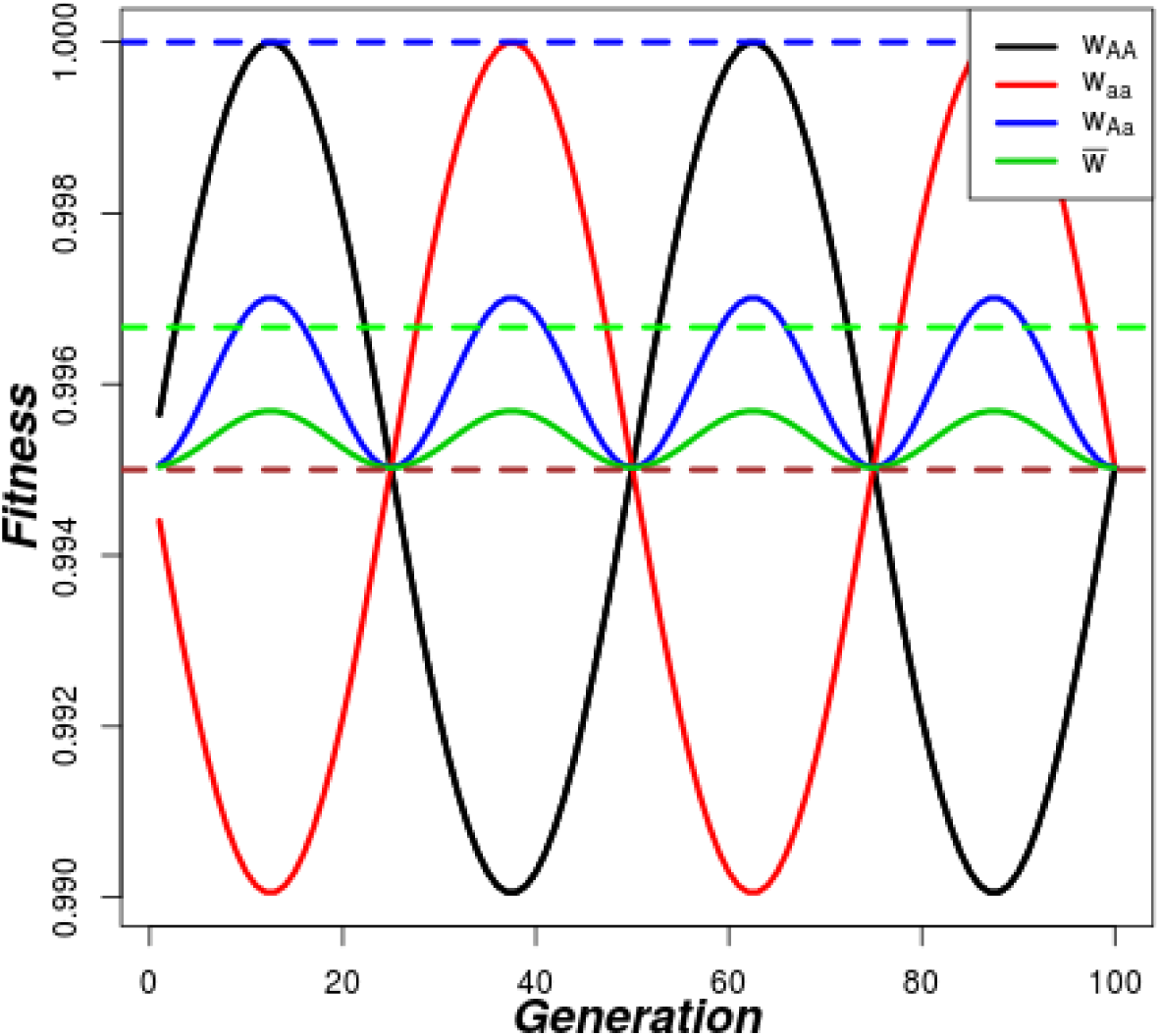
Example of fitness trajectories under fluctuating selection with reversal of dominance. The solid black, red and blue lines represent the fluctuating fitness for homozygotes *AA* and *aa* and heterozygote *Aa* genotypes, respectively, while the green-solid line is for mean population fitness. For the heterozygote advantage mechanism, the dashed blue and brown lines represent fitnesses of heterozygote and homozygote genotypes, respectively, while the green-dashed line is the mean population fitness. The parameters are *s* = 0.005, *c* = 0.2, *κ* = 50.

Denoting *w*_*g,t*_ as the fitness of genotype *g* ∈ {*AA, Aa, aa*} at generation *t* and assuming multiplicative fitness across loci, each offspring is viable with probability

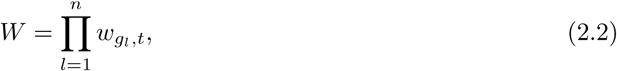

where *g*_*l*_ is the genotype at locus *l*. After all individuals have reproduced, all the viable offspring then replace the parent generation. In both selection mechanisms, mutation is not considered and therefore an allele is not reintroduced once lost at a given locus. The current number of polymorphic loci, i.e. loci where both alleles are still present in the population, is denoted *H*_*t*_.

### 2.2 Parameter space

Two key model parameters are the intrinsic growth rate, *r*, which determines the fertility rate and therefore population growth rate, and the selection coefficient, *s*, which determines the offspring viability and influences the rate at which genetic variation is lost. To determine regions in the *r* − *s* plane with qualitatively different behavior, we consider the average number of surviving offspring per individual (*SOI*), which is given by the product of the average number of offspring per individual *E* (see Equation 2.1) and the average viability, i.e. fitness, of offspring at a given level of genetic variation (*W*(*H*)). To obtain a “critical number of polymorphic loci”, *H*_*c*_, below which population crash to zero occurs, we set *SOI* to 1, the boundary point between population growth and decline, and solve for *H* (see Appendix Equation A.26).

For the heterozygote advantage mechanism where both homozygotes have the same fitness 1 − *s* and heterozygotes have fitness 1, the mean fitness for locus *l* with allele frequency *x*_*l*_ is

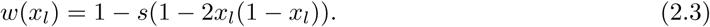

We then approximate the fitness in terms of polymorphic loci by assuming an allele frequency of 0.5 at all polymorphic loci, the expected equilibrium frequency under symmetric heterozygote advantage.

The fitness, *W*(*H*) is thus given by

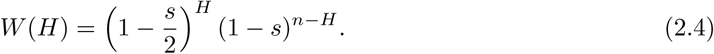

Since the assumption of an allele frequency of 0.5 is generally violated, fitness is overestimated and the critical number of loci is thus underestimated to some extent by this approach. For fluctuating selection with reversal of dominance, the geometric mean fitness of the genotypes is used to calculate the mean population fitness for a given locus. Briefly, we obtain the geometric mean fitness for the three genotypes (*AA, Aa* and *aa*) denoted as *G*_*AA*_, *G*_*Aa*_ and *G*_*aa*_ respectively. In this study, *s*_*A*_ = *s*_*a*_, and therefore *G*_*AA*_ = *G*_*aa*_ = *G*_*h*_, where *G*_*h*_ denotes the geometric mean fitness of either homozygote. Also, assuming an equilibrium frequency of 0.5 and multiplicative fitness across loci, the population fitness becomes

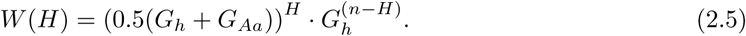

### 2.3 Analysis

We run a total of 100 replicate populations and each replicate population is iterated until either the population goes extinct or appropriate maximum number of generations is reached. The maximum simulation time is chosen in such a way that all polymorphic loci are at least lost in the population. In this study, maximum simulation time of 3,000 generations was sufficient for all cases involving *N*_0_ = *K* = 200 while 30,000 was used for *N*_0_ = *K* = 2000. In our analysis, we define the level of genetic variation in terms of the number of polymorphic loci in the population.

For each replicate independently, the per-capita decline rate is calculated as 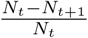 for each generation *t*. The range of population sizes is divided into equally spaced bins of width 10 (population sizes 1 to 10, then 11 to 20,…). For each replicate and bin separately, we then calculate the mean population size and the corresponding mean per-capita rate. To get overall mean population size and mean per-capita rate for each bin, we take the means across all contributing replicates, i.e. replicates that have data points in the respective bin. In the same way, we calculate the per-locus rate of loss of polymorphic loci, 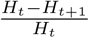, first for each replicate and each generation. Then we averaged, using the same binning procedure as for population size above and with a bin width of 10. For the full Ricker model where relatively large carrying capacity is used, the bin width for population size is 50, while that of polymorphic loci is maintained at 10.

To calculate early-warning signals (auto-correlation at lag 1, coefficient of variation, skewness and kurtosis), the population size is first log-transformed due to presence of extreme values and values close or equal to zero (Dakos et al. 2012). The NAs produced as a result of population extinction are removed before the indicators are computed. Using an overlapping moving window length of 100 generations (200 generations for the Ricker model with larger carrying capacity), we calculate each indicator for each replicate separately, starting with the 100th generation. Here, the most recent available 100 data points are used to estimate each indicator at each generation point until the last generation of population existence. Now, following the same binning procedure described above in calculating per-capita decline rates, we calculate the indicator from the 100th generation (200th generation for Ricker model) to population extinction as a function of population size.

### 2.4 Analytic Approximation of Eco-Evolutionary Model

The individual-based model is approximated using a difference equation (for population size dynamics) coupled with a classical diffusion approximation equation (for the allele frequencies). The first part of the eco-evolutionary model represents the population dynamics as

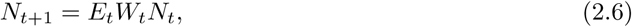

where *E*_*t*_ is the average number of offspring per individual given by Equation (2.1) and *W*_*t*_ is the average population fitness determined by the distribution *f*(*x, t*), of allele frequencies, *x*, at generation *t*, as outlined below.

We approximate *f*(*x, t*) using a diffusion equation (e.g in Kimura et al. (1955))

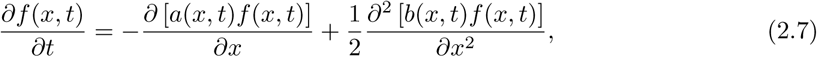

where *a*(*x, t*) and *b*(*x, t*) refer to the infinitesimal mean and infinitesimal variance of the change in *x*. In the case of heterozygote advantage, *a*(*x, t*) = *sx*(1 − *x*)(1 − 2*x*) and 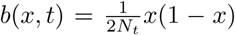 (e.g see Durrett (2008)), while in our fluctuating selection scenario, *a*(*x, t*) = *x*(1 − *x*)(*x*(1 − 2*h*_*t*_) + *h*_*t*_)(1 + *s*_*a,t*_)(*s*_*a*_ − *s*_*A,t*_) (see Appendix A.2) and 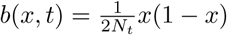.

We numerically obtain a complete solution to Equation (2.7) subject to the initial and boundary conditions as in Zhao et al. (2013) and Xu et al. (2019) (see Appendix Section A.1 for details). In brief, the allele frequency *x* is discretized using a grid with grid points *x*_*i*_ = *i* · *u*, where *i* = 0, 1, …, *m* and *u* = 1*/m*. Similarly, time is discretized with spacing *τ* = 1 corresponding to a full generation with grid points *t*_*k*_ = *v* · *τ, v* = 0, 1,…. We assume that all loci have initial allele frequency *x*_0_. The boundary condition is obtained by imposing the assumption that the system conserves probability at every time step (sum of all probabilities = 1, for all *t*). Thus, no probability flows outside the system as loss or fixation of alleles is captured by the outermost bins (*x* = 0 and *x* = 1). As a final result, we obtain a vector of probabilities *f*(*x*_*i*_, *t*) with *i* ∈ {0, 1, …, *m*} for each time point.

For each time *t* > 0, the probability density *f*(*x, t*) is used to estimate

a. the population mean fitness, *W*_*t*_. With the assumption of multiplicative fitness across all *n* unlinked loci, 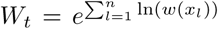, where *w*(*x*) is given by Equation 2.3. Because the exact allele frequency *x*_*l*_ at a given locus is not explicitly known, we approximate *W*_*t*_ using the expectation of fitness at a particular locus over the distribution *f*(*x, t*). Assuming that *n* is large and all loci independently sample an allele frequency from the current allele-frequency distribution,

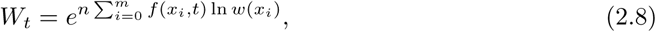

where the sum is over all allele-frequency bins, and
b. the probability that both alleles are still present, 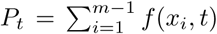. This is then used to estimate the number of polymorphic loci as,

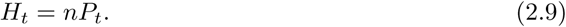

Note that (2.8) together with (2.6) and the numerical solution for *f*(*x,t*) completes the specification of the eco-evolutionary model, while *H*_*t*_ is used as a summary measure of genetic variation at that time.

### 2.5 Data Accessibility

All simulations and analyses were done in the *R* programming language (R Core Team 2015) and the R scripts are provided in the supplementary material.

## 3 Results

### 3.1 Parameter space

We generally obtain three distinct regions in the intrinsic growth rate - offspring viability selection (*r* − *s*) plane with qualitatively distinct behavior (Figure 2). Region I consists of populations with a critical number of polymorphic loci *H*_*c*_ above the total number of loci *n*. This implies that the number of polymorphic loci is below *H*_*c*_ from the start and populations are expected to decline to extinction immediately. In Region III, *H*_*c*_ < 0. This implies that populations in this region can loose all their polymorphic loci without rapid population decline. However, in Region II, 0 ≤ *H*_*c*_ ≤ *n*. In this region, populations are expected to start to decline faster at some intermediate number of polymorphic loci. For fluctuating selection, we observe the same three regions in parameter space (Figure 2B), but Region II is more narrow than for the heterozygote advantage mechanism (Figure 2A). To look at the detailed features in each region, we simulate exemplary cases from each region. In this main text, we present results from Cases 1, 2 and 3 and results from cases labelled *s* · · are given in the appendix.

**Figure 2:**
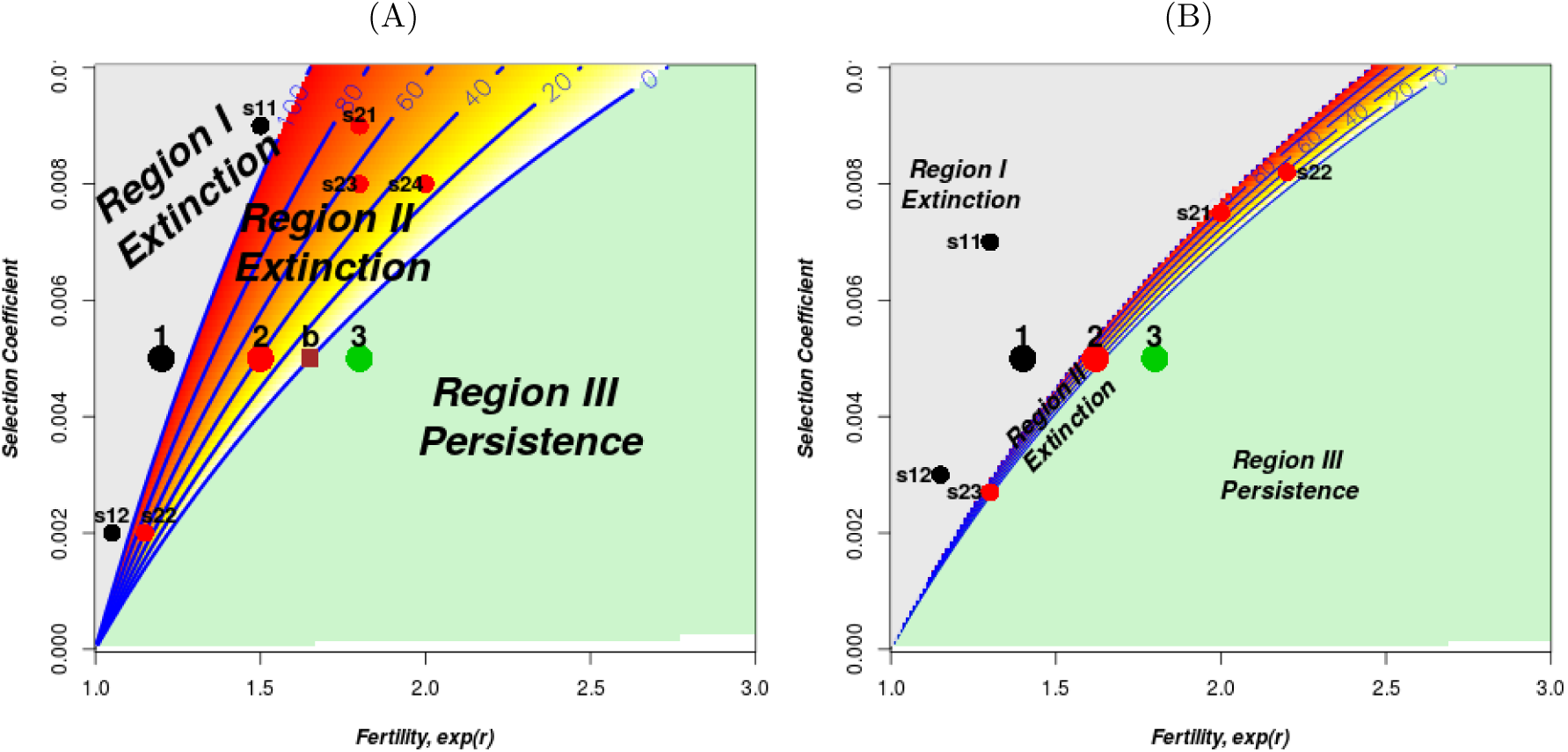
Parameter space divided into three major regions according to whether the population declines even with all loci polymorphic (Region I), starts declining at an intermediate number of polymorphic loci (Region II) or can persist even without polymorphic loci (Region III). (A) is for heterozygote advantage, and (B) is for fluctuating selection (see Section 2.2 for details). The red spectrum and blue lines show the critical number of polymorphic loci below which rapid population decline to extinction occurs. The points numbered 1, 2 and 3 are the cases considered in the main text while the points prefixed with *s* are supplementary cases considered in the appendix. The brown square labelled *b* is a boundary point in Region III but very close to Region II. The parameters used are *n* = 100, *c* = 0.2, *κ* = 50, *K* = 200.

### 3.2 Heterozygote Advantage

Figure 3 shows population trajectories of Cases 1, 2 and 3 representing Regions I, II and III in Figure 2A, respectively, and the neutral control case where all genotypes have the maximum fitness of 1. The dynamics from the analytic approximations fall within the range of the realisations of the IBM in the respective cases (dark blue lines in Figure 3, see Appendix Figure A.1 for the corresponding allele-frequency distributions at different times). For Case 1 where fertility is too low to compensate for viability selection, populations start their decline to extinction even before losing any genetic variation (Figures 3A and 3E). The number of polymorphic loci rapidly decreases to zero at or shortly before population extinction. In Case 3 where fertility is high enough to compensate for viability selection, even after total loss of polymorphic loci, populations basically fluctuate around a population size close to *K* (Figures 3C and 3G). However, in Case 2, we observe that populations start decreasing slowly with fluctuations until a certain point where they rapidly decline to extinction (Figure 3B). Similarly, after a constant phase, polymorphic loci are lost one by one at a small rate until a certain number of polymorphic loci is reached and then a rapid loss of all remaining polymorphic loci is observed (Figure 3F). The critical number of polymorphic loci below which rapid loss occurs roughly matches the estimated number from the analytic parameter space analysis (both around 40). These turning points agree with the suggested cue about existence of threshold values in levels of genetic variation and population size below which populations suddenly decline rapidly to extinction. No such turning points are observed in Cases 1, 3, or the neutral Control case. Also, in line with the proposed cue about similar time scales, we observe that population extinction and loss of polymorphic loci occur at more or less the same time in Cases 1 and 2. The populations survive for longer periods of time after total loss of polymorphic loci in both Case 3 and the Control case scenario (leading to different time scales). In the neutral control, populations simply fluctuate around *K* while polymorphic loci are lost one by one due to drift until complete fixation occurs (Figures 3D and 3H). Note that the neutral case has the same fertility as Case 2, but although there is no balancing selection in the neutral case, polymorphism is lost more slowly.

**Figure 3:**
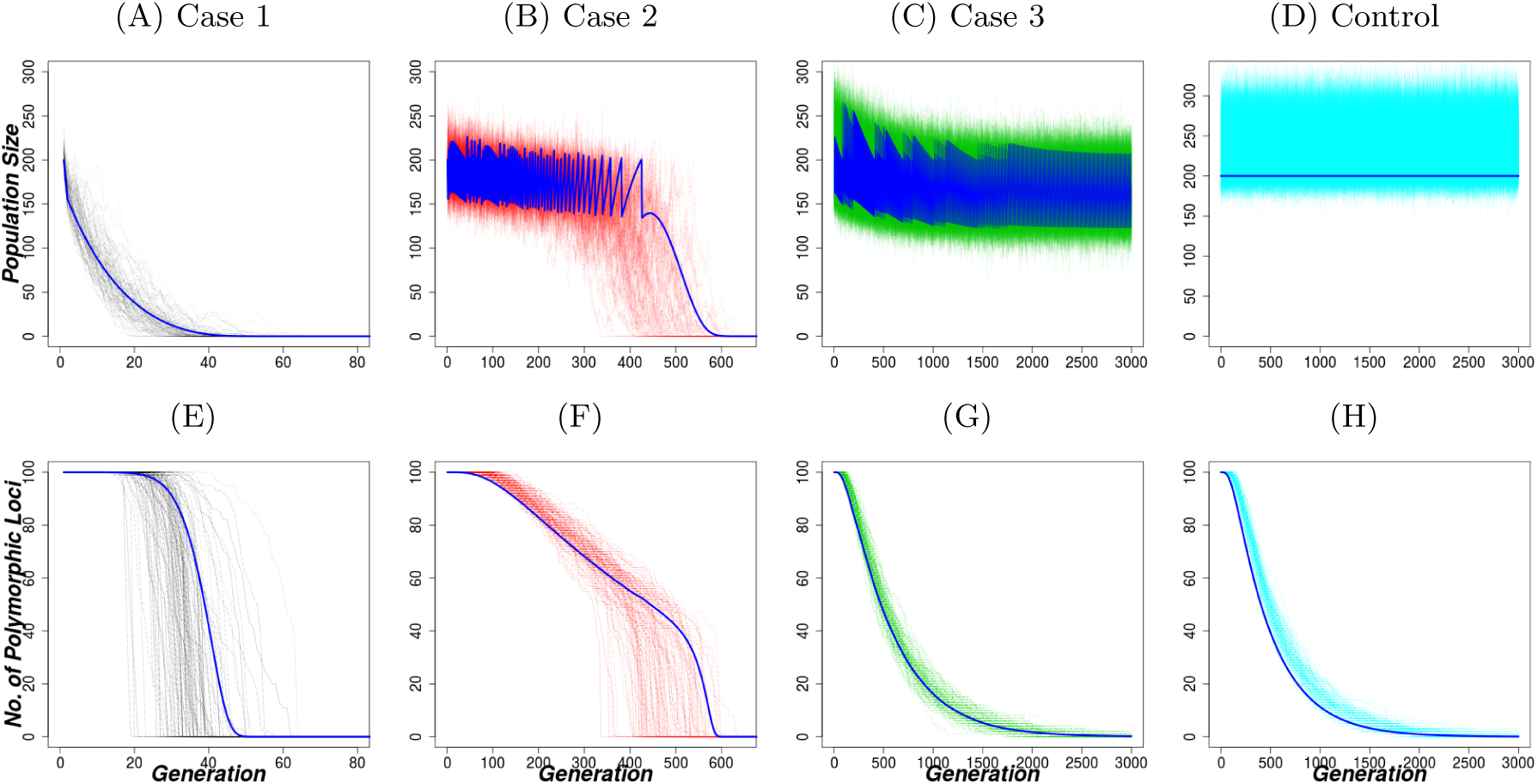
Dynamics of population size and genetic variation under the geometric population growth model with heterozygote advantage. The 1st 3 column panels are for different cases as indicated on Figure 2A. 1st column (Case 1): *r* = ln(1.2), 2nd column (Case 2): *r* = ln(1.5), and 3rd column (Case 3): *r* = ln(1.8). Other parameters are *s* = 0.005, *N*_0_ = *K* = 200, *replicates* = 100. 4th column panel is the control case with same parameters as in 2nd column panel but *s* = 0. The dark blue lines show the analytic approximation for each case with discretization parameters *u* = 0.1 and *τ* = 1. Note the different scaling of the time-axis on the different plots.

Figure 4 shows how the per-capita population decline rates and the per-locus rates of loss of polymorphic loci vary with decreasing number of polymorphic loci and declining population size. When population size is below *K*, the per-capita rate of population decline increases with declining population size and decreasing number of polymorphic loci for Cases 1 and 2 (Figures 4A and 4B). The rate is more or less constant (and positive) as population size decreases in Case 1 with a marked increase near population extinction. In Case 2, the per-capita decline rate increases gradually from negative to positive as population size (below *K*) and number of polymorphic loci decrease, which further supports the existence of suggested threshold values of population size and genetic variation. The negative per-capita decline rate implies positive per-capita growth rate. In Case 3, the per-capita decline rate is almost constant as both population size and number of polymorphic loci decrease. Note that the per-capita decline rate is relatively high and positive whenever population size exceeds *K* (grey-shaded region). This is because our geometric model with a Ricker boundary forces populations to decrease whenever *N*_*t*_ > *K* (see Equation (2.1)).

**Figure 4:**
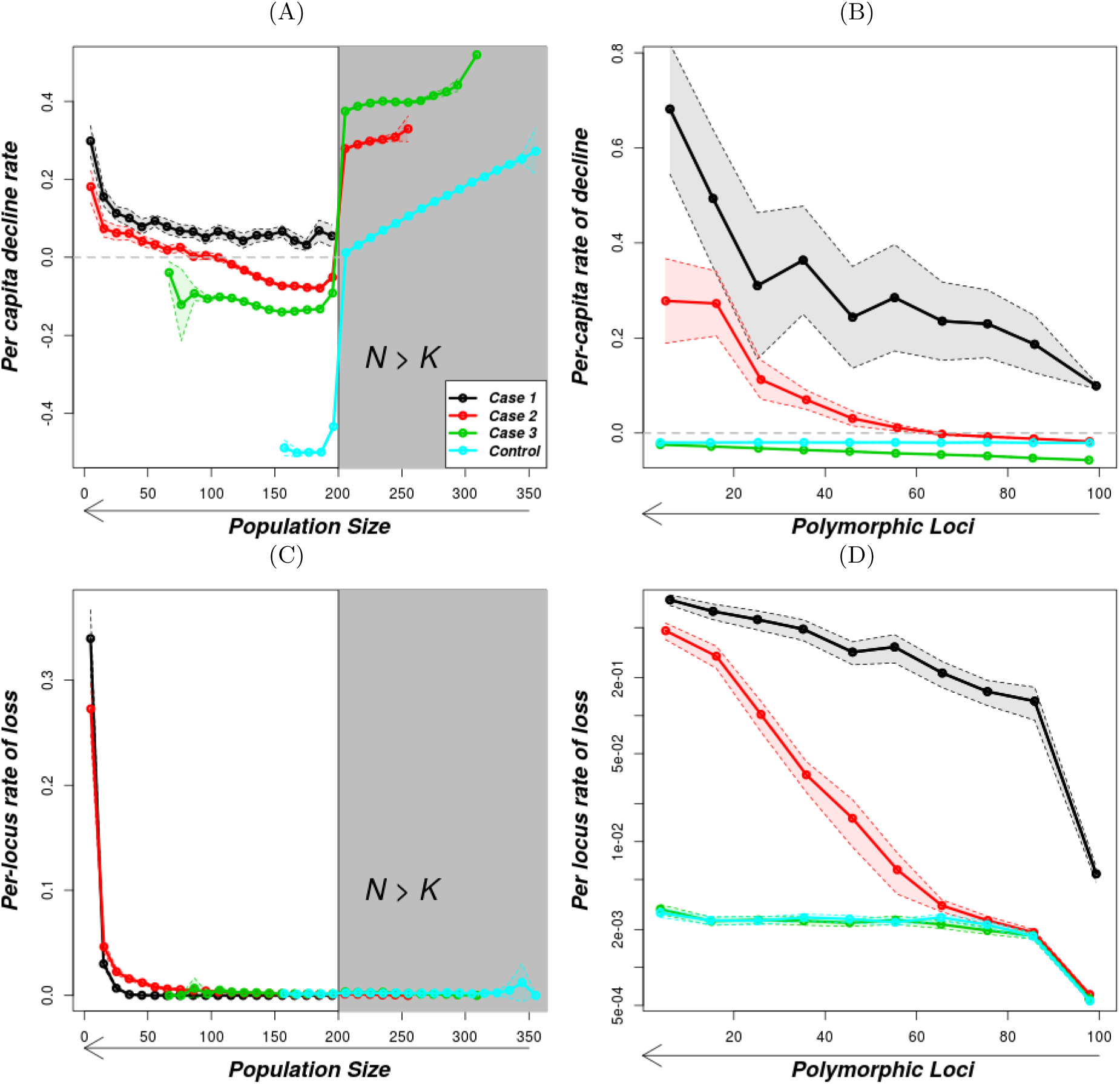
Per-capita rates of population decline and per-locus rates of loss of polymorphic loci under the geometric population growth model with heterozygote advantage. The gray-shaded part in (A) and (C) is the region above *K*. The solid lines are the mean values of the rates as described in the methods and the shaded regions shows the standard error. The horizontal gray line in (A) and (C) is for rate 0. The parameters for the 4 cases are as in Figure 3. In (D), the y-axis is on a logarithmic scale. The arrows indicate the long-term direction of change over time.

Similarly, as population size decreases, the per-locus rate of loss of polymorphic loci first increases gradually for Case 2 until a certain size is reached where the rate increases suddenly and substantially (Figure 4C). For Cases 1 and 3, the rate is more or less constant near zero, but when the population is near extinction, a sudden increase in per-locus rate of loss for Case 1 occurs. Also, as the number of polymorphic loci decreases, the per-locus rate of loss increases for Cases 1 and 2 but is roughly constant for Case 3 (Figure 4D). In summary, Cases 1 and 2 match our definition of an an eco-evolutionary vortex, i.e. per-capita decline rate and per-locus rate of loss increase as both population size and number of polymorphic loci decrease. Other cases in Regions I and II exhibit the same features as Case 1 and 2 (see Figure A.4). Cases 3 and the Control scenario, by contrast, do not match the definition.

### 3.3 Early-Warning Signals

Figure 5 shows variation of early-warning signals with population size for heterozygote advantage. As population size decreases, the early-warning signals display three main phases in Case 2. The populations start with a phase where coefficient of variation, kurtosis and skewness remain constant. This is followed by a gradual change phase and lastly by a rapid change phase. Autocorrelation under heterozygote advantage also shows three phases but in different order (gradual increase, rapid decrease, constant). In Case 3 and the Control case, the populations exhibit either one phase where the indicator remains constant (coefficient of variation) or two phases where the constant phase is followed by a rapid change phase but characterised by large standard errors (kurtosis, skewness and autocorrelation). Gaussian smoothing before estimating the indicators gave similar observations (see Figure A.6 in the appendix). We deliberately leave out Case 1 because the populations rapidly declined from the beginning, making early-warning signals pointless, and most of them went extinct before the length of time window used. For fully density-dependent populations, similar observations are made (Figure A.11). Note though that compared to these aggregated results for populations in a given population-size range, time series of individual replicates are more noisy and may not display clearly distinct phases (see Figure A.5). Among the early-warning signals, the coefficient of variation seems to produce the most consistent patterns in individual replicates.

**Figure 5:**
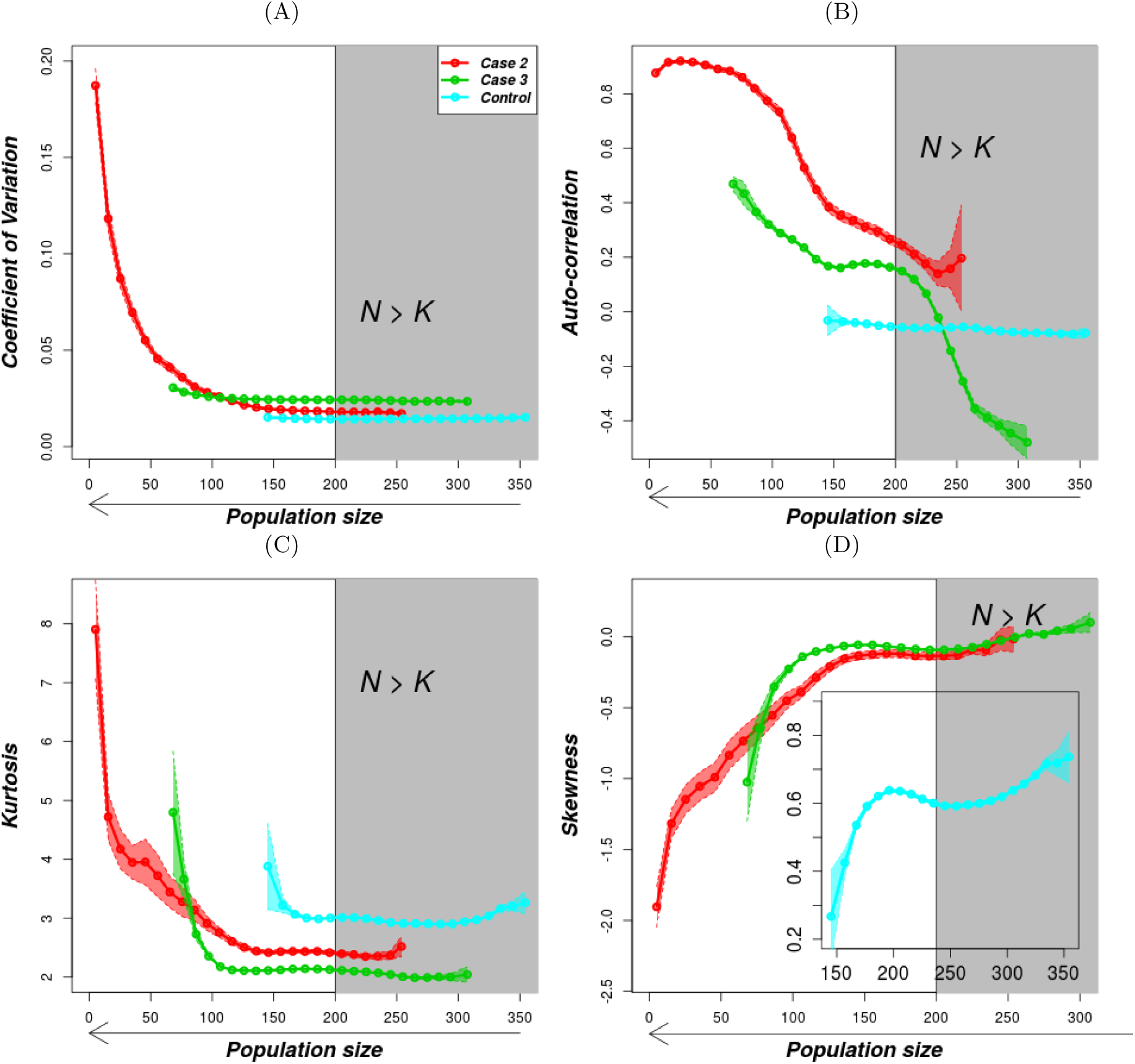
Variation of early-warning signals with population size under the geometric population growth model with heterozygote advantage. The gray-shaded part is the region above *K*. The solid lines are the mean values of the rates as described in the methods and the shaded regions shows the standard error while the gray-shaded rectangle is a region above *K*. The parameters are as in Figure 3. We leave out Case 1 because the populations rapidly decreased from the start and our time window range is above the time of population extinction. The arrows show direction of population size over time.

### 3.4 Fluctuating Selection with Reversal of Dominance

To check that the above observations are not restricted to the heterozygote advantage selection mechanism, we now compare the observations with another form of balancing selection known as fluctuating selection with reversal of dominance (see Figure 1). The dynamics of population size and number of polymorphic loci as time varies (Figure 6), and per-locus rate of loss and per-capita decline rate as population size and number of polymorphic loci decrease (see appendix, Figure A.12), show great similarity to those observed in the corresponding cases for heterozygote advantage. However, fluctuating patterns in population sizes in Cases 2 and 3 are more pronounced and reflect the patterns in the population mean fitness in Figure 1. In summary, only Cases 1 and 2 meet our definition of an eco-evolutionary vortex based on variation of per-capita decline rate and per-locus rate of loss. Also, in agreement with the proposed cues, this mechanism also shows existence of critical population size in Case 2 (since per-capita population decline changes from negative to positive as population size decreases). However, as opposed to heterozygote advantage, the number of polymorphic loci at which rapid decline to zero occurs varies greatly between trajectories (Figure 6E) with no critical number of polymorphic loci observed. The analytic approximation gives almost a linear trend with no sign of a critical number of polymorphic loci. The early-warning signals behaved as expected (Figure A.16), except that autocorrelation at lag 1 did not show a strong response. Case 3 populations do not meet our definition of an eco-evolutionary vortex nor show any of the suggested cues.

**Figure 6:**
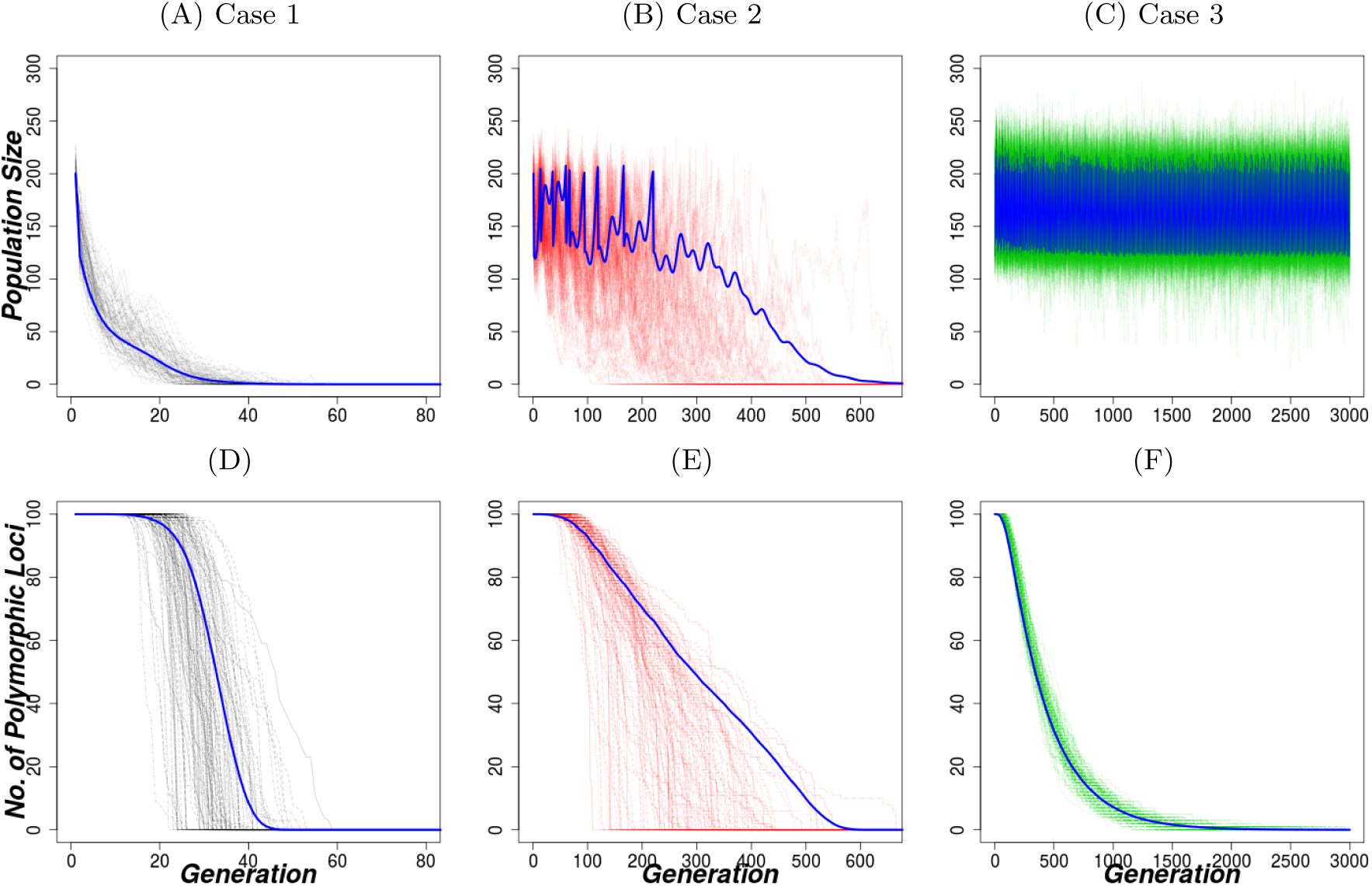
Dynamics of population size and genetic variation under the geometric population growth model with fluctuating selection and reversal of dominance. Each column represents one case indicated in Figure 2B. 1st column (Case 1): *r* = ln(1.4), 2nd column (Case 2): *r* = ln(1.616) and 3rd column (Case 3): *r* = ln(1.8). Other parameters are *s* = 0.005, *N*_0_ = *K* = 200, *replicates* = 100. The blue line is the corresponding analytic approximation with discretization parameters *u* = 0.1 and *τ* = 1. Notice the different scaling of the time-axis in the different columns.

For both selection mechanisms, additional cases in a given region show qualitatively similar results as their respective representative Cases 1, 2 and 3 (See Figures in Sections A.4.1 and A.6.1 in the appendix).

## 4 Discussion

We have developed a quantitative model for the feedback between loss of genetic variation and population decline. We identified parameter combinations that match our definition of an eco-evolutionary vortex and where populations exhibit the proposed cues of such extinction vortices. Our approach involved dividing parameter space into regions displaying qualitatively different behaviours as both population size and genetic variation decrease.

Parameter combinations in Regions I and II (see Figure 2) show clear interdependence between population size and genetic variation. They both agree with the proposed two defining features of an eco-evolutionary vortex, i.e. the per-capita rates of population decline and the per-locus rates of loss of polymorphic loci increase with both decreasing population size and decreasing number of polymorphic loci. Region III and the neutral Control did not match the defining features of an extinction vortex. Also, among the suggested cues, populations in Region I and II do not survive for long once total loss of genetic variation occurs (as seen from the trajectories). Further, only populations in Region II exhibits early-warning signals (see Section 4.1 below) and a critical number of polymorphic loci and population size below which a rapid population decline to extinction occurs. These critical values occur when the average number of offspring that survive per individual (*SOI*) falls below 1 (see also Section A.3 in the appendix). Note that in our models the threshold number of polymorphic loci is higher than the critical number calculated from *SOI* = 1 due to overestimation of fitness by assuming allele frequency of 0.5. Region III populations do well even after total loss of genetic variation and generally do not match our definition and suggested cues. However, for parameter combinations close to the boundary of Region II, the populations may not survive for long after (total) loss of polymorphic loci (see Figure A.8C), and also satisfy both suggested defining features of an eco-evolutionary vortex. This is because loss of polymorphic loci lowers the populations to very low sizes which exposes them to strong demographic stochasticity (Harmon and Braude 2010).

Furthermore, in Region II, the increasing per-capita decline rate as population size and polymorphic loci decrease can be seen as a genetic Allee effect. That is, we have a positive relationship between per-capita growth rate and population size at low population sizes (the hallmark of a demographic Allee effect), mediated by faster loss of genetic variation at small population sizes. Our definition of a genetic Allee effect roughly matches the two-step definition suggested by Luque et al. (2016), although in our case the first step is not a reduction in population size but loss of genetic variation. As drift pushes allele frequencies to the extremes, polymorphic loci start to be lost one by one. This leads to a decrease in individual fitness (decreased number of surviving offspring) and thereby population size. Wittmann et al. (2018) defined genetic Allee effects in terms of thresholds. A strong genetic Allee effect produces an inflection point in the persistence probability graph as a function of initial population size. In our study, the change in per-capita decline rate from negative to positive similarly suggests the existence of an eco-evolutionary Allee effect threshold.

The analytic approximation agrees with the simulations of the IBM in three ways: (1) The shape of individual replicates and analytic approximation is the same. (2) The approximation trajectory lies in the region of possible individual-based model trajectories. (3) The critical number of polymorphic loci is roughly the same, at least under heterozygote advantage. Thus for many purposes it may be possible to use the analytic model, which is computationally less demanding than the individual-based model, as a short-cut to derive results on eco-evolutionary extinction vortices.

### 4.1 Early-warning signals

The aim of checking for early-warning signals in this study was to find out whether signals can be observed before a population enters an extinction vortex. All the indicators behaved as expected for declining populations (Sommer et al. 2017; Dakos et al. 2012). As population size decreases, the populations show generally three phases for cases where reduction in genetic variation results into population extinction (e.g. Region II). That is, an initial constant phase followed by a gradual (linear) change phase and finally a rapid change phase can be observed. Again with regards to our suggested cue that decreasing genetic variation produces early-warning signals in population dynamics, the start of the gradual change phase signals the entry into a vortex while the start of the rapid phase signals nearby extinction. For Region 3, populations generally show one or two phases. The constant phase and either a gradual phase or a rapid phase. But with a rapid change phase, there are relatively large standard errors. It should be noted that decisions should not depend on a single early-warning signal but a combination of them, as results from indicators may not always agree with each other (Gsell et al. 2016). Also, some signals were relatively noisy in individual replicates or only appeared when the population was already very low. As population size decreases, we find changes in signals from coefficient of variation/standard deviation, skewness and kurtosis consistent with each other and also with the changes in per-capita decline rates. However, autocorrelation shows inconsistency with other measured indicators.

### 4.2 Generality and genetic realism

Small populations whose persistence is mainly driven by genetic factors are usually thought to be far below their environmental carrying capacity and little affected by negative density dependence (Lynch and Gabriel 1990; Kim et al. 2016; Wang et al. 2019). Here, we used the geometric model with a Ricker boundary to mimic such populations. To check whether our observations are not an artifact of this assumption, we also used a Ricker model to study fully density-dependent populations. We observe comparable features as shown in Figures A.7 and A.8. However, a higher *K* is required to observe rapid population decline in Region II (Figure A.8). The high *K* allows for sufficiently weak negative density-dependence so that the positive feedback induced by the loss of genetic variation can dominate and cause a rapid decline to extinction (see Appendix section A.5 for analogous figures and more discussion).

The variation of per-locus rates of loss of polymorphic loci with population size and polymorphic loci agree with observations in extinction vortices driven by accumulation of deleterious mutations. Models that consider (deleterious) mutation accumulation show that the rate at which new mutations get fixed increases with increasing number of such mutations already fixed (Coron et al. 2013). Also, Lynch and Gabriel (1990) observed that an extinction vortex due to mutation accumulation leads to a small coefficient of variation in extinction times. In our model, the coefficient of variation for population extinction time for regions with extinction vortex is also substantially below 1 (1 being the expectation with exponentially distributed extinction times). For fluctuating selection, the coefficients of variation of extinction times for the two cases 1 and 2 are 0.22 and 0.35 while for heterozygote advantage, the coefficients are 0.22 and 0.13 for Cases 1 and 2 respectively.

Various forms of balancing selection such as negative frequency-dependent selection, heterozygote advantage and some types of fluctuating selection contribute to maintaining genetic variation in populations. In our study, both mechanisms for the maintenance of genetic variation are based on heterozygote advantage directly or indirectly in the long run. These mechanisms create a positive relationship between the number of polymorphic loci and fitness. Other mechanisms that have a similar relationship might produce similar eco-evolutionary vortices. Future models with mechanisms that are not based on heterozygote advantage such as frequency-dependent selection or more generally evolutionary potential in a variable environment are necessary to check the generality of the above eco-evolutionary vortex.

The model presented here assumes that polymorphism that is lost is lost forever. This is a good approximation if the time scale of extinction is short relative to the time scale at which mutations occur. However, to study if there is a critical population size above which populations are safe from an eco-evolutionary vortex, future work will need to include back-mutations or compensatory mutations.

## Supporting information

R scripts for regenerating data used in the manuscript

## 4.3 Acknowledgements

We thank the Theoretical Biology group, Bielefeld University for the discussions and generously sharing insights especially Koen van Benthem on early-warning signals.

## Appendices

### A Supplementary Information

#### A.1 Diffusion Approximation of Allele-Frequency Distribution

The probability density function, *f*(*x,t*) of allele frequency *x* at time *t* can be approximated using a forward Kolmogorov equation

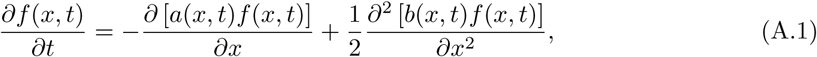

where *a*(*x, t*) and *b*(*x, t*) are the infinitesimal mean and infinitesimal variance of allele frequency *x*, at time *t* (Kimura et al. 1955). Subject to the initial and boundary conditions, Equation (A.1) can be solved numerically and in some cases analytically.

In the case of heterozygote advantage with both homozygotes having fitness of 1 − *s*, where *s* > 0 and heterozygotes having fitness 1, then, it can be shown that

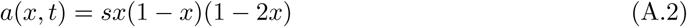

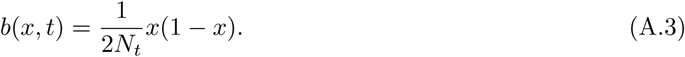

In this study, we solve the diffusion equation numerically adopting the scheme in Zhao et al. (2013) and Xu et al. (2019) without rescaling time. The numerical method gives a complete solution of the problem, i.e., it keeps the law of total probability at all times. The solution is based on Finite Volume Methods (FVM) using a central scheme (function values approximated in the middle of a set of grid cells). We briefly describe the main steps followed here.

The boundary condition stems from conservation of total probability at every time *t*. We assume zero flux density at the boundaries, i.e., no probability flow outside the system as the probability of fixation or loss is captured by the outermost bins. To see this, we first rewrite Equation (A.1) in the form

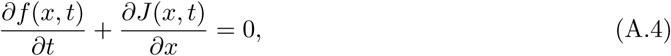

where

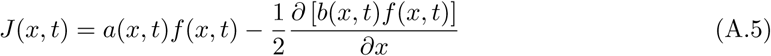

represents a probability flow. Thus the boundary condition requires that

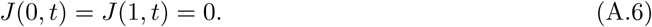

The initial condition is obtained by assuming the populations start at a single allele frequency *x*_0_. Thus *f*(*x*, 0) = *x*_0_, represented by a Dirac function whose mass is concentrated at *x* = *x*_0_.

We discretize *x* into a grid with spacing *u* = 1*/m*. That is *x*_*i*_ = *i* · *u, i* = 0, 1, …, *m*. Also, time is discretized with spacing *τ* and the grid points are *t*_*v*_ = *v* · *τ, v* = 0, 1, … The control volume *D*_*i*_, for the inner mesh points *x*_*i*_ is given by

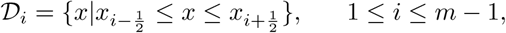

where 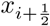 is the grid mid-point between *x*_*i*_ and *x*_*i*+1_.

Equation (A.4) can then be approximated using FVM by

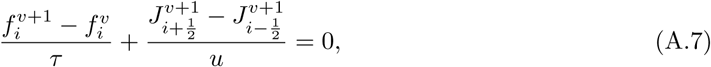

where *J*_•_ is approximated using a central scheme as

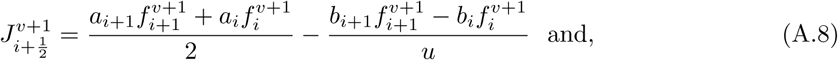

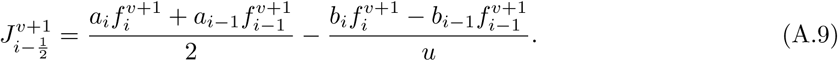

Substituting (A.8) and (A.9) for 1 ≤ *i* ≤ *m* − 1, into Equation (A.7) and after some algebra, we get

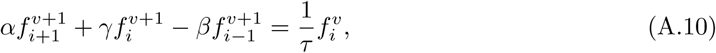

where

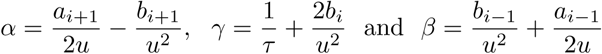

At the boundaries, the mesh points *x*_0_ = 0 and *x*_*m*_ = 1 have the control volume

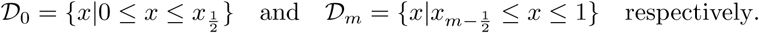

Now, using the boundary condition given in (A.6),

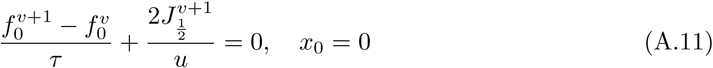

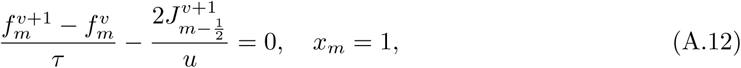

which after some rearrangement yields

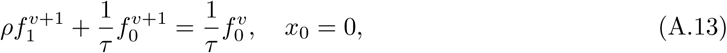

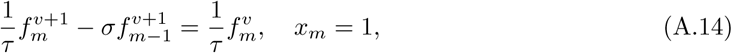

where

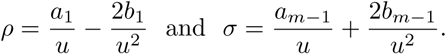

Combining Equations (A.10), (A.13) and (A.14) yields a system of difference equations generally expressed in matrix form as

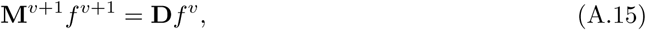

where **D** is a (*m* + 1) × (*m* + 1) diagonal matrix whose leading diagonal is filled with 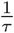 and **M** is also a (*m* + 1) × (*m* + 1) tridiagonal matrix filled as follows. For *i* = 0 and *i* = *m*

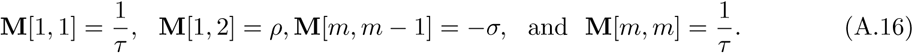

For 1 ≤ *i* ≤ *m* − 1,

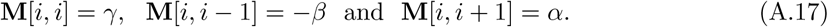

For a detailed discussion of this numerical scheme, see Zhao et al. (2013) and Xu et al. (2019). The density of allele frequencies at time-step *v* + 1 given the density at time-step *v* is

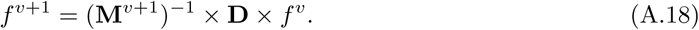

The densities of allele frequencies for some of the scenarios considered in the main text are shown in Figure A.1.

**Figure A.1:**
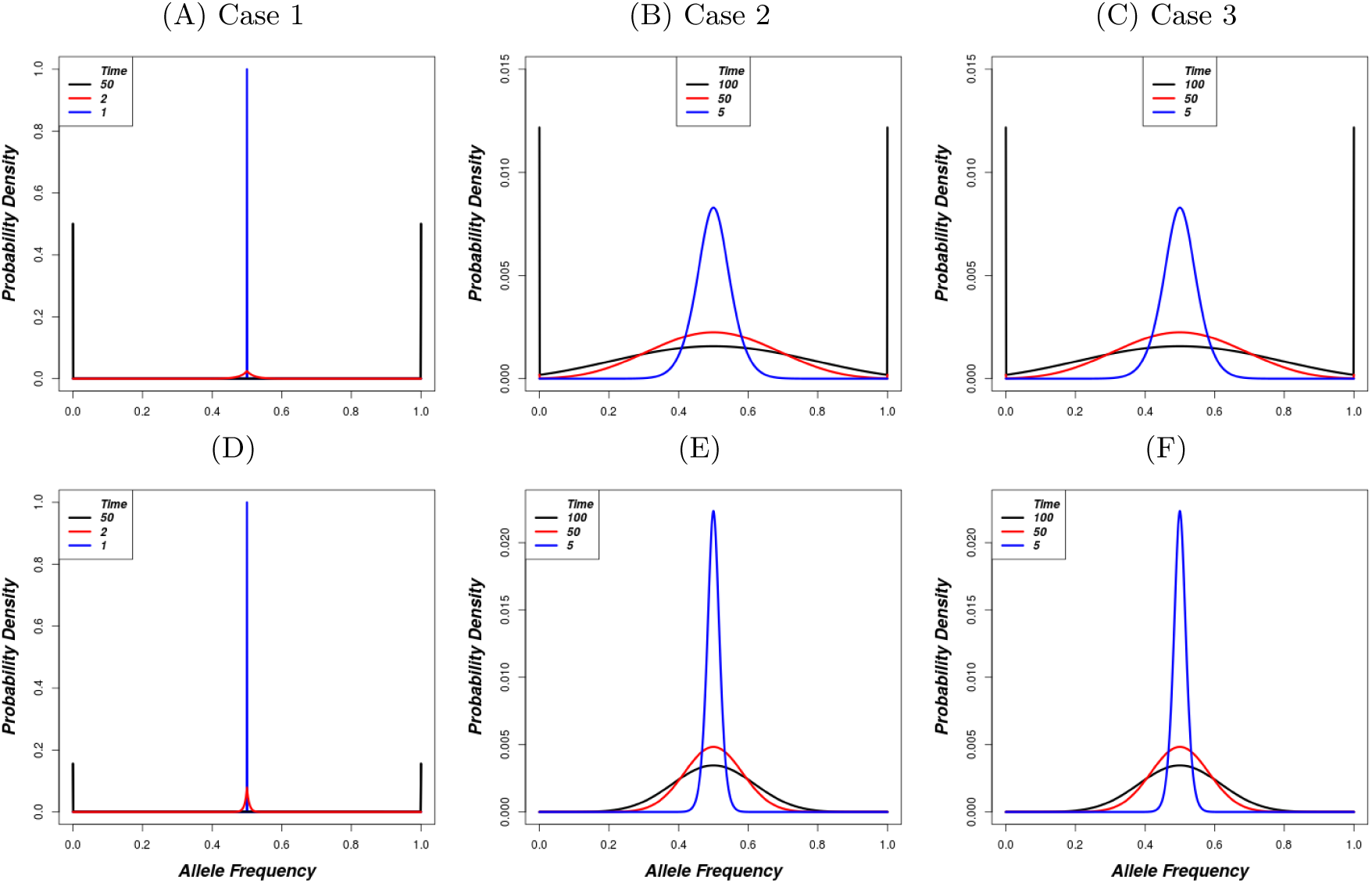
Probability density of allele frequencies *f*(*x, t*) at different times for heterozygous advantage. First row: Geometric model with *N*_0_ = *K* = 200. 2nd row: Ricker model with *N*_0_ = *K* = 2000. Other parameters are the same for both rows i.e. *s* = 0.005, *x*_0_ = 0.5, *m* = 1000, *τ* = 1.

#### A.2 Infinitesimal Mean for Fluctuating Selection

Consider the three genotypes *AA, Aa* and *aa* with fitness 1 − *s*_*A,t*_, 1 − *h*_*t*_*s*_*A,t*_ − (1 − *h*_*t*_)*s*_*a,t*_ and 1 − *s*_*a,t*_ at time *t*, respectively, where *h*_*t*_, *s*_*a,t*_ and *s*_*A,t*_ are as described in the main text (Section 2). Also, we assume the frequency of allele *A* is *x*_*t*_ at time *t*. For convenience, we drop subscript *t* in the allele frequency and in the selection and dominance coefficients. After the action of selection, the allele frequency in the next generation *x*′ is

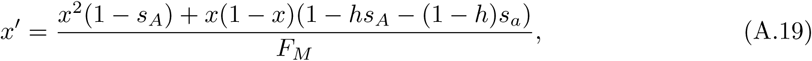

where *F*_*M*_ is the mean fitness given by

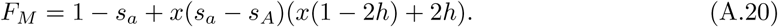

After some algebra, the change in allele frequency *a*(*x, t*) = *x*′ − *x* is

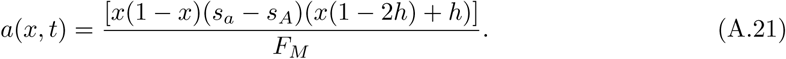

Using a Taylor series expansion

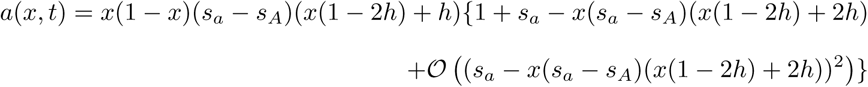

Assuming the coefficients *s*_*a*_ and *s*_*A*_ are small and ignoring terms with 3rd order and above in the expansion yields

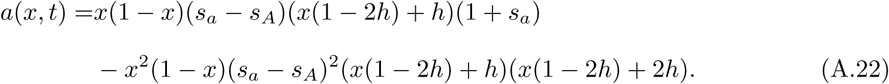

#### A.3 Number of Offspring that Survive per Parent (SOI)

The number of offspring that survive are given by

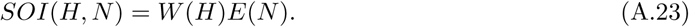

Differentiating *SOI* with respect to *H*,

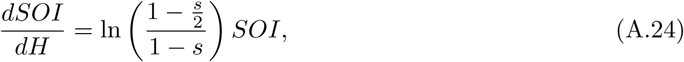

shows that the number of offspring that survives increases with *H* and the second derivative of *SOI*

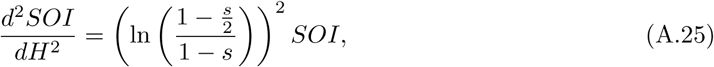

is positive. This implies that *SOI* increases even faster as *H* increases.

Simulations show that the decline in population size and loss of polymorphic loci becomes faster when the number surviving offspring per individual is just less than 1. Assuming *N* << *K* and therefore 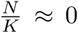 0 so that we can assume that *E*(*N*) ≈ *e*^*r*^, we obtain *H*_*c*_, the critical number of polymorphic loci for *SOI* = 1 as

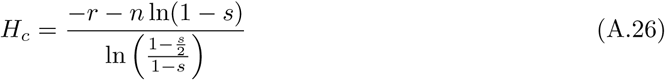

If the ratio of population size and carrying capacity is not small enough to be negligible, then *E*_*t*_ follows the pure Ricker model dynamics, making *SOI* a function of *N*. Differentiating *SOI* with respect to *N* gives

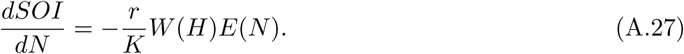

Equation (A.27) shows that *SOI* increases as *N* decreases at constant *H*. The second derivative with respect to *N* is positive implying that *SOI* increases even faster as *N* decreases. As earlier seen in Equations (A.24) and (A.25), *SOI* decreases faster with decrease in *H*. So, the net direction of *SOI* depends on how fast *SOI* decreases with decreasing *H* and increases with decreasing *N*. Small values of *r* combined with higher values of *s* would result into net decrease in *SOI* and when below 1, the population accelerates to extinction.

#### A.4 Supplementary figures on the geometric growth model with heterozygote advantage

##### A.4.1 Other Randomly Chosen Points in Regions I and II

**Figure A.2:**
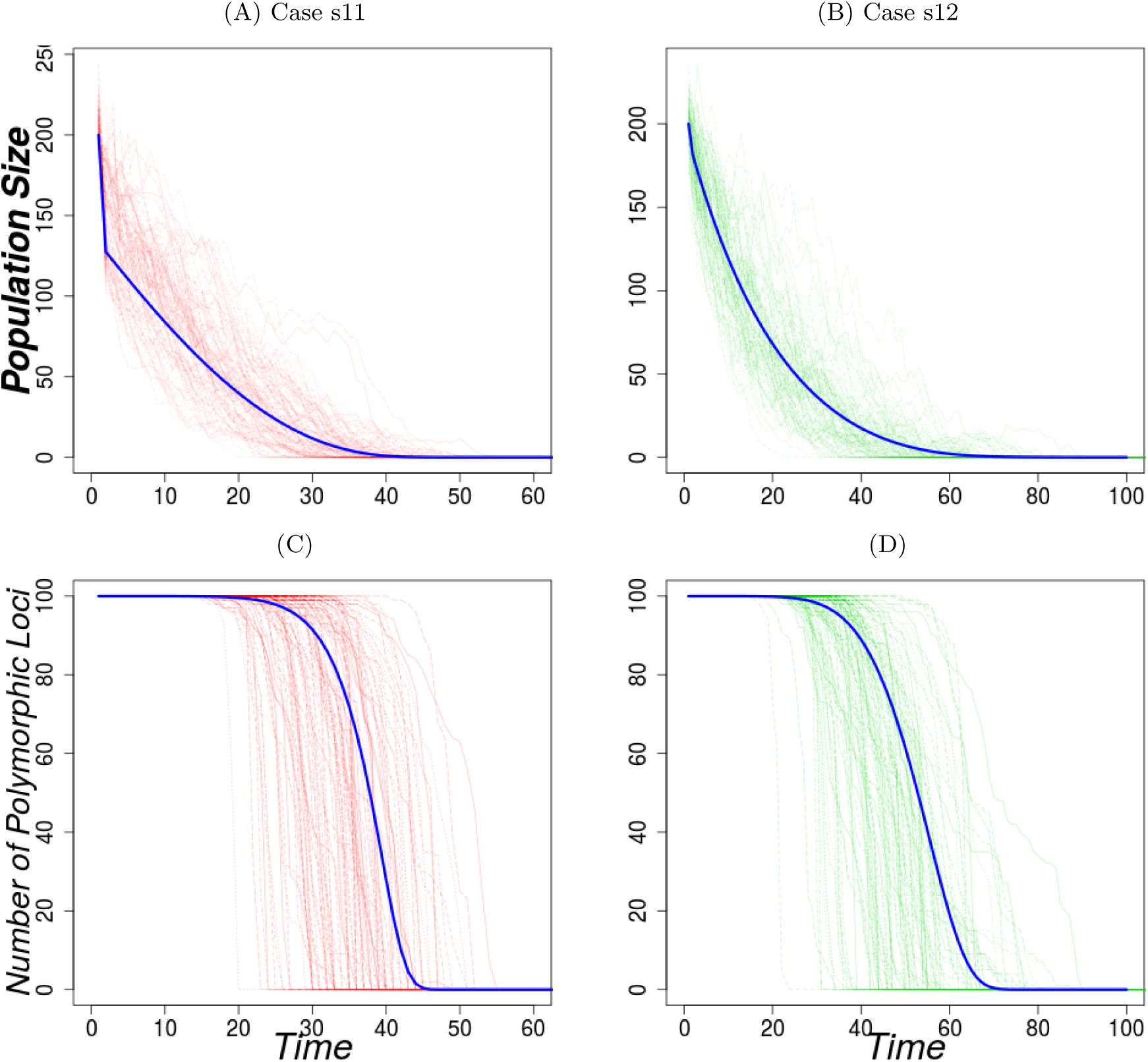
Variation of the population size and the number of polymorphic loci with time under the geometric growth model with heterozygote advantage for Region I supplementary points. The dark blue lines show the analytic approximation with discretization parameters *u* = 0.1 and *τ* = 1. The parameters used for the 1st column (Case s11) are *r* = 1.50, *s* = 0.009 and for the 2nd column (Case s12) *r* = 1.05, *s* = 0.002. Other parameters are *N*_0_ = *K* = 200, *replicates* = 100

**Figure A.3:**
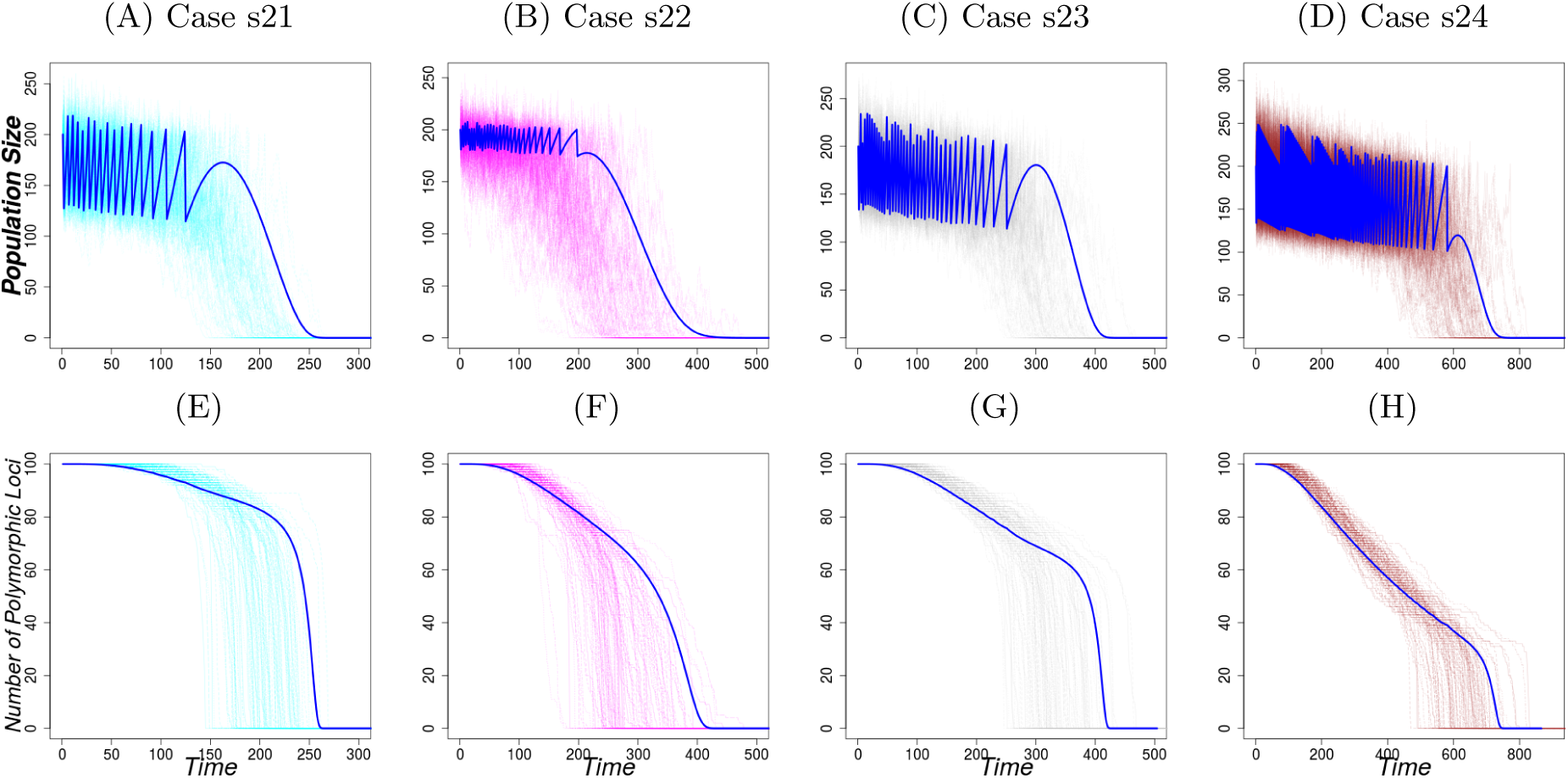
Variation of the population size and the number of polymorphic loci with time under the geometric growth model with heterozygote advantage for Region II supplementary points. The dark blue lines show the analytic approximation with discretization parameters *u* = 0.1 and *τ* = 1. The parameters used for 1st column (Case s21) *r* = 1.80, *s* = 0.009, 2nd column (Case s22) *r* = 1.15, *s* = 0.002, 3rd column (Case s23) *r* = 1.8, *s* = 0.008 and 4th column (Case s24) *r* = 2.00, *s* = 0.008. Other parameters are *N*_0_ = *K* = 200, *replicates* = 100

**Figure A.4:**
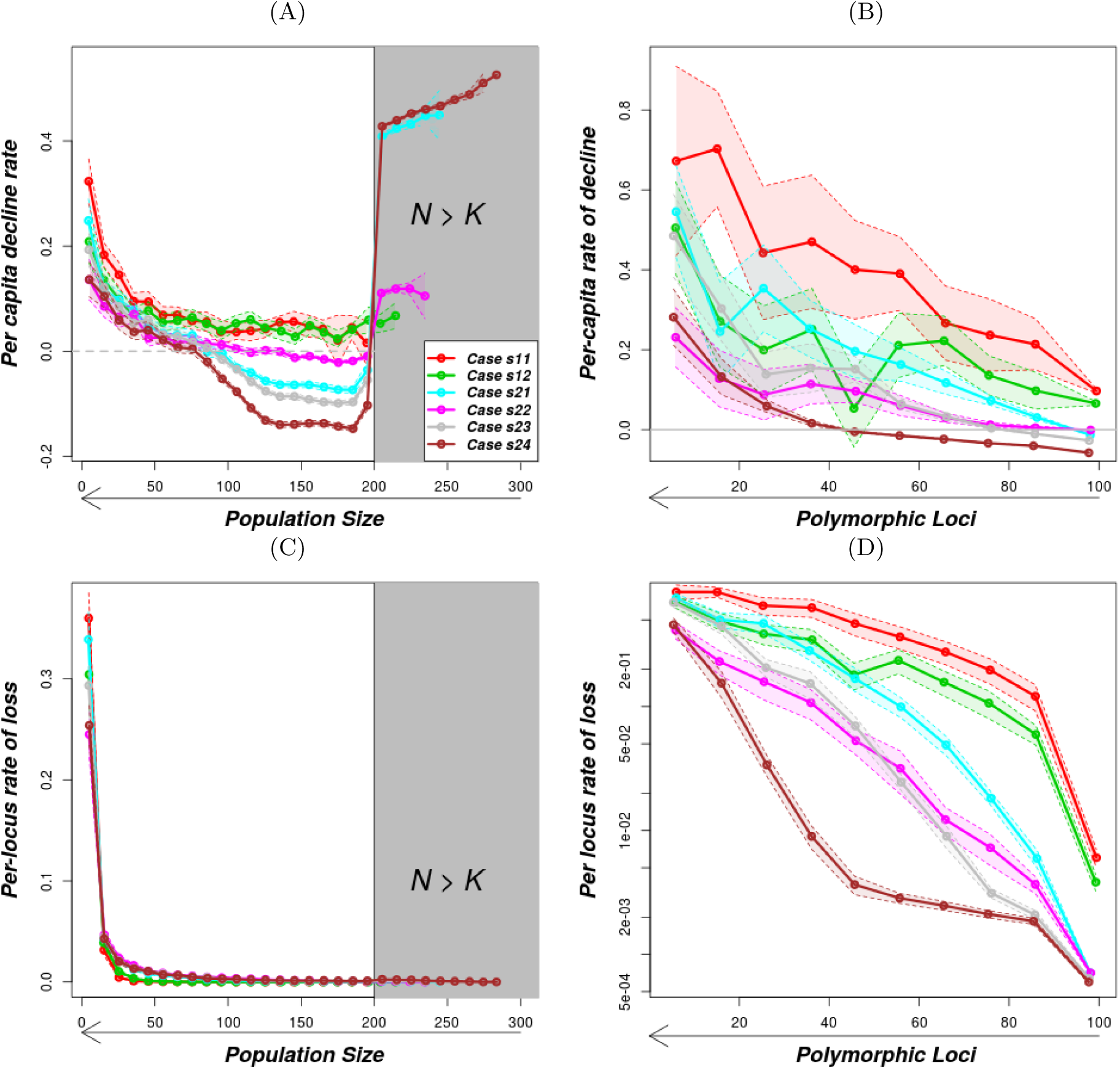
Variation of the per-capita decline rate and per-locus rate of loss with population size and the number of polymorphic loci under the geometric growth model with heterozygote advantage for Region I and II supplementary points. The parameters used are as on Figures A.2 and A.3 for the respective cases.

##### A.4.2 Early-warning signals

**Figure A.5:**
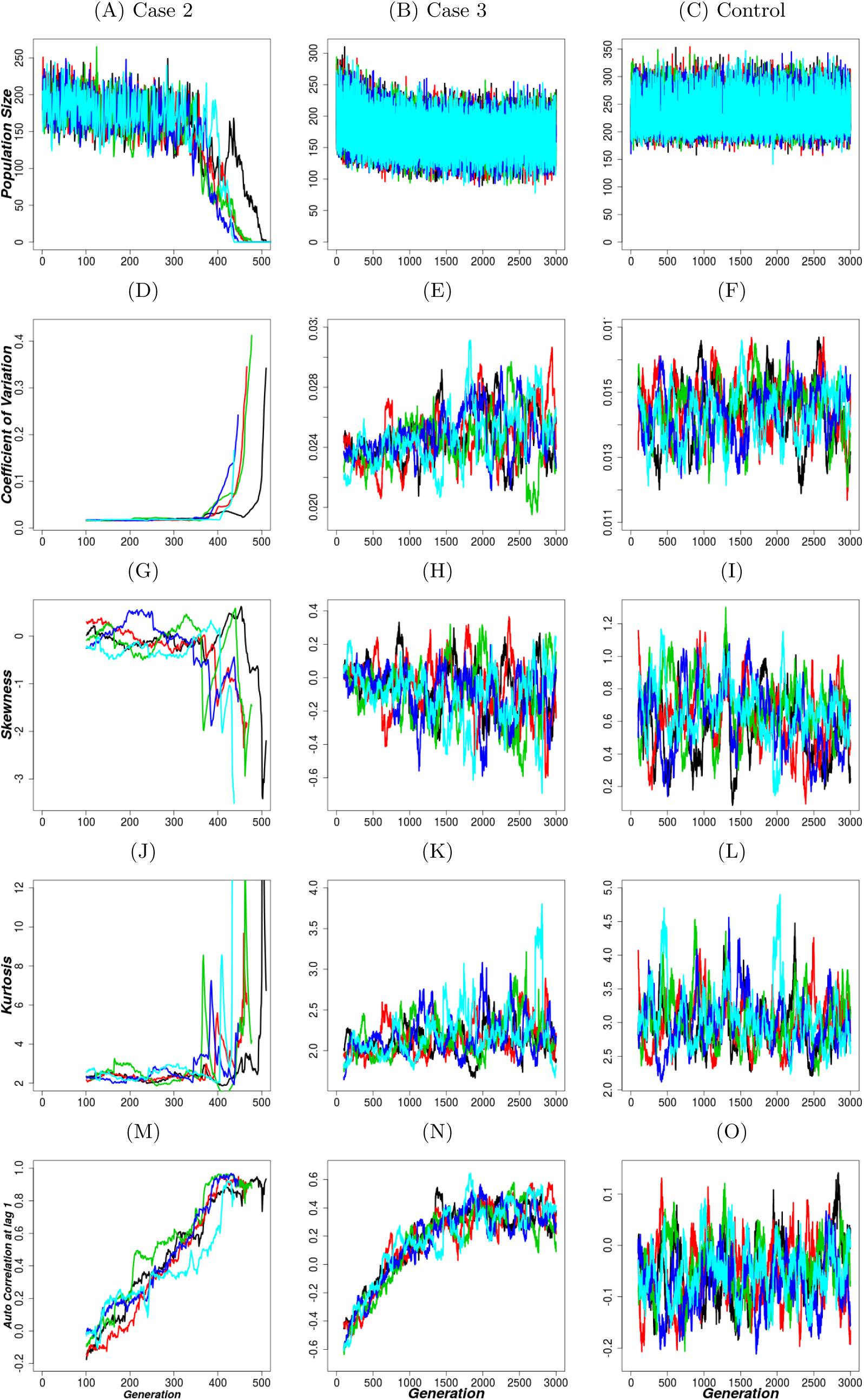
A random sample of 5 IBM realisations showing population-size trajectories (1st row) and the corresponding trajectories of early-warning signals (2nd - 5th row) under the geometric growth model with heterozygote advantage. The indicators start from the the 100th generation (because the window range is 100 here). The parameters for the cases are the same as those in Figure 3.

**Figure A.6:**
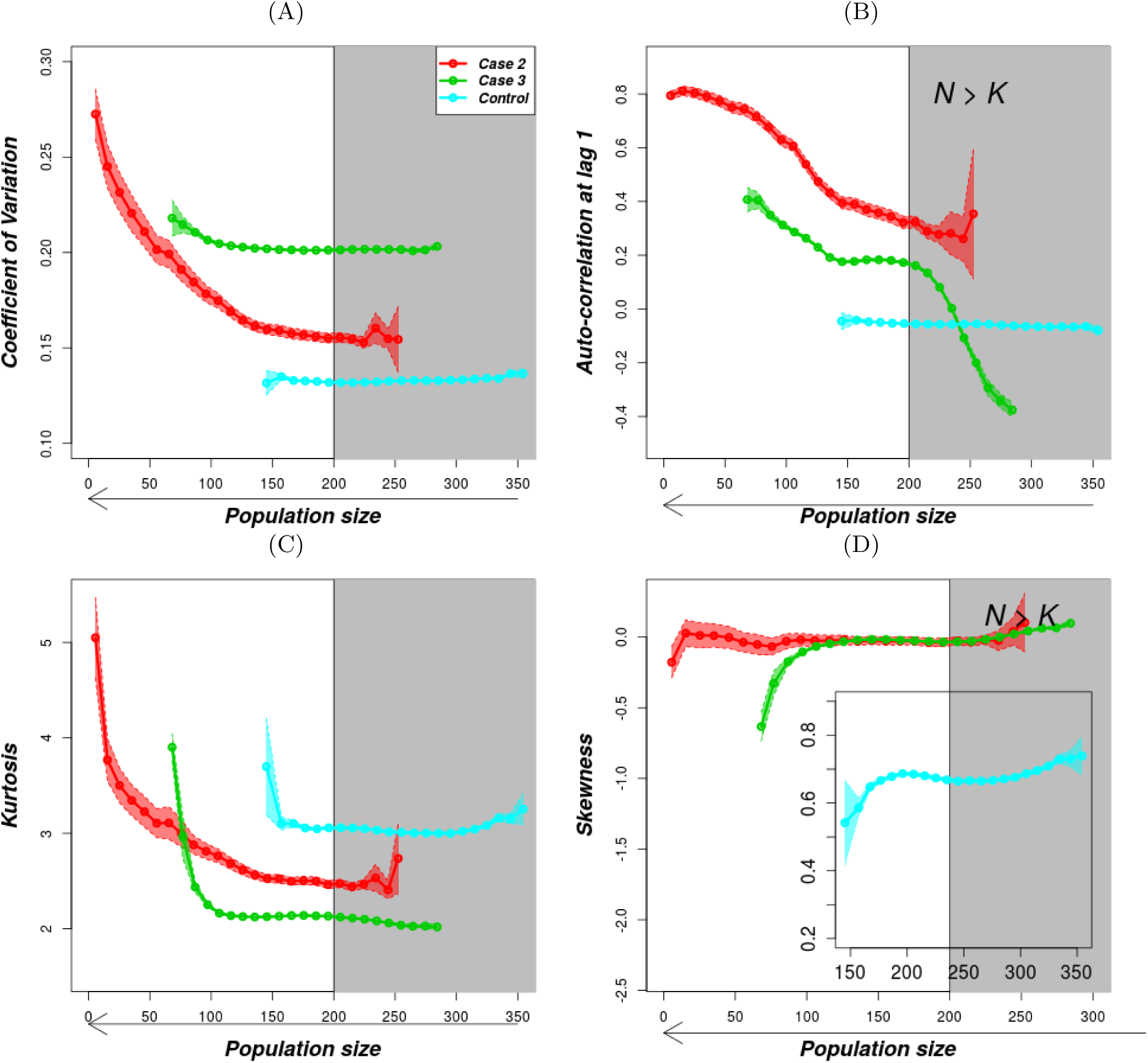
Variation of early-warning indicators with population size after Gaussian fitting under the geometric population growth model with heterozygote advantage. The solid lines are the mean values of the rates as described in the methods and the shaded regions shows the standard error while the gray-shaded rectangle is a region above carrying capacity. The parameters are as in Figure 3.

#### A.5 Supplementary materials for the Ricker model with heterozygote advantage

To produce a rapid decline in population size, there is need for a large carrying capacity. The reason is that the overall population growth rate (product of fitness and fertility) increases exponentially with decreasing population size. The realized population growth rate is given by 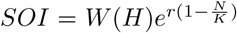. The equilibrium population size is below *K* even when we have 100% polymorphic loci because the mean fitness *W*(*H*) is always below 1.

As genetic variation decreases, the population fitness decreases, while as population size decreases, fertility increases. For a small carrying capacity, fertility may increase faster than the decrease in fitness which results into *SOI* > 1. No rapid population decline to extinction may be observed in this case. Nevertheless, both small and large carrying capacity populations still exhibit similar trends in the per-capita and per-locus rates as population size decreases (see Figures A.9 and A.10). This implies that some small populations may face an extinction vortex unnoticed if the main focus is on how rapid the population size declines.

**Figure A.7:**
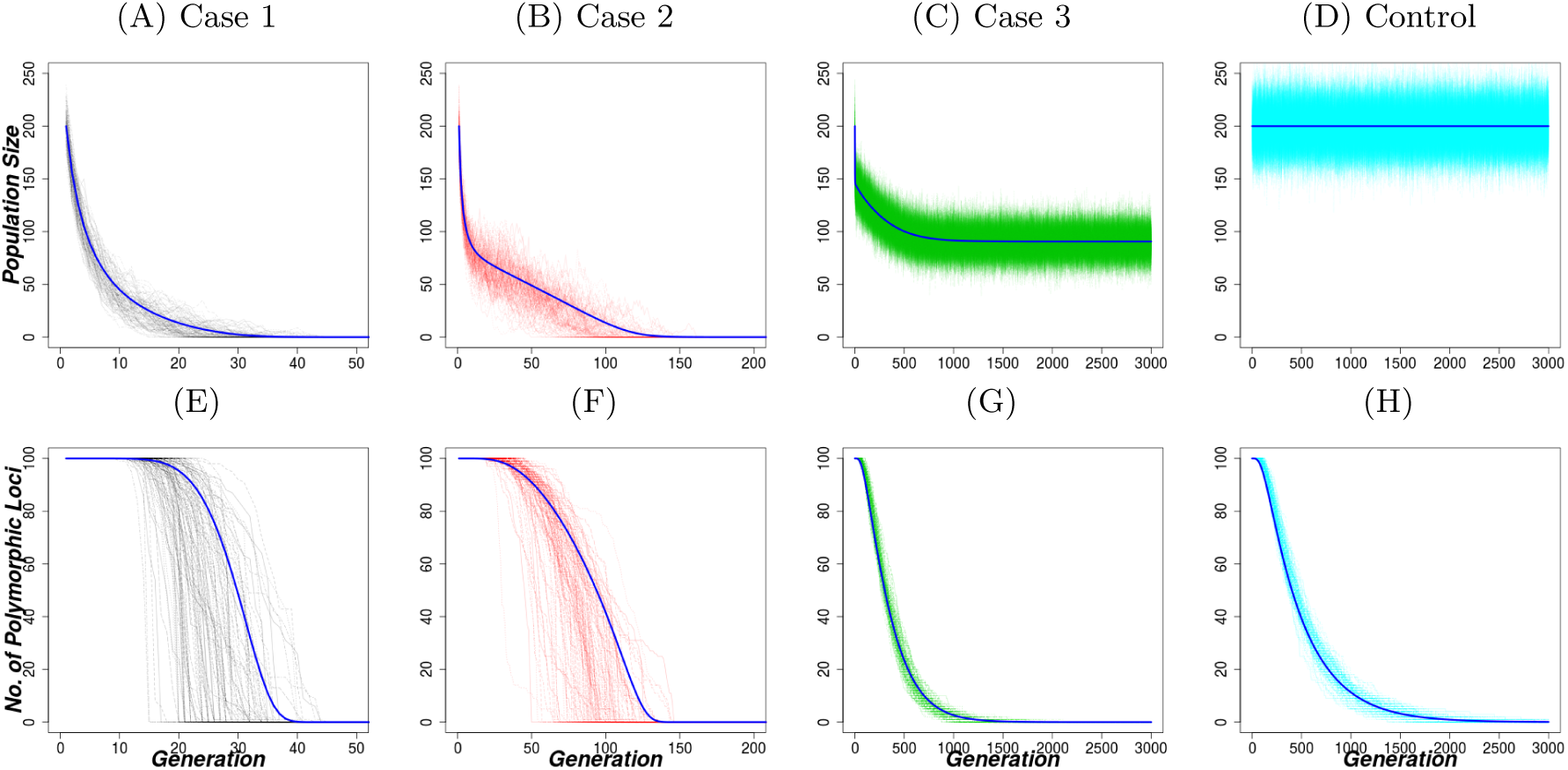
Dynamics of population size and genetic variation under the Ricker population growth model with heterozygote advantage. Each column represents one case indicated in Figure 2A. 1st column (Case 1): *r* = ln(1.2), 2nd column (Case 2): *r* = ln(1.5) and 3rd column (Case 3): *r* = ln(2.5). Other parameters are *s* = 0.005, *N*_0_ = *K* = 200, *replicates* = 100. The 4th column represents the control case with the same parameters as in the 2nd column but *s* = 0.00. The blue line is the corresponding analytic approximation with discretization parameters *u* = 0.1 and *τ* = 1.

**Figure A.8:**
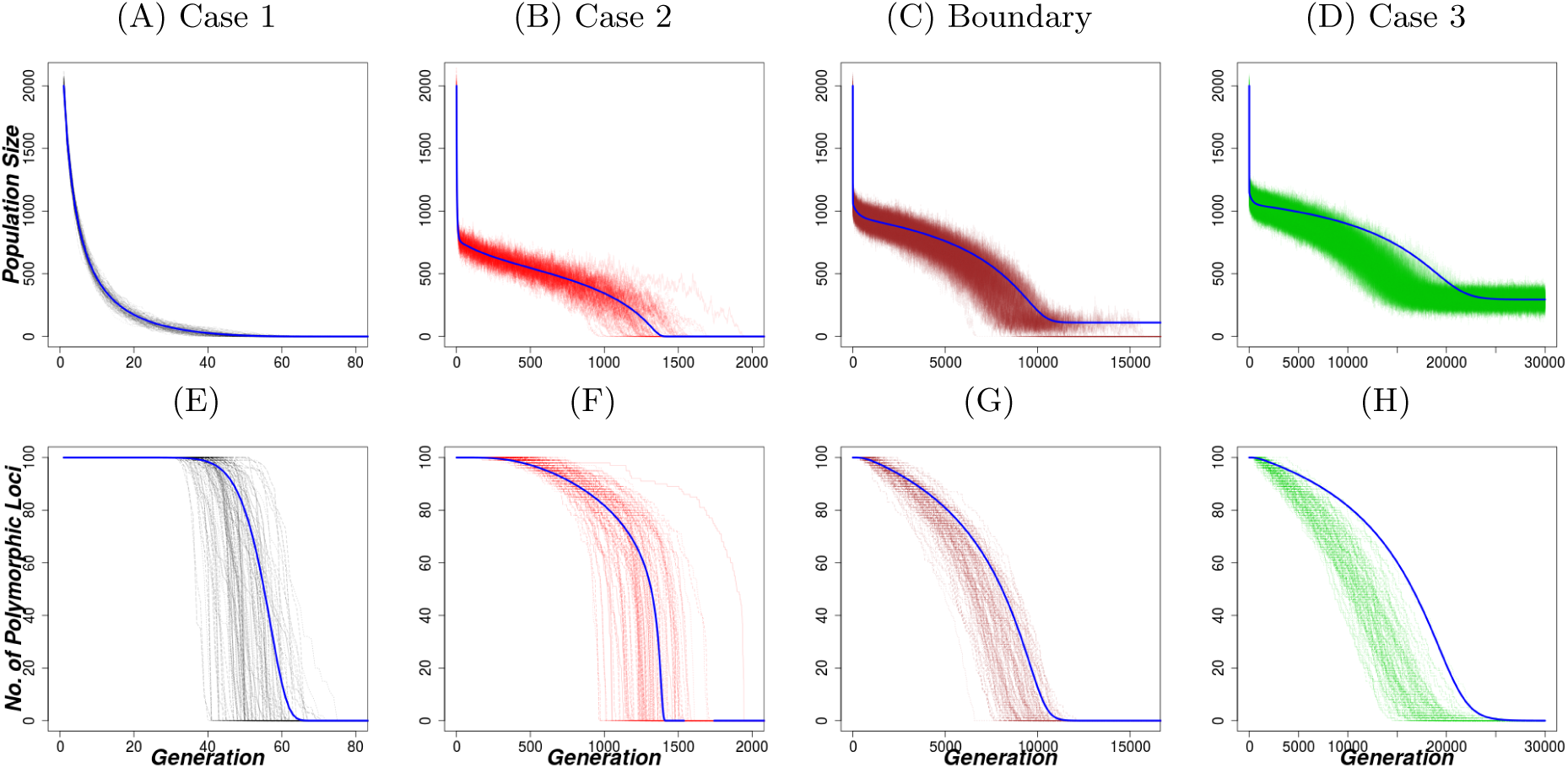
Dynamics of population size and genetic variation under the Ricker population growth model with heterozygote advantage and an increased carrying capacity. Each column represents one case indicated in Figure 2A. 1st column (Case 1): *r* = ln(1.2), 2nd column (Case 2): *r* = ln(1.5) and 3rd column (Case 3): *r* = ln(1.8). Other parameters are *s* = 0.005, *N*_0_ = *K* = 2000, *replicates* = 100. The 4th column represents the boundary case with the same parameters as for the other cases but *r* = ln(1.7). The blue line is the corresponding analytic approximation with discretization parameters *u* = 0.1 and *τ* = 1.

**Figure A.9:**
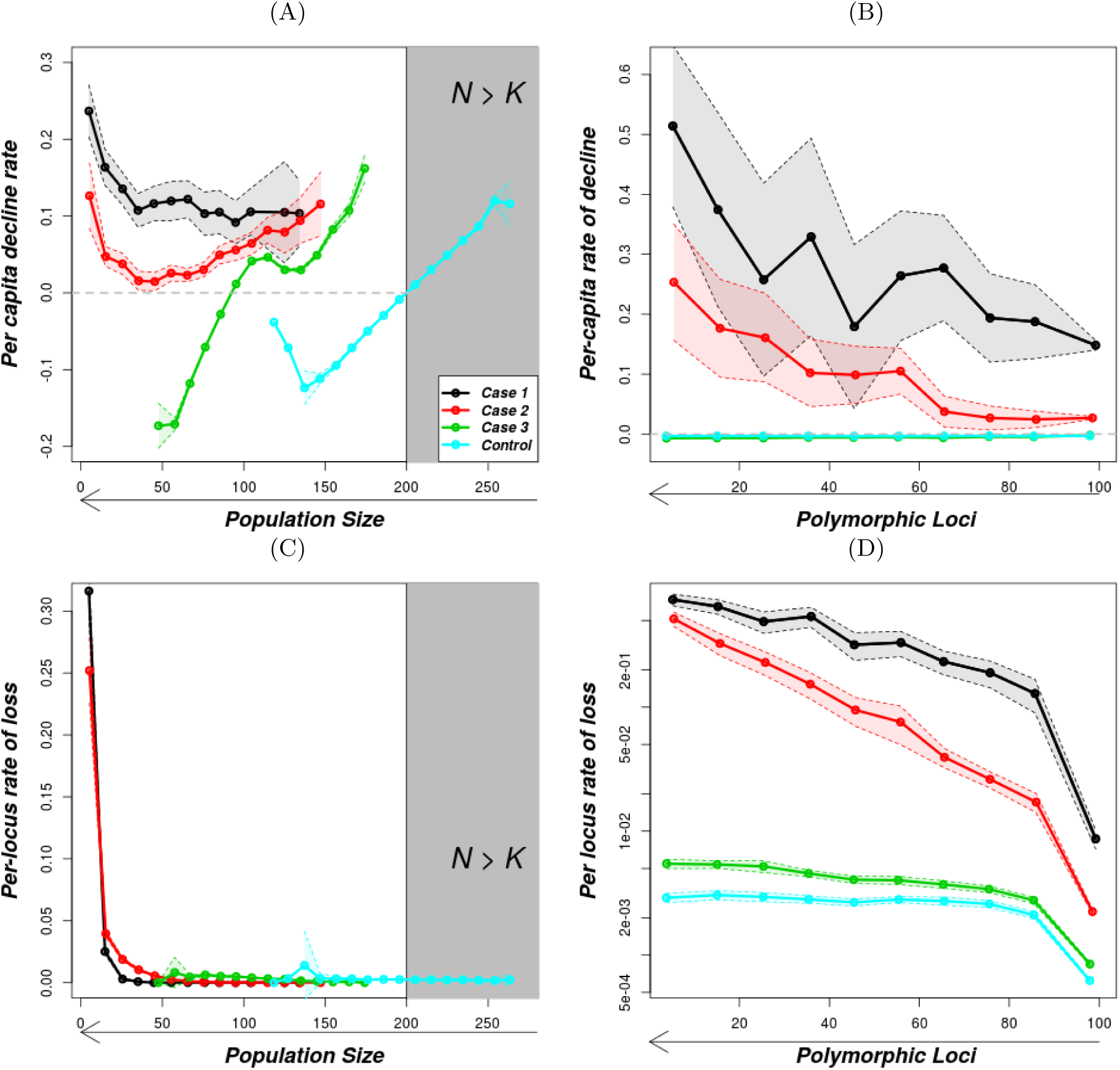
Per-capita rates of population decline and per-locus rates of loss of polymorphic loci under the Ricker population growth model with heterozygote advantage. The solid lines are the mean values of the rates as described in the methods and the shaded regions shows the standard error. The parameters for the 4 cases are as in Figure A.7. In Figure (d), the y-axis is on a logarithmic scale.

**Figure A.10:**
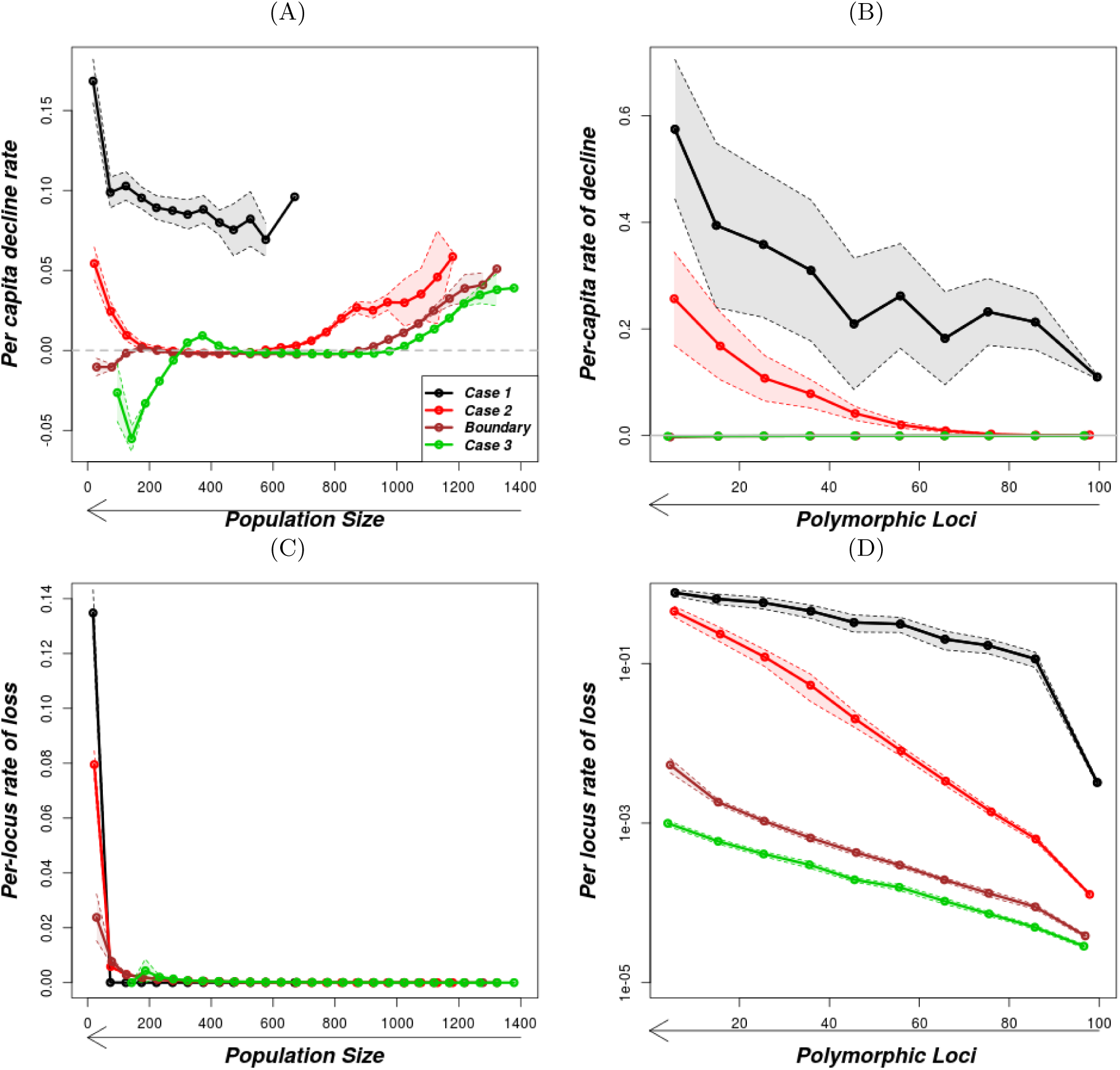
Per-capita rates of population decline and per-locus rates of loss of polymorphic loci under the Ricker population growth model with heterozygote advantage. The solid lines are the mean values of the rates as described in the methods and the shaded regions shows the standard error. The parameters for the 4 cases are as in Figure A.8. In (d), the y-axis is on a logarithmic scale.

**Figure A.11:**
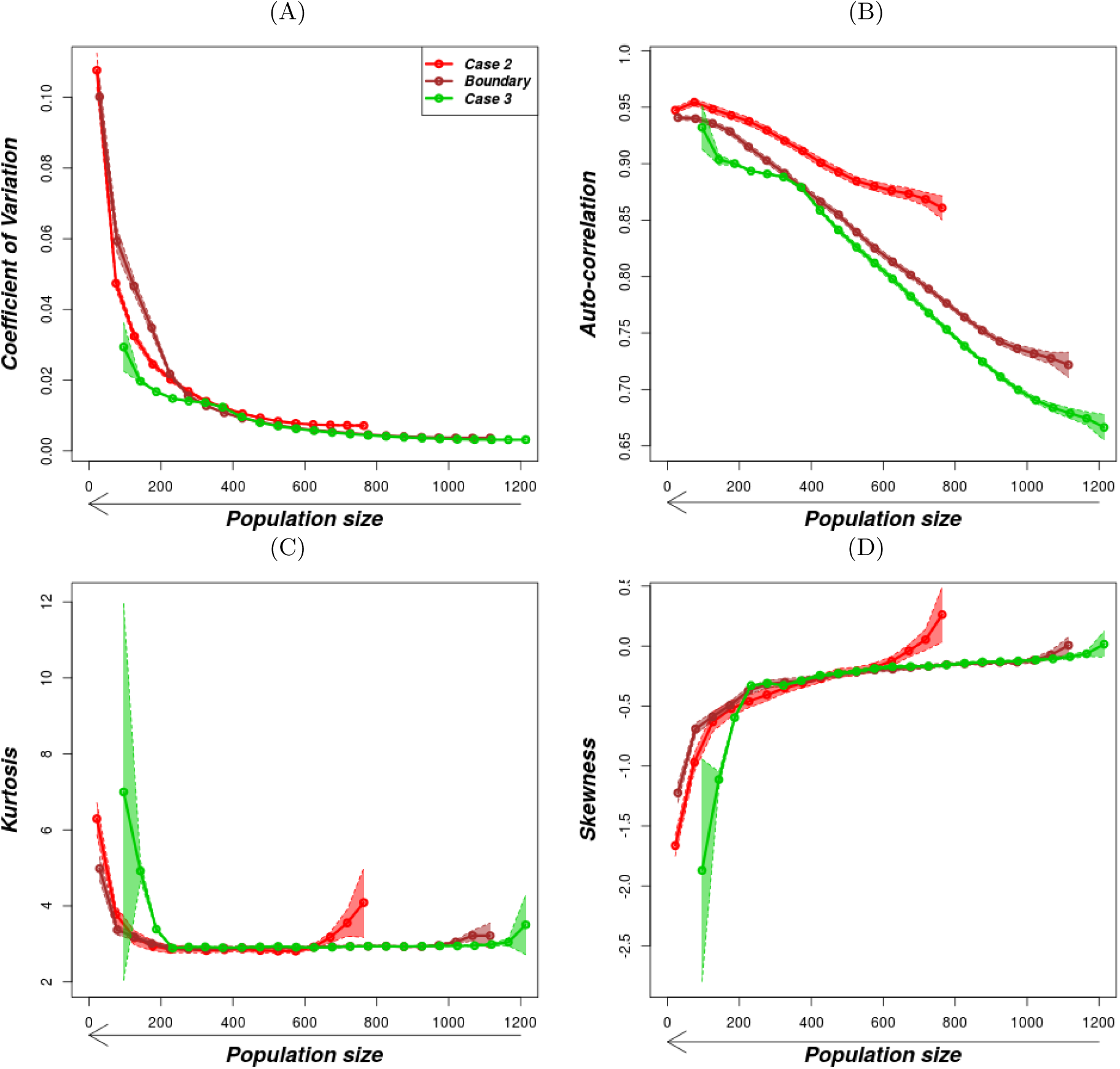
Variation of early-warning indicators with population size under the Ricker population growth model with heterozygote advantage. The solid lines are the mean values of the rates as described in the methods and the shaded regions shows the standard error while the gray-shaded rectangle is a region above carrying capacity. The parameters are as in Figure A.8.

#### A.6 Supplementary figures on the geometric growth model with fluctuating selection and dominance reversal

**Figure A.12:**
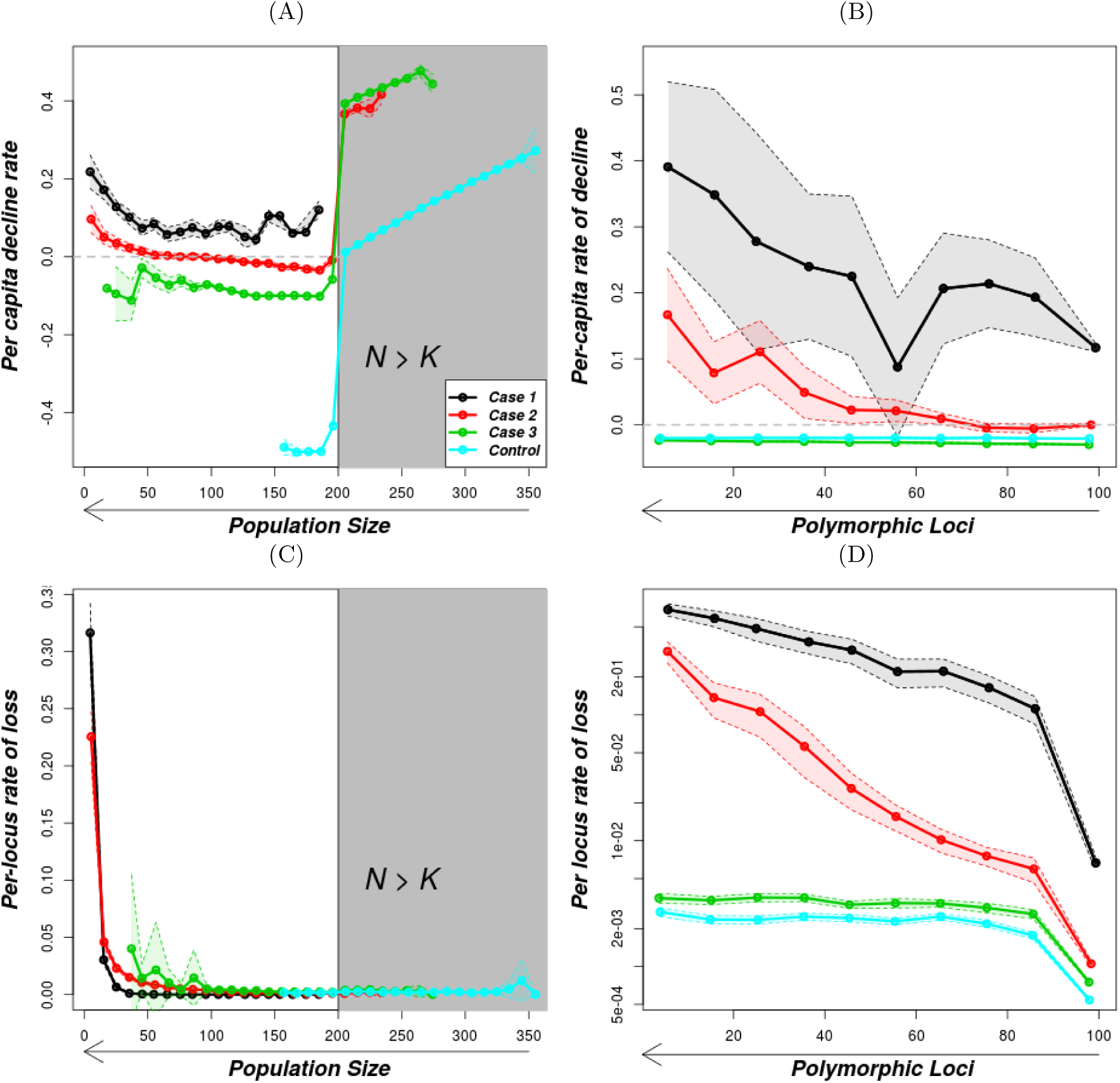
Variation of per-capita decline rate and per-locus rate of loss as population size and number of polymorphic loci decreases under the geometric population growth model and fluctuating selection with reversal of dominance. The gray-shaded part in (a) is the region above carrying capacity. The solid lines are the mean values of the rates as described in the methods and the shaded regions shows the standard error. The horizontal gray line in (a) and (c) is for rate 0. The parameters for the 4 cases are as in Figure 6 and for the control case, the parameters are the same as those in Figure 3 column 4.

##### A.6.1 Other randomly chosen points in Regions I and II

**Figure A.13:**
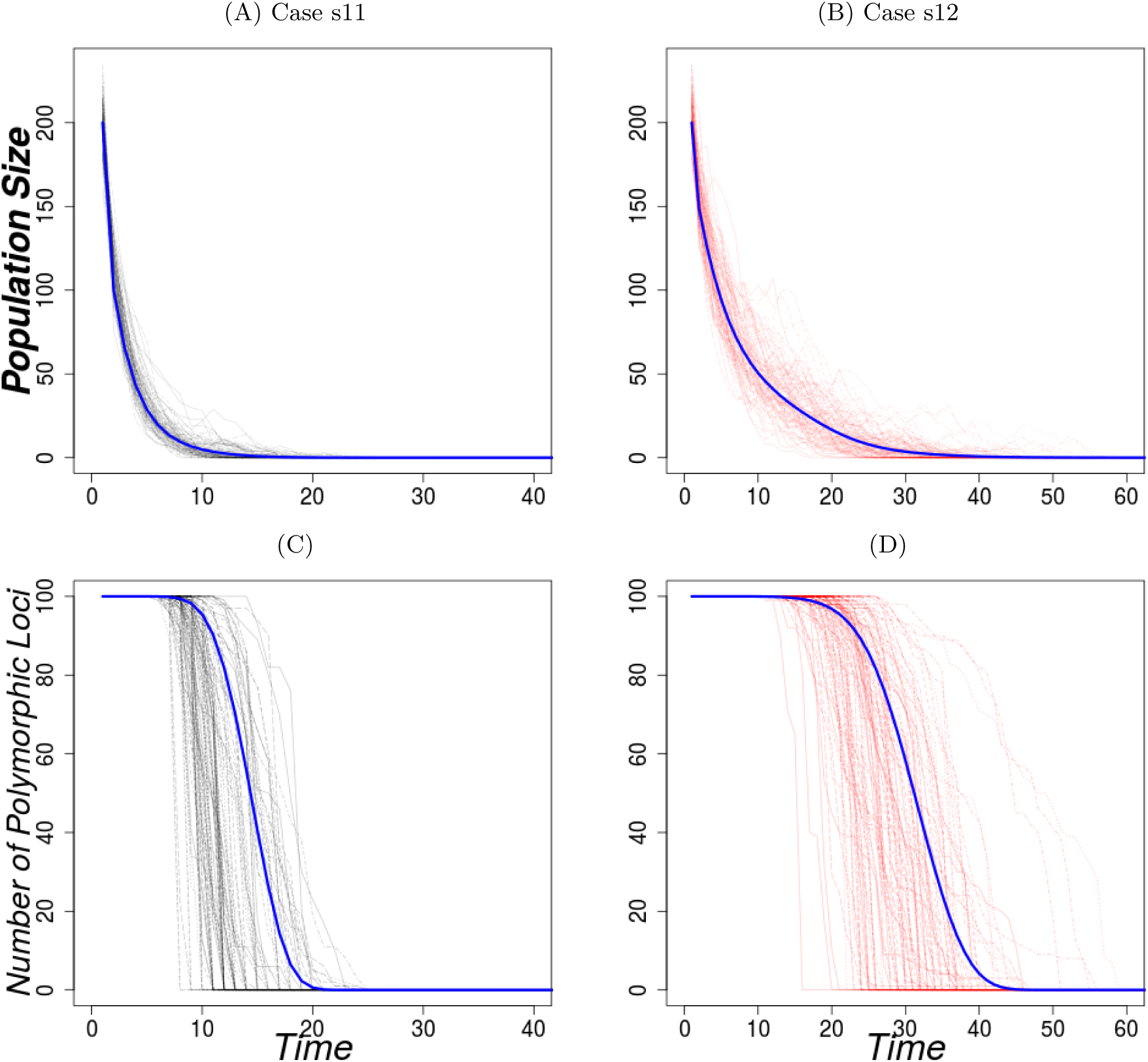
Variation of the population size and the number of polymorphic loci with time under the geometric growth model with fluctuating selection for Region I supplementary points. The dark blue lines show the analytic approximation with discretization parameters *u* = 0.1 and *τ* = 1. The parameters used for the 1st column (Case s11) are *r* = 1.30, *s* = 0.007 and for the 2nd column (Case s12 *r* = 1.15, *s* = 0.003. Other parameters are *N*_0_ = *K* = 200, *replicates* = 100.

**Figure A.14:**
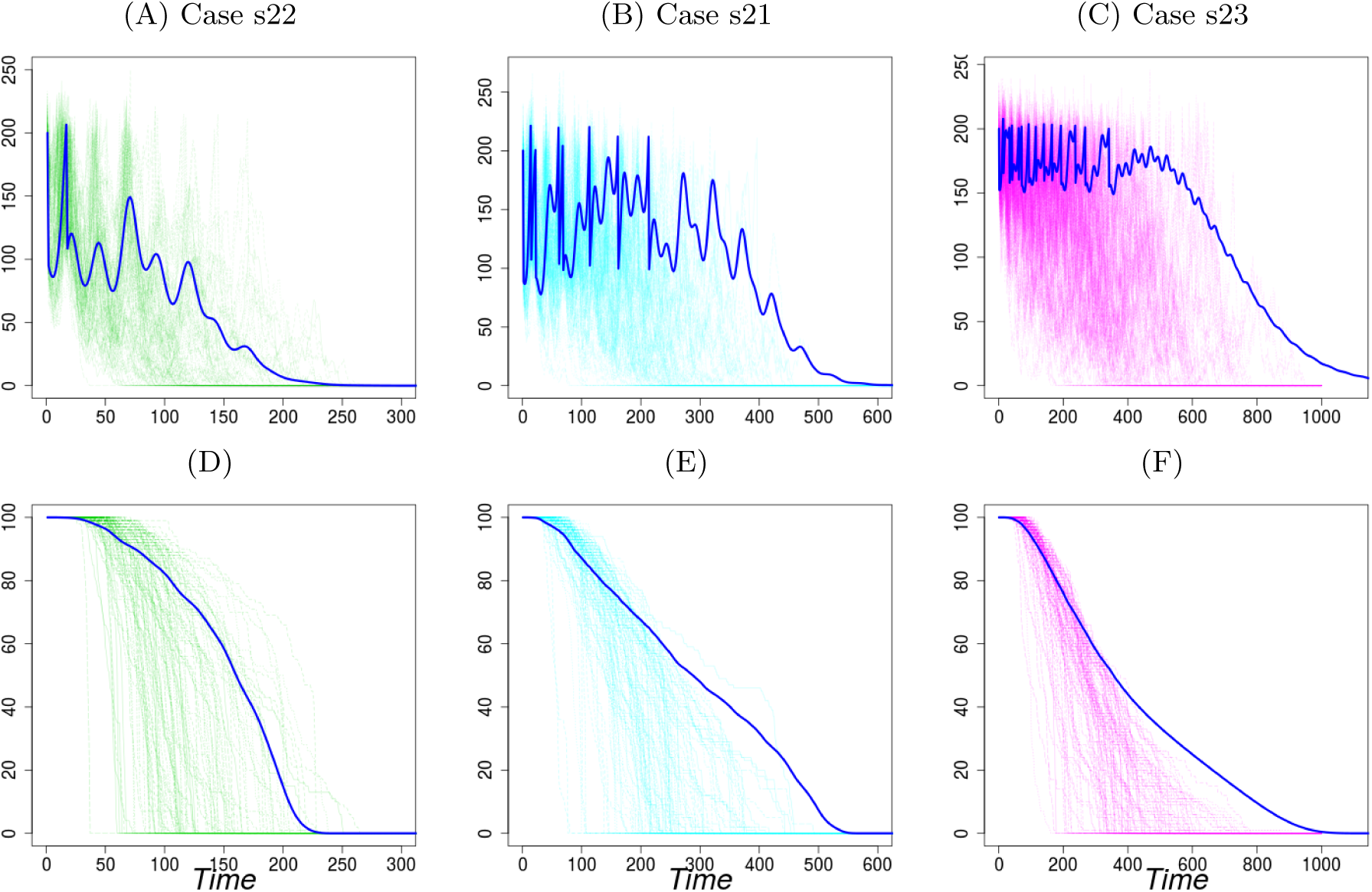
Variation of the population size and the number of polymorphic loci with time under the geometric growth model with fluctuating selection for Region II supplementary points. The dark blue lines show the analytic approximation with discretization parameters *u* = 0.1 and *τ* = 1. The parameters used for the 1st column (Case s11) *r* = 2.00, *s* = 0.0075 and for the 2nd column (Case s12) *r* = 2.20, *s* = 0.0082 and 3rd column (Case s13) *r* = 1.3, *s* = 0.0027. Other parameters are *N*_0_ = *K* = 200, *replicates* = 100.

**Figure A.15:**
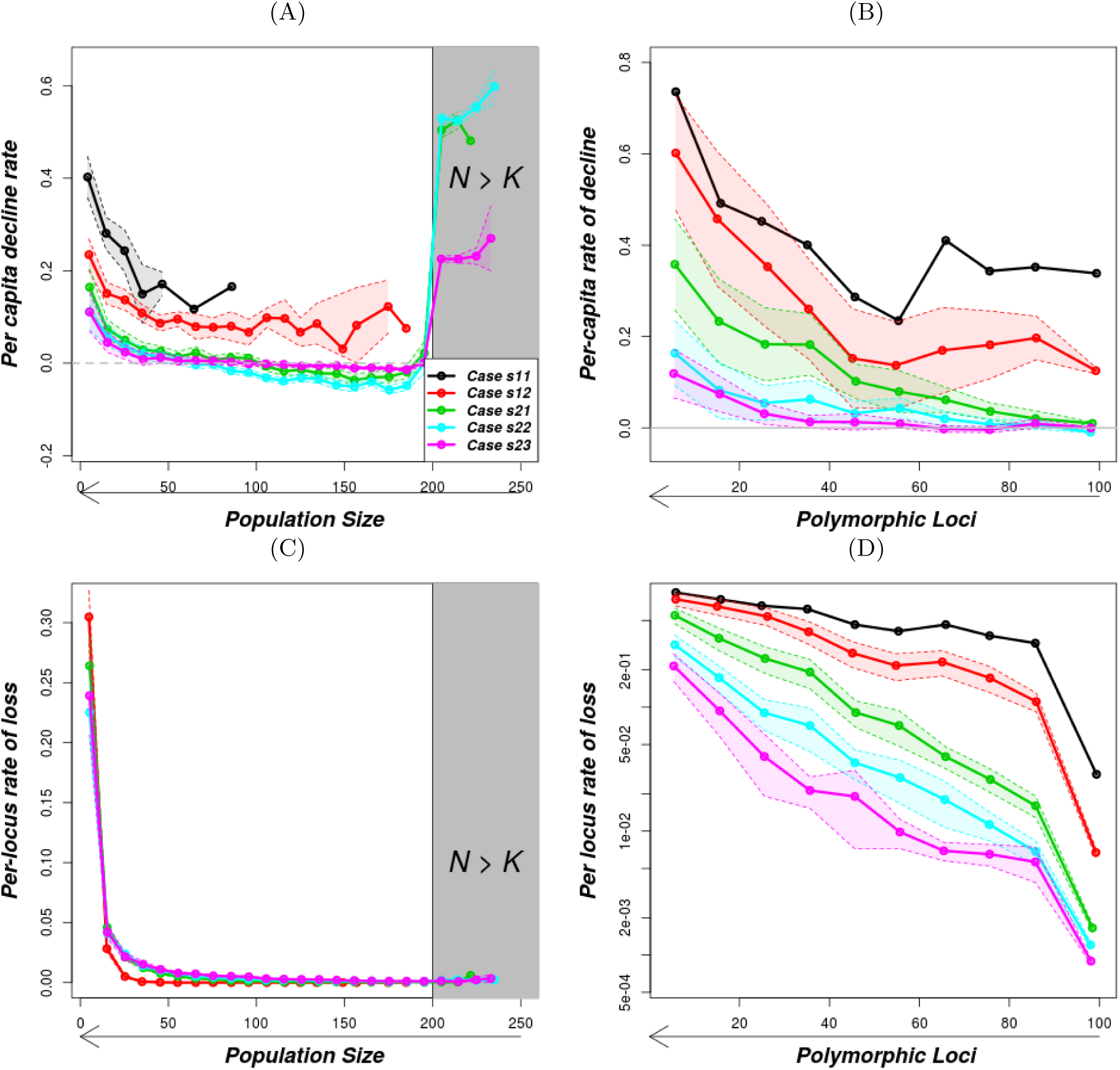
Per-capita decline rate and per-locus rate of loss as population size and the number of polymorphic loci varies under geometric growth model with fluctuating selection for Region I and II supplementary points. The parameters used are as on Figures A.13 and A.14 for the respective cases.

##### A.6.2 Early-warning signals

**Figure A.16:**
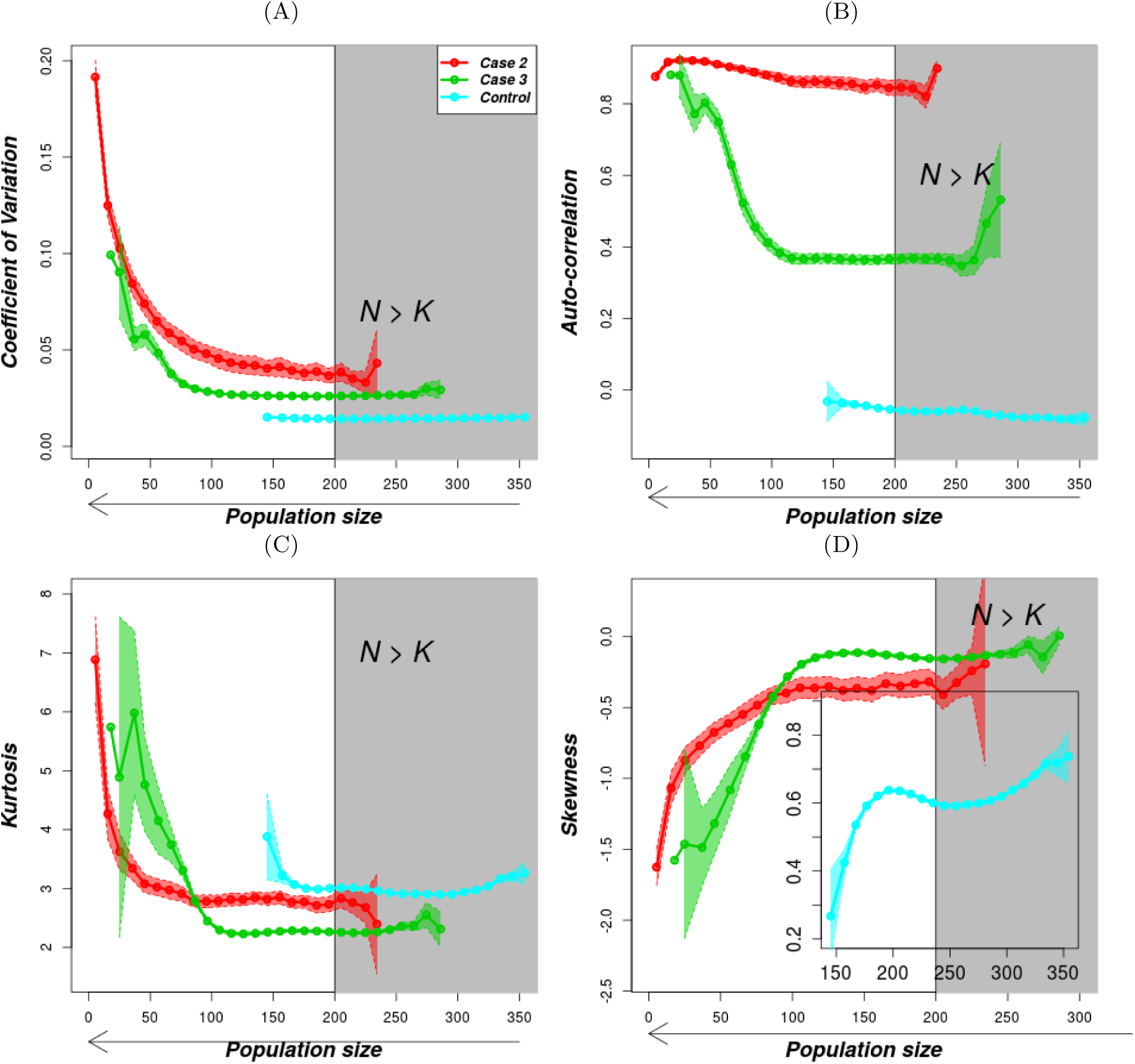
Variation of early-warning indicators with population size under the geometric population growth model and fluctuating selection with reversal of dominance. The solid lines are the mean values of the rates as described in the methods and the shaded regions show the standard error while the gray-shaded rectangle is the region above carrying capacity. The parameters for the 3 cases are as in Figure 6.

## References

Ballou, J., and K. Ralls. 1982. Inbreeding and juvenile mortality in small populations of ungulates: a detailed analysis. Biological Conservation 24:239–272.

Bensch, S., H. Andrén, B. Hansson, H. C. Pedersen, H. Sand, D. Sejberg, P. Wabakken, M. Åkesson, and O. Liberg. 2006. Selection for heterozygosity gives hope to a wild population of inbred wolves. PloS One 1:e72.

Benson, J. F., P. J. Mahoney, T. W. Vickers, J. A. Sikich, P. Beier, S. P. Riley, H. B. Ernest, and W. M. Boyce. 2019. Extinction vortex dynamics of top predators isolated by urbanization. Ecological Applications 29:e01868.

Bertram, J., and J. Masel. 2019. Different mechanisms drive the maintenance of polymorphism at loci subject to strong versus weak fluctuating selection. Evolution 73:883–896.

Blomqvist, D., A. Pauliny, M. Larsson, and L.-Å. Flodin. 2010. Trapped in the extinction vortex? Strong genetic effects in a declining vertebrate population. BMC Evolutionary Biology 10:33.

Chen, J., V. Nolte, and C. Schlötterer. 2015. Temperature stress mediates decanalization and dominance of gene expression in *Drosophila melanogaster*. PLoS Genetics 11:e1004883.

Connallon, T., and S. F. Chenoweth. 2019. Dominance reversals and the maintenance of genetic variation for fitness. PloS Biology 17:e3000118.

Coron, C., S. Méléard, E. Porcher, and A. Robert. 2013. Quantifying the mutational meltdown in diploid populations. The American Naturalist 181:623–636.

Courchamp, F., E. Angulo, P. Rivalan, R. J. Hall, L. Signoret, L. Bull, and Y. Meinard. 2006. Rarity value and species extinction: The anthropogenic Allee effect. PloS Biology 4:e415.

Curtsinger, J. W., P. M. Service, and T. Prout. 1994. Antagonistic pleiotropy, reversal of dominance, and genetic polymorphism. The American Naturalist 144:210–228.

Dakos, V., and J. Bascompte. 2014. Critical slowing down as early warning for the onset of collapse in mutualistic communities. Proceedings of the National Academy of Sciences 111:17546–17551.

Dakos, V., S. R. Carpenter, W. A. Brock, A. M. Ellison, V. Guttal, A. R. Ives, S. Kefi, V. Livina, D. A. Seekell, E. H. van Nes, et al. 2012. Methods for detecting early warnings of critical transitions in time series illustrated using simulated ecological data. PloS One 7:e41010.

de Silva, S., and P. Leimgruber. 2019. Demographic tipping points as early indicators of vulnerability for slow-breeding megafaunal populations. Frontiers in Ecology and Evolution 7:171.

Drake, J. M., and B. D. Griffen. 2010. Early warning signals of extinction in deteriorating environments. Nature 467:456–459.

Durrett, R. 2008. Probability models for DNA sequence evolution. Springer Science & Business Media.

Fagan, W. F., and E. Holmes. 2006. Quantifying the extinction vortex. Ecology Letters 9:51–60.

Frankham, R. 2005. Genetics and extinction. Biological Conservation 126:131–140.

Gilpin, M. E., and M. E. Soulé. 1986. Minimal viable populations: processes of species extinction. Conservation Biology: the science of scarcity and diversity pages 19–34.

Grieshop, K., and G. Arnqvist. 2018. Sex-specific dominance reversal of genetic variation for fitness. PloS Biology 16:e2006810.

Gsell, A. S., U. Scharfenberger, D. Özkundakci, A. Walters, L.-A. Hansson, A. B. Janssen, P. Nõges, P. C. Reid, D. E. Schindler, E. Van Donk, et al. 2016. Evaluating early-warning indicators of critical transitions in natural aquatic ecosystems. Proceedings of the National Academy of Sciences 113:E8089–E8095.

Harmon, L. J., and S. Braude. 2010. Conservation of small populations: effective population sizes, inbreeding, and the 50/500 rule. An introduction to methods and models in ecology, evolution, and conservation biology pages 125–138.

Jarvis, L., K. McCann, T. Tunney, G. Gellner, and J. M. Fryxell. 2016. Early warning signals detect critical impacts of experimental warming. Ecology and Evolution 6:6097–6106.

Kim, B.-J., B.-K. Lee, H. Lee, and G.-S. Jang. 2016. Considering threats to population viability of the endangered Korean long-tailed goral (*Naemorhedus caudatus*) using VORTEX. Animal Cells and Systems 20:52–59.

Kimura, M., et al. 1955. Stochastic processes and distribution of gene frequencies under natural selection. Cold Spring Harbor Symposia on Quantitative Biology 20:33–53.

Lacy, R. C. 1993. VORTEX - a computer simulation model for population viability analysis. Wildlife Research 20:45–65.

Luque, G. M., C. Vayssade, B. Facon, T. Guillemaud, F. Courchamp, and X. Fauvergue. 2016. The genetic Allee effect: a unified framework for the genetics and demography of small populations. Ecosphere 7:e01413.

Lynch, M., and W. Gabriel. 1990. Mutation load and the survival of small populations. Evolution 44:1725–1737.

Miller, W., V. M. Hayes, A. Ratan, D. C. Petersen, N. E. Wittekindt, J. Miller, B. Walenz, J. Knight, J. Qi, F. Zhao, et al. 2011. Genetic diversity and population structure of the endangered marsupial *Sarcophilus harrisii* (Tasmanian devil). Proceedings of the National Academy of Sciences of the United States of America 108:12348–12353.

Palomares, F., J. A. Godoy, J. V. López-Bao, A. RodrÍguez, S. Roques, M. Casas-Marce, E. Revilla, and M. Delibes. 2012. Possible extinction vortex for a population of Iberian lynx on the verge of extirpation. Conservation Biology 26:689–697.

R Core Team. 2015. R: A Language and Environment for Statistical Computing. R Foundation for Statistical Computing, Vienna, Austria.

Ralls, K., K. Brugger, and J. Ballou. 1979. Inbreeding and juvenile mortality in small populations of ungulates. Science 206:1101–1103.

Reed, D. H., and R. Frankham. 2003. Correlation between fitness and genetic diversity. Conservation Biology 17:230–237.

Saccheri, I., M. Kuussaari, M. Kankare, P. Vikman, W. Fortelius, and I. Hanski. 1998. Inbreeding and extinction in a butterfly metapopulation. Nature 392:491–494.

Sommer, S., K. J. van Benthem, D. Fontaneto, and A. Ozgul. 2017. Are generic early-warning signals reliable indicators of population collapse in rotifers? Hydrobiologia 796:111–120.

Spielman, D., B. W. Brook, D. A. Briscoe, and R. Frankham. 2004. Does inbreeding and loss of genetic diversity decrease disease resistance? Conservation Genetics 5:439–448.

Tanaka, Y. 1997. Extinction of populations due to inbreeding depression with demographic distur-bances. Researches on Population Ecology 39:57–66.

Tanaka, Y. 1998. Theoretical aspects of extinction by inbreeding depression. Population Ecology 40:279–286.

Tanaka, Y. 2000. Extinction of populations by inbreeding depression under stochastic environments. Population Ecology 42:55–62.

Wang, T., M. Fujiwara, X. Gao, and H. Liu. 2019. Minimum viable population size and population growth rate of freshwater fishes and their relationships with life history traits. Scientific Reports 9:3612.

Wittmann, M. J., A. O. Bergland, M. W. Feldman, P. S. Schmidt, and D. A. Petrov. 2017. Seasonally fluctuating selection can maintain polymorphism at many loci via segregation lift. Proceedings of the National Academy of Sciences 114:E9932–E9941.

Wittmann, M. J., H. Stuis, and D. Metzler. 2018. Genetic Allee effects and their interaction with ecological Allee effects. Journal of Animal Ecology 87:11–23.

Xu, S., M. Chen, C. Liu, R. Zhang, and X. Yue. 2019. Behavior of different numerical schemes for random genetic drift. BIT Numerical Mathematics pages 1–25.

Zhao, L., X. Yue, and D. Waxman. 2013. Complete numerical solution of the diffusion equation of random genetic drift. Genetics 194:973–985.

