## Supplementary material for "Models for Eco-evolutionary Extinction Vortices and their Detection": R scripts for regenerating data used in the manuscript: AnalysisFunctionsSubmit.R

```
#This section calculates the per-capita decline rate. 
#It takes in an array of population sizes in 3-DIMENSION. The first dimension is number of geneŕations, second is number of replicates and 
#3rd is the number of Cases combined for a given growth model and selection mechanism. 
# We first load our saved RData into a 3-Dimension Array assuming the data and this script are saved in the same folder/directory.
# Otherwise, provide the right path to the RData data

# Some of the packages needed 
library("moments", lib.loc="~/R/x86_64-pc-linux-gnu-library/3.2")
library("Hmisc", lib.loc="~/R/x86_64-pc-linux-gnu-library/3.2") 


#fixed parameters throught the manuscript
replicates = 100 # Number of replicates
numberofloci <- 100 # Number of loci
d0 = 0.0 # death rate. In this study, it is set to zero. No death rate considered
p = 0.5 # initial frequency of allele "A" in the population
percent = 20 # Parameter used during fluctuating selection scenario. It is denoted as c in the methods. Here it is expressed as a percentage.
mut_a = 0 # mutation rate for allele a. Set to 0 throught.
mut_A = 0# mutation rate for allele A. Set to 0 throught.
kapa = 50#

#Varrying parameters, assuming the parameters below where used  
geometric = 1 # 1 selects for geometric growth rate other values select for Ricker model
constantselection = 1 # One selects for heterozygote advantage. Other values select for fluctuating selction with reversal of dominance. 
tmax <- 3000 # Time for running simulations. This worked very well for N0 = K = 200 but 30000 worked well for N0 = K = 2000. The code stops when either population is extinct or this time is reached.
maxsels = 0.005 # selection coefficient
maxselt = maxsels
Founderpop <- 200 # starting population size
K <- Founderpop # Founder population size denoted as N_0 in the text used as carrying capacity
rate = log(c(1.2, 1.5, 1.8)) # Intrinsic growth rate. In the main text, it is denoted as r. Note that you can run one case at a time


POPULATIONSIZESARRAY = array(0, dim = c(tmax, replicates, length(rate)))
POLYMORPHICLOCIARRAY = array(0, dim = c(tmax, replicates, length(rate)))

#Compiling into 3-D array
for (case in 1:length(rate)){
  
 POPULATIONSIZESARRAY[,,case] = get(load(paste("pathToFolder/POPSIZE_r0_log",exp(rate[case]), "_N0_", Founderpop, "_selA_sela_", maxsels, "_mut_", mut_a, "_rep_",replicates, "_KN0_p_", p,"_", numberofloci,"loci", "_ConstantSelection_", constantselection, "_Geometric_",geometric, ".RData", sep = "")) )[1:tmax,]
 POLYMORPHICLOCIARRAY[,,case] = get(load(paste("pathToFolder/POLYLOCI_r0_log",exp(rate[case]), "_N0_", Founderpop, "_selA_sela_", maxsels, "_mut_", mut_a, "_rep_",replicates, "_KN0_p_", p,"_", numberofloci,"loci", "_ConstantSelection_", constantselection, "_Geometric_",geometric, ".RData", sep = "")) )[1:tmax,]
 

 }


Pp = POPULATIONSIZESARRAY# Array of 3 dimension
Hh = POLYMORPHICLOCIARRAY# Array of 3 dimension


#================= Now the analysis 

widthP = 10 # population size class width during partitioning

# For all cases combined for a given selection mechanism and population growth model
PSIZE = list()# Population size
HPOLY = list() # Polymorphic loci
DDIF = list()#Per-capita rate
DDIFH = list()#per-locus rate
DDIFVAR = list()# standard error in per-capita
PSIZEVAR = list()# standard error in population size
DDIFHVAR = list()# standard error in per-locus rate
HPOLYVAR = list()# standard error in polymorphic loci

# For specific cases under a given selection mechanism and population growth model above
for (Case in 1:dim(Pp)[3]){
  BinYX = c(seq(0, max(Pp[,,Case], na.rm = T), widthP))
  PSIZE1 = matrix(NA, ncol = replicates, nrow = (length(BinYX)-1)) # Population size
  HPOLY1 = matrix(NA, ncol = replicates, nrow = (length(BinYX)-1))# polymorphic loci
  DDIF1 = matrix(NA, ncol = replicates, nrow = (length(BinYX)-1))# per-capita rate
  DDIFH1 = matrix(NA, ncol = replicates, nrow = (length(BinYX)-1))# Per-locus rate
  
  # For each replicatesicate under each case
  for (re in 1:replicates){
    yy = Pp[,re,Case]
    xx = -diff(Pp[,re,Case])/(Pp[-dim(Pp)[1],re,Case]) # Per-capita rate at replicatesicate level
    yyH = Hh[,re,Case]
    xxH = -diff(Hh[,re,Case])/(Hh[-dim(Pp)[1],re,Case]) # Per-locus rate at replicatesicate level
    Y=as.vector(t(yy))
    X = c(as.vector(t(xx)), rep(0,1))
    YX = cbind(Y,X)
    YH=as.vector(t(yyH))
    XH = c(as.vector(t(xxH)), rep(0,1))
    YXH = cbind(YH,XH)
    ###
    DDif = c()
    PSize= c()
    
    DDifH = c()
    PSizeH= c()
    DDif = c()
    PSize = c()
    DDifH = c()
    HPoly = c()
    
    DDifVar = c()
    PSizeVar = c()
    DDifHVar = c()
    
    nnH = 0
    
    while(nnH < max(BinYX)){
      x = YX[,1]
      y = YX[,2]
      yH = YXH[,2]
      yHH = YXH[,1]
      YYH =  x[which(x>nnH & x<=(nnH + widthP))]  
      XXH = y[which(x>nnH & x<=(nnH + widthP))]
      XXHH = yH[which(x>nnH & x<=(nnH + widthP))]
      Hcorres = yHH[which(x>nnH & x<=(nnH + widthP))]  
      
      
      if (length(XXH)>1){
        HPoly = c(HPoly, mean(Hcorres[!is.infinite(Hcorres)], na.rm = T))
        
        DDif = c(DDif, mean(XXH[!is.infinite(XXH)], na.rm = T))
        PSize = c(PSize, mean(YYH[!is.infinite(YYH)], na.rm = T))
        DDifH = c(DDifH, mean(XXHH[!is.infinite(XXHH)], na.rm = T))
        
      }else{
        HPoly = c(HPoly,NA)
        DDif =c(DDif,NA) 
        PSize = c(PSize, NA)
        DDifH = c(DDifH, NA)
      }
      
     nnH = nnH + widthP
    }
    PSIZE1[,re] = PSize 
    HPOLY1[,re] = HPoly
    DDIF1[,re ] = DDif
    DDIFH1[,re ] = DDifH
    
  }
  
  # Final Means
  PSIZE[[Case]] = rowMeans(PSIZE1, na.rm = TRUE) # Population size
  HPOLY[[Case]] = rowMeans(HPOLY1, na.rm = TRUE)# Polymorphic Loci
  DDIF[[Case]] = rowMeans(DDIF1, na.rm = TRUE)# Per-capita growth rate
  DDIFH[[Case]] = rowMeans(DDIFH1, na.rm = TRUE)#Per-locus rate of loss
  
  #Standard Errors
  PSIZEVAR[[Case]] = apply(PSIZE1, 1, function(x) sd(x, na.rm = TRUE)/sqrt(length(x[!is.na(x)])))# for population size
  HPOLYVAR[[Case]] = apply(HPOLY1, 1, function(x) sd(x, na.rm = TRUE)/sqrt(length(x[!is.na(x)])))# for polymorphic loci
  DDIFVAR[[Case]] = apply(DDIF1, 1, function(x) sd(x, na.rm = TRUE)/sqrt(length(x[!is.na(x)])))# for per-capita rate
  DDIFHVAR[[Case]] = apply(DDIFH1, 1, function(x) sd(x, na.rm = TRUE)/sqrt(length(x[!is.na(x)])))# for per-locus rate of loss
}


#Plots example for 4 cases. That is, Case 1, 2, 3 and the control case.

########===========Heterozygous Advantage, N0 = 200 ===================
cala = c(1,2, 3, 5)# different colours

# ======= Per-capita rate vs Population size =============
i = 1
plot(PSIZE[[i]][!is.nan(DDIF[[i]])], DDIF[[i]][!is.nan(DDIF[[i]])],  type = 'n', col = cala[i], lwd = 3, xlim = c(5,350), ylim = c(-0.5,0.5), xlab = list("Population Size", font = 4, col = 1, cex = 1.5), ylab = list("Per capita decline rate", font = 4, col = 1, cex = 1.5))
rect(200, -10, 400, 10 , density = -0.1 , angle = 45, col = "gray", border = NULL, lty = par("lty"), lwd = par("lwd"))

for (i in 1:dim(Pp)[3]){
  lines(PSIZE[[i]][!is.na(DDIFVAR[[i]])], DDIF[[i]][!is.na(DDIFVAR[[i]])]-2*DDIFVAR[[i]][!is.na(DDIFVAR[[i]])], col = cala[i], lwd = 1, lty = 2)
  lines(PSIZE[[i]][!is.na(DDIFVAR[[i]])], DDIF[[i]][!is.na(DDIFVAR[[i]])]+2*DDIFVAR[[i]][!is.na(DDIFVAR[[i]])], col = cala[i], lwd = 1, lty = 2)
  lines(PSIZE[[i]][!is.nan(DDIF[[i]])], DDIF[[i]][!is.nan(DDIF[[i]])], type = 'o', col = cala[i], lwd = 3)
  polygon(c(PSIZE[[i]][!is.na(DDIFVAR[[i]])],rev(PSIZE[[i]][!is.na(DDIFVAR[[i]])])),c(DDIF[[i]][!is.na(DDIFVAR[[i]])]-2*DDIFVAR[[i]][!is.na(DDIFVAR[[i]])],rev(DDIF[[i]][!is.na(DDIFVAR[[i]])]+2*DDIFVAR[[i]][!is.na(DDIFVAR[[i]])])), border = NA, col = adjustcolor(cala[i], alpha.f = 0.1) )#rgb(1, 0, 0,0.5))
}

abline(h = 0,  lwd = 2, lty = 2, col = 8)
text(250,-0.26, "N > K", cex = 2.0, srt = 360, font = 3)
arrows(350, -0.65, 0, -0.65, xpd = TRUE)

legend("bottomright", legend = c(  "Case 1", "Case 2", "Case 3", "Control"), lwd = 3, col = cala, text.font = 4, pch = 1)

# ====== Per-locus rate vs population size
i = 1
plot(PSIZE[[i]][!is.nan(PSIZE[[i]])], DDIFH[[i]][!is.nan(DDIFH[[i]])], type = 'n', col = cala[i],  lwd = 3,ylim = c(0.0000099,0.341), xlim = c(5,350), xlab = list("Population Size", font = 4, col = 1, cex = 1.5), ylab = list("Per-locus rate of loss", font = 4, col = 1, cex = 1.5))

rect(200, -10.0000099, 400, 0.5, density = -0.1 , angle = 45, col = "gray", border = NULL, lty = par("lty"), lwd = par("lwd"))
for (i in 1:dim(Pp)[3]){
  lines(PSIZE[[i]][!is.na(DDIFH[[i]])], DDIFH[[i]][!is.na(DDIFH[[i]])], type = 'o', col = cala[i], lwd = 3)
  lines(PSIZE[[i]][!is.na(DDIFHVAR[[i]])], DDIFH[[i]][!is.na(DDIFHVAR[[i]])]-2*DDIFHVAR[[i]][!is.na(DDIFHVAR[[i]])], col = cala[i], lwd = 1, lty = 2)
  lines(PSIZE[[i]][!is.na(DDIFHVAR[[i]])], DDIFH[[i]][!is.na(DDIFHVAR[[i]])]+2*DDIFHVAR[[i]][!is.na(DDIFHVAR[[i]])], col = cala[i], lwd = 1, lty = 2)
  polygon(c(PSIZE[[i]][!is.na(DDIFHVAR[[i]])],rev(PSIZE[[i]][!is.na(DDIFHVAR[[i]])])),c(DDIFH[[i]][!is.na(DDIFHVAR[[i]])]-2*DDIFHVAR[[i]][!is.na(DDIFHVAR[[i]])],rev(DDIFH[[i]][!is.na(DDIFHVAR[[i]])]+2*DDIFHVAR[[i]][!is.na(DDIFHVAR[[i]])])), border = NA, col = adjustcolor(cala[i], alpha.f = 0.1) )
  
}


text(250,0.1, "N > K", cex = 2.0, srt = 360, font = 3)

arrows(350, -0.05, 0, -0.05, xpd = TRUE)


#######################################################
#This section calculates the per-locus rate of loss. 
#It takes in an array of number Polymorphic loci whose first dimension is number of geneŕations, second is number of replicates and 
#3rd is the number of Cases combined for a given growth model and selection mechanism. 

Pp = POPULATIONSIZESARRAY
Hh = POLYMORPHICLOCIARRAY

replicates = dim(Pp)[2]
widthH = 10 # Class width for number of polymorphic loci during partitioning

PPSIZE = list()# Population size
NH = list()#polymorphic loci
DDIFN = list()#per-capita rate
DDIFHH = list()#per-locus rate
DDIFHHVAR = list() # std error in per-locus rate of loss
NHVAR = list() # std error in number of polymorphic loci
DDIFNVAR = list()#std error in per-capita rate
PPSIZEVAR = list()#std error in population size

for (Case in 1:dim(Hh)[3]){
  
  BinYXH = seq(0,max(YXH[,1], na.rm = T), widthH)
  
  PPSIZE1 = matrix(NA, ncol = replicates, nrow = (length(BinYXH)-1))
  NH1 = matrix(NA, ncol = replicates, nrow = (length(BinYXH)-1))
  DDIFN1 = matrix(NA, ncol = replicates, nrow = (length(BinYXH)-1))
  DDIFHH1 = matrix(NA, ncol = replicates, nrow = (length(BinYXH)-1))
 
  
  for (re in 1:replicates){
    yy = Pp[,re,Case]
    xx = -diff(Pp[,re,Case])/(Pp[-dim(Pp)[1],re,Case])
    yyH = Hh[,re,Case]
    xxH = -diff(Hh[,re,Case])/(Hh[-dim(Hh)[1],re,Case])
    Y=as.vector(t(yy))
    X = c(as.vector(t(xx)), rep(0,1))
    YX = cbind(Y,X)
    YH=as.vector(t(yyH))
    XH = c(as.vector(t(xxH)), rep(0,1))
    YXH = cbind(YH,XH)
    ###
    DDifHH = c()
    N_H= c()
    DDifN = c()
    PSizeH= c()
    PPSize = c()
    DDifHHVar = c()
    N_HVar = c()
    DDifNVar = c()
    nn = 0
    
    while(nn < max(BinYXH)){
      y = YX[,2]
      yH = YXH[,2]
      xH = YXH[,1]
      xHN = YX[,1]
      YYH =  xH[which(xH>nn & xH<=(nn + widthH))]  
      XXH = yH[which(xH>nn & xH<=(nn + widthH))]
      XXHN = y[which(xH>nn & xH<=(nn + widthH))]
      PSizeCorres = xHN[which(xH>nn & xH<=(nn + widthH))]
      
      PPSize = c(PPSize, mean(PSizeCorres[!is.infinite(PSizeCorres)]))
      DDifHH = c(DDifHH, mean(XXH[!is.infinite(XXH)], na.rm = T))
      N_H = c(N_H, mean(YYH[!is.infinite(YYH)], na.rm = T))
      DDifN = c(DDifN, mean(XXHN[!is.infinite(XXHN)], na.rm = T))
      
      nn = nn + widthH
    }
    PPSIZE1[,re] = PPSize
    NH1[,re] = N_H
    DDIFN1[,re] = DDifN
    DDIFHH1[,re] = DDifHH
    
  }
  PPSIZE[[Case]] = rowMeans(PPSIZE1, na.rm = TRUE)
  NH[[Case]] = rowMeans(NH1, na.rm = TRUE) 
  DDIFN[[Case]] = rowMeans(DDIFN1, na.rm = TRUE)
  DDIFHH[[Case]] = rowMeans(DDIFHH1, na.rm = TRUE)
  
  DDIFHHVAR[[Case]] = apply(DDIFHH1, 1, function(x) sd(x, na.rm = TRUE)/sqrt(length(x[!is.na(x)])))
  NHVAR[[Case]] = apply(NH1, 1,  function(x) sd(x, na.rm = TRUE)/sqrt(length(x[!is.na(x)])))
  DDIFNVAR[[Case]] = apply(DDIFN1, 1,  function(x) sd(x, na.rm = TRUE)/sqrt(length(x[!is.na(x)])))
  PPSIZEVAR[[Case]] = apply(PPSIZE1, 1, function(x) sd(x, na.rm = TRUE)/sqrt(length(x[!is.na(x)])))
}


#Plots 
# =========== Per-locus rate vs polymorphic loci ==================================
i = 1
plot(NH[[i]][!is.nan(NH[[i]])], DDIFHH[[i]][!is.nan(DDIFHH[[i]])], type = 'o', log = 'y', col = cala[i], lwd = 3, xlim = c(4,100), ylim = c(0.0006,0.75), xlab = list(" Polymorphic Loci", font = 4, col = 1, cex = 1.5), ylab = list("Per locus rate of loss", font = 4, col = 1, cex = 1.5))
for (i in 1:dim(Pp)[3]){
  lines(NH[[i]][!is.nan(NH[[i]])], DDIFHH[[i]][!is.nan(DDIFHH[[i]])], type = 'o', col = cala[i], lwd = 3) 
  lines(NH[[i]][!is.nan(NH[[i]])], DDIFHH[[i]][!is.nan(DDIFHH[[i]])]-2*DDIFHHVAR[[i]][!is.nan(DDIFHHVAR[[i]])], col = cala[i], lwd = 1, lty = 2)
  lines(NH[[i]][!is.nan(NH[[i]])], DDIFHH[[i]][!is.nan(DDIFHH[[i]])]+2*DDIFHHVAR[[i]][!is.nan(DDIFHHVAR[[i]])], col = cala[i], lwd = 1, lty = 2)
  polygon(c(NH[[i]][!is.nan(NH[[i]])],rev(NH[[i]][!is.nan(NH[[i]])])),c(DDIFHH[[i]][!is.nan(DDIFHH[[i]])]-2*DDIFHHVAR[[i]][!is.nan(DDIFHHVAR[[i]])],rev(DDIFHH[[i]][!is.nan(DDIFHH[[i]])]+2*DDIFHHVAR[[i]][!is.nan(DDIFHHVAR[[i]])])), border = NA, col = adjustcolor(cala[i], alpha.f = 0.1) )
  
}

arrows(100, 0.0002, 0, 0.0002, xpd = TRUE)


# =========== Per-locus rate vs population size ==================================
i = 1
plot(NH[[i]][!is.nan(NH[[i]])], DDIFN[[i]][!is.nan(DDIFN[[i]])],  type = 'o', col = cala[i], lwd = 3, xlim = c(4,100), ylim = c(-0.0406,0.55), xlab = list(" Polymorphic Loci", font = 4, col = 1, cex = 1.5), ylab = list("Per-capita rate of decline", font = 4, col = 1, cex = 1.5))

for (i in 1:dim(Pp)[3]){
  lines(NH[[i]][!is.nan(NH[[i]])], DDIFN[[i]][!is.nan(DDIFN[[i]])], type = 'o', col = cala[i], lwd = 3)
  lines(NH[[i]][!is.nan(NH[[i]])], DDIFN[[i]][!is.nan(DDIFN[[i]])]-2*DDIFNVAR[[i]][!is.nan(DDIFNVAR[[i]])], col = cala[i], lwd = 1, lty = 2)
  lines(NH[[i]][!is.nan(NH[[i]])], DDIFN[[i]][!is.nan(DDIFN[[i]])]+2*DDIFNVAR[[i]][!is.nan(DDIFNVAR[[i]])], col = cala[i], lwd = 1, lty = 2)
  polygon(c(NH[[i]][!is.nan(NH[[i]])],rev(NH[[i]][!is.nan(NH[[i]])])),c(DDIFN[[i]][!is.nan(DDIFN[[i]])]-2*DDIFNVAR[[i]][!is.nan(DDIFNVAR[[i]])],rev(DDIFN[[i]][!is.nan(DDIFN[[i]])]+2*DDIFNVAR[[i]][!is.nan(DDIFNVAR[[i]])])), border = NA, col = adjustcolor(cala[i], alpha.f = 0.1) )
  
}


arrows(100, -0.13, 0, -0.13, xpd = TRUE)
abline(h = 0, lwd = 2, lty = 2, col =8)

# ====================== End of analysis for per-capita rate of decline and per-locus rate of loss
#=================================================
# 
#=========================================================0
#========================================================
######======================== Early Warning signals =====================================
# This section estimates early-warning signals =============================================
############################################################################################
Pp = POPULATIONSIZESARRAY
Hh = POLYMORPHICLOCIARRAY
replicates = dim(Pp)[2] # number of replicates
WindowRange = 100 # Window range
CCM1 = array(NA, dim = c((dim(Pp)[1]-WindowRange), replicates, dim(Pp)[3]))# Auto-correlation at lag 1
SKK1 = array(NA, dim = c((dim(Pp)[1]-WindowRange), replicates, dim(Pp)[3]))# Skewness
KTS1 = array(NA, dim = c((dim(Pp)[1]-WindowRange), replicates, dim(Pp)[3]))# Kurtosis
StdDev1 = array(NA, dim = c((dim(Pp)[1]-WindowRange), replicates, dim(Pp)[3]))# Std deviation
CoeVar1 = array(NA, dim = c((dim(Pp)[1]-WindowRange), replicates, dim(Pp)[3])) # Coefficient of variation
PP1  = array(NA, dim = c((dim(Pp)[1]-WindowRange), replicates, dim(Pp)[3])) #New population sizes from the WindowRangeth generation to extinction.


#log transformation of population size before analysis
Pp = log(POPULATIONSIZESARRAY)


for (Case in 1:dim(Pp)[3]){
  
  CCM = matrix(NA,nrow = (dim(Pp)[1]-WindowRange), ncol = replicates)
  SKK = matrix(NA,nrow = (dim(Pp)[1]-WindowRange), ncol = replicates)
  KTS = matrix(NA,nrow = (dim(Pp)[1]-WindowRange), ncol = replicates)
  StdDev = matrix(NA,nrow = (dim(Pp)[1]-WindowRange), ncol = replicates)
  CoeVar = matrix(NA,nrow = (dim(Pp)[1]-WindowRange), ncol = replicates)
  PP = matrix(NA,nrow = (dim(Pp)[1]-WindowRange), ncol = replicates)
  
for (re in 1:replicates){
    
    ti = 1
    for (tt in WindowRange:(dim(Pp)[1]-1)){
      if((tt)<=(max(which(Pp[,re,Case]>=0)))){
        CCM[ti,re] = acf(log(Pp[ti:(tt-0),re,Case]), plot = FALSE, lag.max = 1, na.action = na.pass)$acf[2]
        SKK[ti,re] = skewness(log(Pp[ti:(tt-0),re,Case]), na.rm = T)
        KTS[ti, re] = kurtosis(log(Pp[ti:(tt-0),re,Case]), na.rm = T)
        StdDev[ti,re] = sd(log(Pp[ti:(tt-0),re,Case]), na.rm = T)
        CoeVar[ti,re] = StdDev[ti,re]/mean(log(Pp[ti:(tt-0),re,Case]), na.rm = T)
        PP[ti,re] = POPULATIONSIZESARRAY [(tt-0),re, Case]
        
      }
      
      ti = ti + 1
    }
    
  }
  CCM1[,,Case] = CCM 
  SKK1[,,Case] = SKK
  KTS1[,,Case] = KTS
  StdDev1[,,Case] = StdDev
  CoeVar1[,,Case] = CoeVar
  PP1[,,Case] = PP
  
}
klk
# ============== N0 = 200 for Heterozygous advantage =========================================

#Plots
Case = 4
ca = Case
ttb = 3000 # time-axis limit
rrep = 1:5 # number of replicates considered in the raw plots
Ma = matrix(NA, nrow = WindowRange, ncol = length(rrep))

# Population size 
par(mgp=c(2.3,1,0),mar=c(5,4,4,2)+0.1)
matplot(POPULATIONSIZESARRAY [,rrep, Case], lwd = 3, lty = 1, cex.axis = 1.5, type = 'l', xlim = c(0,ttb), ylim = c(0,max(POPULATIONSIZESARRAY [,rrep, Case], na.rm = TRUE)),  ylab = list("Population Size", font = 4, cex = 2.0, col =1), xlab = list("", font = 4, cex = 2.0, col =1))


#Auto-correlation

par(mgp=c(2.3,1,0),mar=c(5,4,4,2)+0.1)
matplot(rbind(Ma, CCM1[,rrep,Case]), lwd = 3, lty = 1, type = 'l', cex.axis = 1.5, xlim = c(0,ttb), ylim = c(-0.2,1),  ylab = list("Auto Correlation at lag 1", font = 4, cex = 1.5, col =1), xlab = list("Generation", font = 4, cex = 1.5, col =1))

# skewness
par(mgp=c(2.3,1,0),mar=c(5,4,4,2)+0.1)
matplot(rbind(Ma, SKK1[,rrep,Case]), lwd = 3, lty = 1, type = 'l', cex.axis = 1.5, xlim = c(0,ttb), ylim = c(-3.5,0.6), ylab = list("Skewness", font = 4, cex = 2, col =1), xlab = list("Generation", font = 4, cex = 2, col =1))

#Kurtosis
par(mgp=c(2.3,1,0),mar=c(5,4,4,2)+0.1)
matplot(rbind(Ma, KTS1[,rrep, Case]), lwd = 3, lty = 1, cex.axis = 1.5, type = 'l', xlim = c(0,ttb), ylim = c(2.,12.0),  ylab = list("Kurtosis", font = 4, cex = 2, col =1), xlab = list("Generation", font = 4, cex = 2, col =1))

#std deviation
par(mgp=c(2.3,1,0),mar=c(5,4,4,2)+0.1)
matplot(rbind(Ma, StdDev1[,rrep, Case]), lwd = 3, lty = 1, cex.axis = 1.5, type = 'l', xlim = c(0,ttb) , ylim = c(0.025,0.488), ylab = list("Standard Deviation", font = 4, cex = 2, col =1), xlab = list("Generation", font = 4, cex = 2, col =1))

#Coefficient of variation
par(mgp=c(2.3,1,0),mar=c(5,4,4,2)+0.1)
matplot(rbind(Ma, CoeVar1[,rrep, Case]), lwd = 3, lty = 1, type = 'l', cex.axis = 1.5, xlim = c(0,ttb), ylim = c(0.012,0.43), ylab = list("Coefficient of Variation", font = 4, cex = 2, col =1), xlab = list("Generation", font = 4, cex = 2, col =1))

# ===== End of raw plots of Early-warning indicators vs time

#########################################
#==========Analysis for early-warning indicators vs Population size ==================================
#=================================================

replicates = dim(Pp)[2]
widthP = 10 # Class width of the population size

PPSIZE = list()# populaztion size
PPSIZEsd = list()#std error population size
CCOEVAR = list()# Coefficient of variation (COV)
CCOEVARsd = list()#std error in COV

STDVAR = list()# std deviation
STDVARsd = list()# std error in std deviation

SKNESS = list()# Skewness
SKNESSsd = list()#std error in skewness

KTSS = list()#Kurtosis
KTSSsd = list()# std error in kurtosis

AUTO = list()# Auto-correlation at lag 1
AUTOsd = list()# std error in auto-correlation at lag 1

Pp = PP1 

for (Case in 2:dim(Pp)[3]){
  BinYX = c(seq(0,max(Pp[,,Case], na.rm = T), widthP))# partitioning new population size
  
  PPSIZE1 = matrix(NA, ncol = replicates, nrow = (length(BinYX)-1)) 
  CCOEVAR1 = matrix(NA, ncol = replicates, nrow = (length(BinYX)-1))
  STDVAR1 = matrix(NA, ncol = replicates, nrow = (length(BinYX)-1))
  SKNESS1 = matrix(NA, ncol = replicates, nrow = (length(BinYX)-1))
  KTSS1 = matrix(NA, ncol = replicates, nrow = (length(BinYX)-1))
  AUTO1 = matrix(NA, ncol = replicates, nrow = (length(BinYX)-1))
  
  for (re in 1:replicates){
    yy = Pp[,re,Case]
    
    CV = CoeVar1[,re,Case]
    SD = StdDev1[,re,Case] 
    SK = SKK1[,re,Case]
    KT = KTS1[,re,Case]
    AC = CCM1[,re,Case]
    
    Y = c(as.vector(t(yy)), rep(NA,WindowRange*1))
    XCV = c(as.vector(t(CV)), rep(NA,WindowRange*1))
    XSD = c(as.vector(t(SD)), rep(NA,WindowRange*1))
    XSK = c(as.vector(t(SK)), rep(NA,WindowRange*1))
    XKT = c(as.vector(t(KT)), rep(NA,WindowRange*1))
    XAC = c(as.vector(t(AC)), rep(NA,WindowRange*1))
    YX = cbind(Y,XCV, XSD, XSK, XKT, XAC)
    
    
    
    PPSize = c()
    
    CCoevar = c()
    CCoevarsd = c()
    
    CSD = c()
    CSDsd = c()
    
    STDvar = c()
    STDvarsd = c()
    
    CSK = c()
    CSKsd = c()
    
    CKT = c()
    CKTsd = c()
    
    CAC = c()
    CACsd = c()
    
    nn = 0
    
    while(nn < max(BinYX)){
      xHN = YX[,1]
      
      yCV = YX[,2]
      ySD = YX[,3]
      ySK = YX[,4]
      yKT = YX[,5]
      yAC = YX[,6]
      
      PSize =  xHN[which(xHN>nn & xHN<=(nn + widthP))]  
      
      
      coevar  = yCV[which(xHN>nn & xHN<=(nn + widthP))]
      stddev  = ySD[which(xHN>nn & xHN<=(nn + widthP))]
      
      sk  = ySK[which(xHN>nn & xHN<=(nn + widthP))]
      
      
      kt  = yKT[which(xHN>nn & xHN<=(nn + widthP))]
      
      
      ac  = yAC[which(xHN>nn & xHN<=(nn + widthP))]
      
      
      PPSize = c(PPSize, mean(PSize[!is.infinite(PSize)]))
      
      CCoevar = c(CCoevar, mean(coevar[!is.infinite(coevar)], na.rm = T))
      
      CSD = c(CSD, mean(stddev[!is.infinite(stddev)], na.rm = T))
      
      CSK = c(CSK, mean(sk[!is.infinite(sk)], na.rm = T))
      
      CKT = c(CKT, mean(kt[!is.infinite(kt)], na.rm = T))
      
      CAC = c(CAC, mean(ac[!is.infinite(ac)], na.rm = T))
      
      nn = nn + widthP
    }
    PPSIZE1[,re] = PPSize
    CCOEVAR1[,re] = CCoevar
    STDVAR1[,re] = CSD
    SKNESS1[,re] = CSK
    KTSS1[,re] = CKT
    AUTO1[,re] = CAC 
    
  }
  
  PPSIZE[[Case]] = rowMeans(PPSIZE1, na.rm = TRUE)
  PPSIZEsd[[Case]] = apply(PPSIZE1, 1, function(x) sd(x, na.rm = TRUE)/sqrt(length(x[!is.na(x)])))
  CCOEVAR[[Case]] = rowMeans(CCOEVAR1, na.rm = TRUE)
  CCOEVARsd[[Case]] = apply(CCOEVAR1, 1,  function(x) sd(x, na.rm = TRUE)/sqrt(length(x[!is.na(x)])))
  
  STDVAR[[Case]] = rowMeans(STDVAR1, na.rm = TRUE)
  STDVARsd[[Case]] = apply(STDVAR1, 1,  function(x) sd(x, na.rm = TRUE)/sqrt(length(x[!is.na(x)])))
  
  SKNESS[[Case]] = rowMeans(SKNESS1, na.rm = TRUE)
  SKNESSsd[[Case]] = apply(SKNESS1, 1,  function(x) sd(x, na.rm = TRUE)/sqrt(length(x[!is.na(x)])))
  
  KTSS[[Case]] = rowMeans(KTSS1, na.rm = TRUE)
  KTSSsd[[Case]] = apply(KTSS1, 1,  function(x) sd(x, na.rm = TRUE)/sqrt(length(x[!is.na(x)])))
  
  AUTO[[Case]] = rowMeans(AUTO1, na.rm = TRUE)
  AUTOsd[[Case]] = apply(AUTO1, 1,  function(x) sd(x, na.rm = TRUE)/sqrt(length(x[!is.na(x)])))
}


#Plots
cala = c(1,2, 3,5)

# Coefficient of variation
i = 1
plot(PPSIZE[[i]][!is.nan(CCOEVARsd[[i]])], CCOEVAR[[i]][!is.nan(CCOEVARsd[[i]])], type = 'n', col = cala[i], lwd = 3, ylim = c(0.0014,0.2), xlim = c(0,350), xlab = list("Population size", font = 4, col = 1, cex = 1.5), ylab = list("Coefficient of Variation", font = 4, col = 1, cex = 1.5))
rect(200, -0.014, 400, 10.23 , density = -0.1 , angle = 45, col = "gray", border = NULL, lty = par("lty"), lwd = par("lwd"))

for (i in 2:dim(Pp)[3]){
  lines(PPSIZE[[i]][!is.nan(CCOEVARsd[[i]])], CCOEVAR[[i]][!is.nan(CCOEVARsd[[i]])], type = 'o', col = cala[i], lwd = 3)
  lines(PPSIZE[[i]][!is.nan(CCOEVARsd[[i]])], CCOEVAR[[i]][!is.nan(CCOEVARsd[[i]])]-2*CCOEVARsd[[i]][!is.nan(CCOEVARsd[[i]])], col = cala[i], lwd = 1, lty = 2)
  lines(PPSIZE[[i]][!is.nan(CCOEVARsd[[i]])], CCOEVAR[[i]][!is.nan(CCOEVARsd[[i]])]+2*CCOEVARsd[[i]][!is.nan(CCOEVARsd[[i]])], col = cala[i], lwd = 1, lty = 2)
  polygon(c(PPSIZE[[i]][!is.na(CCOEVARsd[[i]])],rev(PPSIZE[[i]][!is.na(CCOEVARsd[[i]])])),c(CCOEVAR[[i]][!is.na(CCOEVARsd[[i]])]-2*CCOEVARsd[[i]][!is.na(CCOEVARsd[[i]])],rev(CCOEVAR[[i]][!is.na(CCOEVARsd[[i]])]+2*CCOEVARsd[[i]][!is.na(CCOEVARsd[[i]])])), border = NA, col = adjustcolor(cala[i], alpha.f = 0.5) )
  
}

arrows(350, -0.03, 0, -0.03, xpd = TRUE)
legend("topright", legend = c( "Case 2", "Case 3", "Control"), lwd = 3, col = cala[-1], text.font = 4, pch = 1)

text(250,0.070, "N > K", cex = 2.0, srt = 360, font = 3)

# Std deviation
i = 1
plot(PPSIZE[[i]][!is.nan(STDVARsd[[i]])], STDVAR[[i]][!is.nan(STDVARsd[[i]])], type = 'n', col = cala[i], lwd = 3, ylim = c(0.008,0.26), xlim = c(0,350), xlab = list("Population size", font = 4, col = 1, cex = 1.5), ylab = list("Standard deviation", font = 4, col = 1, cex = 1.5))
rect(200, -0.024, 400, 10.28 , density = -0.1 , angle = 45, col = "gray", border = NULL, lty = par("lty"), lwd = par("lwd"))

for (i in 2:dim(Pp)[3]){
  lines(PPSIZE[[i]][!is.nan(STDVARsd[[i]])], STDVAR[[i]][!is.nan(STDVARsd[[i]])], type = 'o', col = cala[i], lwd = 3)
  lines(PPSIZE[[i]][!is.nan(STDVARsd[[i]])], STDVAR[[i]][!is.nan(STDVARsd[[i]])]-2*STDVARsd[[i]][!is.nan(STDVARsd[[i]])], col = cala[i], lwd = 1, lty = 2)
  lines(PPSIZE[[i]][!is.nan(STDVARsd[[i]])], STDVAR[[i]][!is.nan(STDVARsd[[i]])]+2*STDVARsd[[i]][!is.nan(STDVARsd[[i]])], col = cala[i], lwd = 1, lty = 2)
  polygon(c(PPSIZE[[i]][!is.na(STDVARsd[[i]])],rev(PPSIZE[[i]][!is.na(STDVARsd[[i]])])),c(STDVAR[[i]][!is.na(STDVARsd[[i]])]-2*STDVARsd[[i]][!is.na(STDVARsd[[i]])],rev(STDVAR[[i]][!is.na(STDVARsd[[i]])]+2*STDVARsd[[i]][!is.na(STDVARsd[[i]])])), border = NA, col = adjustcolor(cala[i], alpha.f = 0.5) )
  
}

arrows(350, -0.033, 0, -0.033, xpd = TRUE)
text(250,0.12, "N > K", cex = 2.0, srt = 360, font = 3)
legend("topright", legend = c( "Case 2", "Case 3", "Control"), lwd = 3, col = cala[-1], text.font = 4, pch = 1)

# Skewness
i = 1
plot(PPSIZE[[i]][!is.nan(SKNESSsd[[i]])], SKNESS[[i]][!is.nan(SKNESSsd[[i]])], type = 'n', col = cala[i], lwd = 3, ylim = c(-2.4,0.3), xlim = c(0,300), xlab = list("Population size", font = 4, col = 1, cex = 1.5), ylab = list("Skewness", font = 4, col = 1, cex = 1.5))
rect(200, -12.8, 400, 10.1 , density = -0.01 , angle = 45, col = "gray", border = NULL, lty = par("lty"), lwd = par("lwd"))
for (i in 2:(dim(Pp)[3]-1)){
  lines(PPSIZE[[i]][!is.nan(SKNESSsd[[i]])], SKNESS[[i]][!is.nan(SKNESSsd[[i]])], type = 'o', col = cala[i], lwd = 3)
  lines(PPSIZE[[i]][!is.nan(SKNESSsd[[i]])], SKNESS[[i]][!is.nan(SKNESSsd[[i]])]-2*SKNESSsd[[i]][!is.nan(SKNESSsd[[i]])], col = cala[i], lwd = 1, lty = 2)
  lines(PPSIZE[[i]][!is.nan(SKNESSsd[[i]])], SKNESS[[i]][!is.nan(SKNESSsd[[i]])]+2*SKNESSsd[[i]][!is.nan(SKNESSsd[[i]])], col = cala[i], lwd = 1, lty = 2)
  polygon(c(PPSIZE[[i]][!is.na(SKNESSsd[[i]])],rev(PPSIZE[[i]][!is.na(SKNESSsd[[i]])])),c(SKNESS[[i]][!is.na(SKNESSsd[[i]])]-2*SKNESSsd[[i]][!is.na(SKNESSsd[[i]])],rev(SKNESS[[i]][!is.na(SKNESSsd[[i]])]+2*SKNESSsd[[i]][!is.na(SKNESSsd[[i]])])), border = NA, col = adjustcolor(cala[i], alpha.f = 0.5) )
  
}


arrows(350, -2.8, 0, -2.8, xpd = TRUE)
text(250,0.15, "N > K", cex = 2.0, srt = 360, font = 3)

i = 4
XXX = cbind(PPSIZE[[i]][!is.nan(SKNESSsd[[i]])], SKNESS[[i]][!is.nan(SKNESSsd[[i]])])
sbp = subplot(plot(XXX, col=cala[i], type = 'o', pch='.', xlab='', ylab='', cex.axis=1.3, lwd = 3, ylim = c(0.2,0.9)), x = 120, y = -2.2, size=c(3,3), vadj=0.0, hadj=0.0 )

par(sbp)
lines(XXX, col=cala[i], type = 'o', lwd = 3)
polygon(c(PPSIZE[[i]][!is.nan(SKNESSsd[[i]])],rev(PPSIZE[[i]][!is.nan(SKNESSsd[[i]])])),c(SKNESS[[i]][!is.nan(SKNESSsd[[i]])]-2*SKNESSsd[[i]][!is.nan(SKNESSsd[[i]])],rev(SKNESS[[i]][!is.nan(SKNESSsd[[i]])]+2*SKNESSsd[[i]][!is.nan(SKNESSsd[[i]])])), border = NA, col = adjustcolor(cala[i], alpha.f = 0.5) )


# Kurtosis
i = 1
plot(PPSIZE[[i]][!is.nan(KTSSsd[[i]])], KTSS[[i]][!is.nan(KTSSsd[[i]])], type = 'n', col = cala[i], lwd = 3, ylim = c(1.50,8.0),xlim = c(0,350), xlab = list("Population size", font = 4, col = 1, cex = 1.5), ylab = list("Kurtosis", font = 4, col = 1, cex = 1.5))
rect(200, -2, 400, 20 , density = -0.01 , angle = 45,  col = "gray", border = NULL, lty = par("lty"), lwd = par("lwd"))

for (i in 2:dim(Pp)[3]){
  lines(PPSIZE[[i]][!is.nan(KTSSsd[[i]])], KTSS[[i]][!is.nan(KTSSsd[[i]])], type = 'o', col = cala[i], lwd = 3)
  lines(PPSIZE[[i]][!is.nan(KTSSsd[[i]])], KTSS[[i]][!is.nan(KTSSsd[[i]])]-2*KTSSsd[[i]][!is.nan(KTSSsd[[i]])], col = cala[i], lwd = 1, lty = 2)
  lines(PPSIZE[[i]][!is.nan(KTSSsd[[i]])], KTSS[[i]][!is.nan(KTSSsd[[i]])]+2*KTSSsd[[i]][!is.nan(KTSSsd[[i]])], col = cala[i], lwd = 1, lty = 2)
  polygon(c(PPSIZE[[i]][!is.na(KTSSsd[[i]])],rev(PPSIZE[[i]][!is.na(KTSSsd[[i]])])),c(KTSS[[i]][!is.na(KTSSsd[[i]])]-2*KTSSsd[[i]][!is.na(KTSSsd[[i]])],rev(KTSS[[i]][!is.na(KTSSsd[[i]])]+2*KTSSsd[[i]][!is.na(KTSSsd[[i]])])), border = NA, col = adjustcolor(cala[i], alpha.f = 0.5) )
  
}

arrows(350, 0.4, 0, 0.4, xpd = TRUE)
text(250,7, "N > K", cex = 2.0, srt = 360, font = 3)


#Auto-correlation at lag 1
i = 1
plot(PPSIZE[[i]][!is.nan(AUTOsd[[i]])], AUTO[[i]][!is.nan(AUTOsd[[i]])], type = 'n', col = cala[i], lwd = 3,xlim = c(0,350), ylim = c(-0.15,0.92), xlab = list("Population size", font = 4, col = 1, cex = 1.5), ylab = list("Auto-correlation", font = 4, col = 1, cex = 1.5))
rect(200, -10.4, 400, 10.9 , density = -0.01 , angle = 45, col = "gray", border = NULL, lty = par("lty"), lwd = par("lwd"))

for (i in 2:dim(Pp)[3]){
  lines(PPSIZE[[i]][!is.nan(AUTOsd[[i]])], AUTO[[i]][!is.nan(AUTOsd[[i]])], type = 'o', col = cala[i], lwd = 3)
  lines(PPSIZE[[i]][!is.nan(AUTOsd[[i]])], AUTO[[i]][!is.nan(AUTOsd[[i]])]-2*AUTOsd[[i]][!is.nan(AUTOsd[[i]])], col = cala[i], lwd = 1, lty = 2)
  lines(PPSIZE[[i]][!is.nan(AUTOsd[[i]])], AUTO[[i]][!is.nan(AUTOsd[[i]])]+2*AUTOsd[[i]][!is.nan(AUTOsd[[i]])], col = cala[i], lwd = 1, lty = 2)
  polygon(c(PPSIZE[[i]][!is.na(AUTOsd[[i]])],rev(PPSIZE[[i]][!is.na(AUTOsd[[i]])])),c(AUTO[[i]][!is.na(AUTOsd[[i]])]-2*AUTOsd[[i]][!is.na(AUTOsd[[i]])],rev(AUTO[[i]][!is.na(AUTOsd[[i]])]+2*AUTOsd[[i]][!is.na(AUTOsd[[i]])])), border = NA, col = adjustcolor(cala[i], alpha.f = 0.5) )
  
}

arrows(350, -0.3, 0, -0.3, xpd = TRUE)
text(250,0.6, "N > K", cex = 2.0, srt = 360, font = 3)

#======== End of plots for early-warning signals vs population size ==============================

#=============== End of Analysis ==================================================================
```
