## Supplementary material for "Models for Eco-evolutionary Extinction Vortices and their Detection": R scripts for regenerating data used in the manuscript: IndividualBasedModelSubmit.R

```
#==== This script simulates Geometric population growth model and Ricker model. Also, it simulates heterozygote advantage and fluctuating selection with reversal of dominance

library("doMC")
cores=detectCores()
cl <- makeCluster(cores) 
registerDoMC(cores)
require(reshape2)

#function for Ricker model
ricker <- function(Rate,N,K){
  EX <- exp(Rate*(1-N/K))
  return(EX)
}

# This function calculates for number of polymorphic loci and population size

FITNESS_ALLOCATION<-function( allalleles_mat, Fitness){
  if(nrow(allalleles_mat)>0){
    refma<-cbind(rep(1:nrow(allalleles_mat),each=numberofloci),seq(1,numberofloci*2,2))#do before function so we do not repeat uneccessary!
    refmb<-cbind(rep(1:nrow(allalleles_mat),each=numberofloci),seq(2,numberofloci*2,2))
    #setting up locus descriptions
  ma<-matrix(paste(allalleles_mat[refma],allalleles_mat[refmb],sep=""), ncol=numberofloci, nrow = nrow(allalleles_mat), byrow = T) #pasting together so we have AA, aa, Aa, and aA ||| rows are individualsalleles, collumn are 
  
  #calculating fitness
  ma1<-t(apply(ma,1,function(x){rbind(sum(x=="AA"),sum(x=="aa"),sum(x=="Aa"|x=="aA"))})) # transfering it to the number of aa, AA, and aA/Aa Loci ||| collumns are individuals row1==AA,row2==aa,row3==aA/Aa
  colnames(ma1)<-c("AA","aa","Aa/aA")
  indifit<-apply(t(Fitness^t(ma1)),1,prod) #transfering to fitness (multiplicative) per inidiviudal
  popfit <- mean(indifit)
  #finding polymorphism at population level (polymorphism== which loci are still polymorphic, ma2==how polymorphic are the loci)
 ##polymorphicloci<-numberofloci-sum(apply(ma1==0,1,function(x){sum(x==c(F,T,T))==3|sum(x==c(T,F,T))==3}))# number of poly loci at population level #TRUE means non polymorphic!!
 # propor_aa = sum(ma1[,"aa"])/sum(ma1)
  propor_AA = sum(ma1[,"AA"])/sum(ma1)
  propor_Aa = sum(ma1[,"Aa/aA"])/sum(ma1)
  pop_freq_A<-(sum(ma1[,1])*2+sum(ma1[,3])) /(sum(ma1)*2) #proportion of loci being A
  
  
  ma2 <- apply(ma,2,function(x){rbind(sum(x=="AA"),sum(x=="aa"),sum(x=="Aa"|x=="aA"))})
  rownames(ma2)<-c("AA","aa","Aa/aA")
  degreeloci_freq_a = (ma2[2,]*2+ma2[3,]) /(colSums(ma2)*2)
  degreeloci_freq_A = (ma2[1,]*2+ma2[3,]) /(colSums(ma2)*2)
  
  popsize = nrow(allalleles_mat)

 
  wa = which(degreeloci_freq_a>0); wA = which(degreeloci_freq_A>0); WW = c(wa,wA)
  polymorphicloci = length(WW)-length(unique(WW))#
  #-------------------------------------------------
    
  return(list(popsize, polymorphicloci))
  }else{
  polymorphicloci = 0
  popsize = 0
  return(list(popsize, polymorphicloci))
    
}}

# This function returns a matrix for offspring details every generation. 
BABYMAT = function(allalleles_mat, mut_a, mut_A, deathrate, Rate, Geometric){

N = nrow(allalleles_mat) # number of parents

reproducers = 1-rbinom(N,1,deathrate) # This applies death of parents. Currently set to 0.
allalleles_reproducer = allalleles_mat[(reproducers==1),,drop = FALSE]
tot_reproducers = sum(reproducers) # Total number of parents for reproduction

if (Geometric == 1){
  if ( N>=K){
    tot_offspring = rpois(1,tot_reproducers*ricker(Rate,N,K))
  }else{
    tot_offspring = rpois(1,tot_reproducers*exp(Rate))
  }
}else{
tot_offspring = rpois(1,tot_reproducers*ricker(Rate,N,K))
}

if(tot_offspring>0){
  
mother_index = sample(tot_reproducers,tot_offspring,replace = TRUE)#1:tot_reproducers
father_index = sample(tot_reproducers,tot_offspring,replace = TRUE)

# parent genes
fathergenes = allalleles_reproducer[father_index,,drop = FALSE]
mothergenes =  allalleles_reproducer[mother_index,,drop = FALSE]

#rearrangement of genes
genesmother<-matrix(c(mothergenes[,seq(1,ncol(mothergenes),2)],mothergenes[,seq(2,ncol(mothergenes),2)]),ncol = 2, byrow = FALSE)
genesfather<-matrix(c(fathergenes[,seq(1,ncol(fathergenes),2)],fathergenes[,seq(2,ncol(fathergenes),2)]), ncol = 2, byrow = FALSE)


frompa = apply(genesfather, 1, function(x) {sample(x,1,replace = T)})
fromma = apply(genesmother, 1, function(x) {sample(x,1,replace = T)})
genesbaby = matrix(0, nrow = tot_offspring, ncol = 2*numberofloci)
xx = seq(1,ncol(genesbaby), 2); yy = seq(2, ncol(genesbaby),2)
genesbaby[,xx] = fromma
genesbaby[,yy] = frompa

# mutation occurs here
index_a = which(genesbaby=="a")
index_A = which(genesbaby=="A")

MUTS_a = which(runif(length(which(genesbaby=="a")))<=mut_a) #selecting places where mutation happens for a
MUTS_A = which(runif(length(which(genesbaby=="A")))<=mut_A) #selecting places where mutation happens for A
if (length(MUTS_a)>0){
  genesbaby[index_a[MUTS_a]] = "A"
}
if (length(MUTS_A)>0){
  genesbaby[index_A[MUTS_A]] = "a"
}

return(genesbaby)
}
}

# This function returns the fitness of each individual
OFFSPRING_FITNESS <- function( allalleles_mat, Fitness){
  if(nrow(allalleles_mat)>0){
    refma<-cbind(rep(1:nrow(allalleles_mat),each=numberofloci),seq(1,numberofloci*2,2))#do before function so we do not repeat uneccessary!
    refmb<-cbind(rep(1:nrow(allalleles_mat),each=numberofloci),seq(2,numberofloci*2,2))
    #setting up locus descriptions
    ma<-matrix(paste(allalleles_mat[refma],allalleles_mat[refmb],sep=""), ncol=numberofloci, nrow = nrow(allalleles_mat), byrow = T) #pasting together so we have AA, aa, Aa, and aA ||| rows are individualsalleles, collumn are 
    
    #calculating fitness
    ma1<-t(apply(ma,1,function(x){rbind(sum(x=="AA"),sum(x=="aa"),sum(x=="Aa"|x=="aA"))})) # transfering it to the number of aa, AA, and aA/Aa Loci ||| collumns are individuals row1==AA,row2==aa,row3==aA/Aa
    colnames(ma1)<-c("AA","aa","Aa/aA")
    indifit<-apply(t(Fitness^t(ma1)),1,prod) #transfering to fitness (multiplicative) per inidiviudal
    
    return(indifit)
  }else{
    return(-1)
  }
  
  }


# This is the individual-based model that calls all the above functions.
IBM <- function(founderpop, numberofloci, K, mut_A, mut_a,  replicates, tmax, p0, smaxs,Rate,deathrate, smaxt, ConstantSelection, Geometric, percent) {
  
  foreach(sim = 1:replicates, .combine = cbind)%dopar%{
    
    populationsize <- rep(0,tmax)# Population size
    NumberLociPoly = rep(0, tmax)# Number of polymorphic loci
    
    noMate = 0
    
    alleles <- c("a", "A") # alleles present at each locus
    
    # Initial fitness of genotypes
    Fit_AA = 1
    Fit_Aa = 1
    Fit_aa = 1
    
    Fitness_init = c(Fit_AA, Fit_aa, Fit_Aa)
    
    inpop = rpois(1,founderpop)# Starting pop
    allalleles <- matrix(sample(alleles, inpop * 2* numberofloci,replace=TRUE, prob = c(p0, 1-p0)),ncol=2*numberofloci)#allels to be distributed among individuals
    
    # Initial parameters from the initial population "al"
    Output1 = FITNESS_ALLOCATION(allalleles_mat = allalleles, Fitness = Fitness_init)
    populationsize[1] <- Output1[[1]]
    NumberLociPoly[1] =  Output1[[2]]
    
      Ti = 1:tmax
      s <- smaxs*sin(2*pi*Ti/50 + pi) # selection coeff for genotype AA
      te <- smaxs*sin(2*pi*Ti/50)
            
      hh1 = 0.5-(percent/100)*sin(2*pi*Ti/50)
      
      
      Ft_AA = (1-te)/(1+smaxs) 
      Ft_aa = (1-s)/(1+smaxs)
      Ft_Aa =  (1-hh1*te-(1-hh1)*s)/(1+smaxs) 


     t = 2
    
    # looping through generations as long as population size is above 0
    while (t <= tmax & populationsize[t-1] > 0){
      #fitness of genotypes
     if (ConstantSelection==1){

       cFit_AA = 1-smaxt 
       cFit_aa = 1-smaxs
       cFit_Aa = 1
       Fitness = c(cFit_AA, cFit_aa, cFit_Aa)

      }else{
     
     Fitness = c(Ft_AA[t-1], Ft_aa[t-1], Ft_Aa[t-1])
      }
      
     N = nrow(allalleles)
     
      if (N > 0){
        #let all the individuals reproduce
        
        offspringmat = BABYMAT(allalleles_mat = allalleles, mut_a = mut_a, mut_A = mut_A, deathrate = deathrate, Rate = Rate, Geometric = Geometric)
        
        
        if (is.numeric(nrow(offspringmat))){
        offspringfit = OFFSPRING_FITNESS(allalleles_mat = offspringmat, Fitness = Fitness)
        
        Viable = which(runif(nrow(offspringmat))<=offspringfit)
        
        if (length(Viable)>0){
        allalleles = offspringmat[Viable, ,drop =FALSE]
        
        # updating the measurements where viable offspring replace parents
        
        Output2 = FITNESS_ALLOCATION(allalleles_mat = allalleles, Fitness = Fitness)
        populationsize[t] <- Output2[[1]]
        NumberLociPoly[t] =  Output2[[2]]
         }
        }else{
          populationsize[t] <- 0
          NumberLociPoly[t] =  0
          
        }
      }
      
      t = t + 1
    }
    return(list(populationsize, NumberLociPoly ))
  }
}

# Here one can switch on specific growth model and selection mechansim. It is advised to keep this constant and vary either growth rate or selection coefficient for easy analysis using our provided code
#fixed parameters throught the manuscript
replicates = 100 # Number of replicates
numberofloci <- 100 # Number of loci
d0 = 0.0 # death rate. In this study, it is set to zero. No death rate considered
p = 0.5 # initial frequency of allele "A" in the population
percent = 20 # Parameter used during fluctuating selection scenario. It is denoted as c in the methods. Here it is expressed as a percentage.
mut_a = 0 # mutation rate for allele a. Set to 0 throught.
mut_A = 0# mutation rate for allele A. Set to 0 throught.
kapa = 50 #

#Varrying parameters, assuming the parameters below where used  
geometric = 1 # 1 selects for geometric growth rate other values select for Ricker model
constantselection = 1 # One selects for heterozygote advantage. Other values select for fluctuating selction with reversal of dominance. 
tmax <- 3000 # Time for running simulations. This worked very well for N0 = K = 200 but 30000 worked well for N0 = K = 2000. The code stops when either population is extinct or this time is reached.
maxsels = 0.005 # selection coefficient
maxselt = maxsels
Founderpop <- 200 # starting population size
K <- Founderpop # Founder population size denoted as N_0 in the text used as carrying capacity
rate = log(c(1.2, 1.5, 1.8)) # Intrinsic growth rate. In the main text, it is denoted as r. Note that you can run one case at a time

# Actual simulation. My simulated data is saved as an array in 3-Dimension.  

#PopulationArray = array(0,dim = c(tmax, replicates, length(rate)))# Array of population sizes in 3 dimension. It includes all cases considered only when varrying one variable. Here I varry rate.
#PolymorphicLociArray = array(0,dim = c(tmax, replicates, length(rate)))# Array of polymorphic loci in 3 dimension. It includes all cases considered only when varrying one variable. Here I varry rate.

for (case in 1:length(rate)){ # assuming growth rate is a vector of values
  
SIMU <- IBM(K = K, founderpop = Founderpop, mut_A = mut_A, mut_a = mut_a, tmax = tmax, Rate = rate, deathrate = d0, replicates = replicates, p0= p, percent = percent, smaxs = maxsels, smaxt = maxselt, numberofloci = numberofloci, ConstantSelection = constantselection, Geometric = geometric)

PopHeteroLoci = matrix(0, nrow = tmax, ncol = replicates) # polymorphic loci
POPSIZE = matrix(0, nrow = tmax, ncol = replicates) # population size

for (s in 1:replicates){
  POPSIZE[,s] = SIMU[1,][[s]]
  PopHeteroLoci[,s] = unlist(SIMU[2,][[s]])
  
}

# Note that here, the file name is generated for the respective parameter combination.
# We assume for this specific case, rate is varrying and other parameters are kept constant and so the indexing "case" should be on rate.
#If another parameter is varrying, the indexing should change accordingly right from the for loop for cases.

save(POPSIZE, file = paste("POPSIZE_r0_log",exp(rate[case]), "_N0_", Founderpop, "_selA_sela_", maxsels, "_mut_", mut_a, "_rep_",replicates, "_KN0_p_", p,"_", numberofloci, "loci", "_ConstantSelection_", constantselection, "_Geometric_",geometric, ".RData", sep = ""))   
save(PopHeteroLoci, file = paste("POLYLOCI_r0_log",exp(rate[case]), "_N0_", Founderpop, "_selA_sela_", maxsels, "_mut_", mut_a, "_rep_",replicates, "_KN0_p_", p,"_", numberofloci, "loci", "_ConstantSelection_", constantselection, "_Geometric_",geometric, ".RData", sep = ""))   

}

# ======= After simulating the data for population sizes and polymorphic loci for the different parameter combinations, combine it into an array of 3-dimension for easy analysis using our code provided.
```
