## Supplementary material for "Models for Eco-evolutionary Extinction Vortices and their Detection": R scripts for regenerating data used in the manuscript: NumericalApproximationSubmit.R

```
#++++++++++++++++++++++++++++++++++++++++++++++++++++++++++++++++++
#==================Complete scheme with M(x,t) evaluated as in the paper for Complete numerical solution====================================
# This function evaluates the numerical scheme described. 

AnalyticApprox_Numeric = function(HeterozygoteAdva, geometric, x0, H0, N0, Capacity, rate, tmax, se, tau, KK, kapa){
  
  h = 1/KK # spacing over [0,1]
  tt = seq(1,tmax,tau) # time steps
  ii = 1:(KK-1) # inner grid for (0,1)
  xx_plus = (ii + 1)*h  
  xx_minus = (ii - 1)*h
  xx_1 = (1)*h
  xx_Mminus1 = (KK-1)*h 
  
  xx = c(0, 0.5*(xx_plus+xx_minus), 1)
  
  het = numeric(length(tt)) # stores heterozygosity
  het[1] = 1#x0 # initial heterozygosity
  
  Ht =numeric(length(tt)) # stores number of polymorphic loci
  Ht[1] = H0
  Ne = numeric(length(tt) + 1) # Population size
  Ne[1] = N0
  
  ff = matrix(0, nrow = c(KK+1), ncol = length(tt)) # stores density for each time step
  ff[median(0:KK+1),1] = 1/h # start with a dirac function represented by a pick at initial frequency
  #++++++++++++++++++++
  
  if(HeterozygoteAdva == 1){
    Ww = function(fq){
      (2*fq*se*(1-fq) + (1-se))
      
    }
    WW = exp(H0*((h/2)*ff[1,1]*log(Ww(xx[1])) + (h/2)*ff[KK+1,1]*log(Ww(xx[KK+1])) + sum(h*ff[-c(1,KK+1),1]*log(Ww(xx[2:KK]))) ))  #fitness
    #------------------------------------------------------------------
    
    #calculating Ne at timestep 2 because in the iteration, we use population size at t+1 to get density at t+1
    if (geometric == 1){
      if (Ne[1]<Capacity){
        Ne[2] = WW*Ne[1]*exp(rate)
      }else{
        Ne[2] = WW*Ne[1]*exp(rate*(1-Ne[1]/Capacity))
      }}else{
        Ne[2] = WW*Ne[1]*exp(rate*(1-Ne[1]/Capacity))
      }
    #++++++++++++++++++++++++
    
    
    for (ti in 1:(length(tt)-1)){
      
      M_minus = se*xx_minus*(1-xx_minus)*(1-2*xx_minus)
      M_plus = se*xx_plus*(1-xx_plus)*(1-2*xx_plus)
      
      V_iplus1 = xx_plus*(1-xx_plus)/(4*Ne[ti+1])
      V_iminus1 = xx_minus*(1-xx_minus)/(4*Ne[ti+1])
      V_i = (ii)*h*(1-(ii)*h)/(4*Ne[ti+1])
      
      
      M_1 = se*xx_1*(1-xx_1)*(1-2*xx_1)
      V_1 = xx_1*(1-xx_1)/(4*Ne[ti+1])
      
      M_Mminus1 = se*xx_Mminus1*(1-xx_Mminus1)*(1-2*xx_Mminus1)
      V_Mminus1 = xx_Mminus1*(1-xx_Mminus1)/(4*Ne[ti+1])
      
      #Elements of the matrix
      bita = -(V_iminus1/(h^2) + M_minus/(2*h))
      gama = (2*V_i/(h^2) + 1/tau)
      alpha = M_plus/(2*h) - V_iplus1/(h^2)
      
      #boundary x = 0
      rho = M_1/h - 2*V_1/(h^2)
      epsilon = 1/tau 
      #boundary x = 1
      phi = 1/tau
      sigma = -(M_Mminus1/h + 2*V_Mminus1/(h^2))
      right = 1/tau
      
      # filling the coefficient matrix M on LHS
      Coemat = matrix(0,nrow = (KK+1), ncol = (KK+1))
      Coemat[1,1] = epsilon
      Coemat[1,2] = rho
      Coemat[KK+1, KK] = sigma
      Coemat[KK+1, KK+1] = phi
      
      for (i in 2:KK){
        Coemat[i, i] = gama[i-1]
        Coemat[i,i-1] = bita[i-1]
        Coemat[i, i+1] = alpha[i-1]
      }
      
      # Matrix on RHS
      Rightmat = matrix(0, nrow = (KK+1), ncol = (KK+1))
      diag(Rightmat) = right
      
      # Updating values of interest
      ff[,ti+1] = solve(Coemat)%*%Rightmat%*%ff[,ti]
      het[ti+1] = 1*(KK-1)*mean(h*ff[-c(1,KK+1),ti])
      Ht[ti+1] =  Ht[ti]*(1 - (het[ti] - het[ti+1])/het[ti])
      
      pq2 = 0.5*(KK-1)*mean(h*ff[-c(1,KK+1),ti])
      frq = (1-sqrt(abs(1-2*pq2)))/2
      # WW = (2*frq*se*(1-frq) + (1-se))^Ht[ti] * (1-se)^(H0-Ht[ti])   #exp(Ht[1]*log(2*frq*se*(1-frq) + (1-se)))
      #---------------------------------------------------------------
      WW = exp(H0*((h/2)*ff[1,ti]*log(Ww(xx[1])) + (h/2)*ff[KK+1,ti]*log(Ww(xx[KK+1])) + sum(h*ff[-c(1,KK+1),ti]*log(Ww(xx[2:KK]))) ))  #fitness
      
      #------------------------------------------------------------------
      
      if (geometric == 1){
        if (Ne[ti+1]<Capacity){
          Ne[ti+2] = WW*Ne[ti+1]*exp(rate)
        }else{
          Ne[ti+2] = WW*Ne[ti+1]*exp(rate*(1-Ne[ti+1]/Capacity))
        }}else{
          Ne[ti+2] = WW*Ne[ti+1]*exp(rate*(1-Ne[ti+1]/Capacity))
        }
      
    } 
  }else{
    
    #======== Fluctuating selection with reversal of dominance
    #==================Complete scheme with M(x,t) evaluated as in the paper for Complete numerical solution Fluctuating selection ============================
    
    percent = 20
    sA <- se*sin(2*pi*tt/kapa) # selection coeff for genotype AA
    sa <- se*sin(2*pi*tt/kapa + pi)#smaxt*cos(Ti)^2# selection coeff for aa
    hh1 = 0.5-(percent/100)*sin(2*pi*tt/kapa)#(smaxs - smaxs*percent/100)*cos(Ti)^10
    
    #=============
    
    Ww = function(fq){
      tii = 1
      (1-sa[tii] + fq*(sa[tii]-sA[tii])*(fq+2*hh1[tii]-2*fq*hh1[tii]))/(1+se)
      
    }
    WW = exp(H0*((h/2)*ff[1,1]*log(Ww(xx[1])) + (h/2)*ff[KK+1,1]*log(Ww(xx[KK+1])) + sum(h*ff[-c(1,KK+1),1]*log(Ww(xx[2:KK]))) ))  #fitness
    #------------------------------------------------------------------
    
    #calculating Ne at timestep 2 because in the iteration, we use population size at t+1 to get density at t+1
    if (geometric == 1){
      if (Ne[1]<Capacity){
        Ne[2] = WW*Ne[1]*exp(rate)
      }else{
        Ne[2] = WW*Ne[1]*exp(rate*(1-Ne[1]/Capacity))
      }}else{
        Ne[2] = WW*Ne[1]*exp(rate*(1-Ne[1]/Capacity))
      }
    #++++++++++++++++++++++++
    
    
    for (ti in 1:(length(tt)-1)){
      #++++++++++++++++++
      M_minus = xx_minus*(1-xx_minus)*(xx_minus*(1-2*hh1[ti+1])+hh1[ti+1])*(1+sa[ti+1])*(sa[ti+1]-sA[ti+1]) - xx_minus^2*(1-xx_minus)*(sa[ti+1]-sA[ti+1])^2*(xx_minus*(1-2*hh1[ti+1])+hh1[ti+1])*(xx_minus*(1-2*hh1[ti+1])+2*hh1[ti+1])
      M_plus = xx_plus*(1-xx_plus)*(xx_plus*(1-2*hh1[ti+1])+hh1[ti+1])*(1+sa[ti+1])*(sa[ti+1]-sA[ti+1]) - xx_plus^2*(1-xx_plus)*(sa[ti+1]-sA[ti+1])^2*(xx_plus*(1-2*hh1[ti+1])+hh1[ti+1])*(xx_plus*(1-2*hh1[ti+1])+2*hh1[ti+1])
      #+++++++++++++++++++
      
      
      
      V_iplus1 = xx_plus*(1-xx_plus)/(4*Ne[ti+1])
      V_iminus1 = xx_minus*(1-xx_minus)/(4*Ne[ti+1])
      V_i = (ii)*h*(1-(ii)*h)/(4*Ne[ti+1])
      
      
      M_1 = xx_1*(1-xx_1)*(xx_1*(1-2*hh1[ti+1])+hh1[ti+1])*(1+sa[ti+1])*(sa[ti+1]-sA[ti+1]) - xx_1^2*(1-xx_1)*(sa[ti+1]-sA[ti+1])^2*(xx_1*(1-2*hh1[ti+1])+hh1[ti+1])*(xx_1*(1-2*hh1[ti+1])+2*hh1[ti+1])
      V_1 = xx_1*(1-xx_1)/(4*Ne[ti+1])
      
      M_Mminus1 = xx_Mminus1*(1-xx_Mminus1)*(xx_Mminus1*(1-2*hh1[ti+1])+hh1[ti+1])*(1+sa[ti+1])*(sa[ti+1]-sA[ti+1]) - xx_Mminus1^2*(1-xx_Mminus1)*(sa[ti+1]-sA[ti+1])^2*(xx_Mminus1*(1-2*hh1[ti+1])+hh1[ti+1])*(xx_Mminus1*(1-2*hh1[ti+1])+2*hh1[ti+1])
      V_Mminus1 = xx_Mminus1*(1-xx_Mminus1)/(4*Ne[ti+1])
      
      #Elements of the matrix
      bita = -(V_iminus1/(h^2) + M_minus/(2*h))
      gama = (2*V_i/(h^2) + 1/tau)
      alpha = M_plus/(2*h) - V_iplus1/(h^2)
      
      #boundary x = 0
      rho = M_1/h - 2*V_1/(h^2)
      epsilon = 1/tau 
      #boundary x = 1
      phi = 1/tau
      sigma = -(M_Mminus1/h + 2*V_Mminus1/(h^2))
      right = 1/tau
      
      # filling the coefficient matrix M on LHS
      Coemat = matrix(0,nrow = (KK+1), ncol = (KK+1))
      Coemat[1,1] = epsilon
      Coemat[1,2] = rho
      Coemat[KK+1, KK] = sigma
      Coemat[KK+1, KK+1] = phi
      
      for (i in 2:KK){
        Coemat[i, i] = gama[i-1]
        Coemat[i,i-1] = bita[i-1]
        Coemat[i, i+1] = alpha[i-1]
      }
      
      # Matrix on RHS
      Rightmat = matrix(0, nrow = (KK+1), ncol = (KK+1))
      diag(Rightmat) = right
      
      # Updating values of interest
      ff[,ti+1] = solve(Coemat)%*%Rightmat%*%ff[,ti]
      het[ti+1] = 1*(KK-1)*mean(h*ff[-c(1,KK+1),ti])
      Ht[ti+1] =  Ht[ti]*(1 - (het[ti] - het[ti+1])/het[ti])
      
      pq2 = 0.5*(KK-1)*mean(h*ff[-c(1,KK+1),ti])
      frq = (1-sqrt(abs(1-2*pq2)))/2
      
      #---------------------------------------------------------------
      Ww = function(fq){
        tii = ti+1
        (1-sa[tii] + fq*(sa[tii]-sA[tii])*(fq+2*hh1[tii]-2*fq*hh1[tii]))/(1+se)
        
      }
      WW = exp(H0*((h/2)*ff[1,ti]*log(Ww(xx[1])) + (h/2)*ff[KK+1,ti]*log(Ww(xx[KK+1])) + sum(h*ff[-c(1,KK+1),ti]*log(Ww(xx[2:KK]))) ))  #fitness
      
      #------------------------------------------------------------------
      
      if (geometric == 1){
        if (Ne[ti+1] < Capacity){
          Ne[ti+2] = WW*Ne[ti+1]*exp(rate)
        }else{
          Ne[ti+2] = WW*Ne[ti+1]*exp(rate*(1-Ne[ti+1]/Capacity))
        }}else{
          Ne[ti+2] = WW*Ne[ti+1]*exp(rate*(1-Ne[ti+1]/Capacity))
        }
      
    }
  }
  return(list(Ne, Ht,ff, xx, tt))
}


HeterozygoteAdva = 1 # value of 1 selects for heterozygote advantage otherwise it is for fluctuating selction with reversal of dominance
geometric = 1# value 1 implies geometric otherwise Ricker
x0 = 0.5# initial allele frequency
H0 = 100# number of loci
N0 = 200# Initial population size
Capacity = N0 #Carrying capacity
rate = log(1.6160) # per-capita growth rate r_0
tmax = 1000 # time to run the simulations
se = 0.005 # selection coefficient
tau = 1 # time spacing for grid points
KK = 10# Number of grid points for allele frequency grid
kapa = 50 # Number of time-steps in a complete cycle for fluctuating selection with dominance reversals

ANA_APPROX_NUM = AnalyticApprox_Numeric(HeterozygoteAdva, geometric, x0, H0, N0, Capacity, rate, tmax, se, tau, KK, kapa)

# Extracting data
NE = ANA_APPROX_NUM[[1]]
HT = ANA_APPROX_NUM[[2]]
Dens = ANA_APPROX_NUM[[3]]
freqx = ANA_APPROX_NUM[[4]]
TT = ANA_APPROX_NUM[[5]]
#Some plots ==================
TTb = 3000# Time axis limit

# Population size vs time
plot(TT, NE[-(length(TT)+1)], type = 'l', xlim = c(0,TTb))
lines(TT, NE[-(length(TT)+1)], lwd = 3, col = 5)

# Polymorphic loci vs time
plot(TT, HT, type = 'l', xlim = c(0,TTb))
lines(TT, HT, lwd = 3, col = 5)

#==============================================
# Plots for some densities
plot(freqx, Dens[,100]/sum(Dens[,100]), type = 'l', ylim = c(0,0.015), lwd = 3, ylab = list("Probability Density", font = 4, col = 1, cex = 1.5), xlab = list("Allele Frequency", font = 4, col = 1, cex = 1.5))
lines(freqx, Dens[,50]/sum(Dens[,50]), lwd = 3, col = 2)
lines(freqx, Dens[,5]/sum(Dens[,5]), lwd = 3, col = 4)
legend("top", legend = c("Time", "100", "50", "5"), col = c(NA, 1,2,4), lwd = 3, text.font = 4)


# Plots for some densities
plot(freqx, Dens[,50]/sum(Dens[,50]), type = 'l', ylim = c(0,1), lwd = 3, ylab = list("Probability Density", font = 4, col = 1, cex = 1.5), xlab = list("Allele Frequency", font = 4, col = 1, cex = 1.5))
lines(freqx, Dens[,1]/sum(Dens[,1]), lwd = 3, col = 4)
lines(freqx, Dens[,50]/sum(Dens[,50]), lwd = 3, col = 1)
lines(freqx, Dens[,2]/sum(Dens[,2]), lwd = 3, col = 2)
legend("topleft", legend = c("Time", "50", "2", "1"), col = c(NA, 1,2,4), lwd = 3, text.font = 4)


#++++++++++++++++++++++++++++++++++++++++++++++++++++++++++++++++++
```
