## Supplementary material for "Models for Eco-evolutionary Extinction Vortices and their Detection": R scripts for regenerating data used in the manuscript: ParameterSpaceSubmit.R

```
# ===========This script simulates parameter space for heterozygote advantage and fluctuating selection
#========== This script also provide an example of fitness trajectory mainly used in the main paper

# packages for plotting. May not be mandatory 
library("viridis", lib.loc="~/R/x86_64-pc-linux-gnu-library/3.2")
library("plot3D", lib.loc="~/R/x86_64-pc-linux-gnu-library/3.2")

Het_Adv_space = function(rate, se, n, p0, fr){
  
H_rsn = array(0,dim = c(length(rate), length(se), length(fr)))
for (i in 1:length(rate)){
  for ( j in 1:length(se)){
    for (k in 1:length(fr)){
      H_rsn[i,j,k] = log(exp(-rate[i]*(1-fr[k]))*(1-se[j])^(-n))/(log((1-0.5*se[j])/(1-se[j])))
    }
  }
}

return(H_rsn)
}

rate = log(seq(1.0, 3, 0.001))# per-capita growth rate
se = seq(0,0.01,0.0001) # selection coefficients
n = 100 # number of polymorphic loci
percent = 0.2# parameter for fluctuating selection denoted as c in the main text, methods section
fr = seq(0,0.2,0.1) # fraction of individuals left after total loss of polymorphic loci. here, we consider 0 throught the paper.

PARASPACEHET = Het_Adv_space(rate, se, n, p0, fr) 

# Ploting parameter space for Heterozygote advantage 
f = 1 # Here, we take a case where the fraction of individuals (N/K) that survive after loss of polymorphic loci to be 0
image(exp(rate),se,PARASPACEHET[,,f], zlim = c(0,n), add = F, col = rev(adjustcolor(heat.colors(n), alpha.f = 1)), ylab = list("Selection Coefficient", font = 4, col = 1), xlab = list("Fertility, exp(rate)", font = 4, col = 1))
image(exp(rate),se,PARASPACEHET[,,f], zlim = c(n,1000*n), add = T, col = rev(adjustcolor(gray.colors((n)), alpha.f =0.9)))
image(exp(rate),se,PARASPACEHET[,,f], zlim = c(-100*n,0), add = T, col = adjustcolor(3, alpha.f =0.2))
contour(exp(rate),se,PARASPACEHET[,,f], drawlabels = T, zlim = c(0,n), labcex = 1.3, lwd = 3, col = 4, nlevels = 5, add = T, method = "flattest", vfont = c("sans serif", "plain"))
text(1.25,0.008, "Region I\n Extinction", cex = 2, srt = 38, font = 4)
text(1.7,0.007, "Region II\n Extinction", cex = 2, font = 4)
text(2.3,0.004, "Region III\n Persistence", cex = 2, font = 4)

points(c(1.2), c(0.005), pch = 16, cex = 3, col = c(1))
points(c( 1.5), c(0.005), pch = 16, cex = 3, col = c(2))
points(c( 1.8), c( 0.005), pch = 16, cex = 3, col = c(3))
points(1.65,0.005, cex = 2, pch = 15, col = 'brown')

text(c(1.2), c(0.0054), "1", cex = 1.5, font = 2)
text(c( 1.5), c(0.0054), "2", cex = 1.5, font = 2)
text(c( 1.8), c( 0.0054), "3", cex = 1.5, srt = 360, font = 2)
text(c( 1.65), c( 0.0054), "b", cex = 1.5, srt = 360, font = 2)


points(c(1.05), c(0.002), pch = 16, cex = 2, col = c(1))
points(c( 1.5), c(0.009), pch = 16, cex = 2, col = c(1))
points(c( 1.8), c( 0.009), pch = 16, cex = 2, col = c(2))
points(1.15,0.002, cex = 2, pch = 16, col = 2)
points(c(1.8), c(0.008), pch = 16, cex = 2, col = c(2))
points(c( 2.0), c(0.008), pch = 16, cex = 2, col = c(2))

text(c(1.05), c(0.0023), "s12", cex = 1.1, font = 2)
text(c( 1.5), c(0.0093), "s11", cex = 1.1, font = 2)
text(c( 1.8), c( 0.0092), "s21", cex = 1.1, srt = 360, font = 2)
text(1.19,0.0023, "s22", cex = 1.1, font = 2)
text(c(1.74), c(0.008), "s23", cex = 1.1, font = 2)
text(c( 1.94), c(0.008), "s24", cex = 1.1, srt = 360, font = 2)


#=====================================================================================
#========== Fluctuating Selection Vortex =============================================
#=====================================================================================
#=========
#===========================================================

Fluct_sel_paraspace = function(rate, se, n, fr, tmax, kapa, percent, p0){
  HHc = array(0, dim = c(length(rate), length(se), length(fr)))
for (ss in 1:length(se)){
  for (r in 1:length(rate)){
    for (ff in 1:length(fr)){
      smaxs = smaxt = se[ss]
      T16 = seq(0,2*pi, length.out = kapa)
      Ti  <- rep(rep(T16, ceiling(tmax/(length(T16)-1))),1)# time points in a season for fluctuating selection
      s <- smaxs*sin(Ti+pi) # selection coeff for genotype AA
      te <- smaxs*sin(Ti)# selection coeff for aa
      
      
      hh1 = 0.5-(percent*sin(Ti))
      
      
      Fit_AA = 1-te 
      Fit_aa = 1-s
      Fit_Aa =  1-hh1*te-(1-hh1)*s     #
      
      Ft_AA = Fit_AA/max(Fit_aa,Fit_Aa, Fit_AA)
      Ft_Aa = Fit_Aa/max(Fit_aa,Fit_Aa, Fit_AA)
      Ft_aa = Fit_aa/max(Fit_aa,Fit_Aa, Fit_AA)  
      
      G3 = exp(mean(log(Ft_aa)))
      G1 = exp(mean(log(Ft_AA)))
      G2 = exp(mean(log(Ft_Aa)))
      
      HHc[r,ss,ff] = log((0.5*(G1+G3))^(-n)*exp(-rate[r]*(1-fr[ff])))/log((p0^2*(G1+2*G2+G3))/(0.5*(G1+G3)))
    }}}

return(HHc)
}


rate = log(seq(1.0, 3, 0.001))# per-capita growth rate
se = seq(0,0.01,0.0001) # selection coefficients
n = 100 # number of polymorphic loci
percent = 0.2# parameter for fluctuating selection denoted as c in the main text, methods section
fr = seq(0,0.2,0.1) # fraction of individuals left after total loss of polymorphic loci. here, we consider 0 throught the paper.
tmax = 100 # maximum time used for calculating geometric mean fitness
p0 = 0.5 # Initial allele frequency for A
kapa = 50 # number of time-steps ina full cycle for fluctuating selection

# calling the function above
PARASPACEFLUCT = Fluct_sel_paraspace(rate, se, n, fr, tmax, kapa, percent, p0)

#Plot for Parameter space for fluctuating selection with recversal of dominance

f = 1# here, we take a case where a 0 fraction of individuals remain after total loss of polymorphic loci
image(exp(rate), se, PARASPACEFLUCT[,,f], zlim = c(0,n),  add = F, col = rev(adjustcolor(heat.colors(n), alpha.f = 1)),ylab = list("Selection Coefficient", font = 4, col = 1), xlab = list("Fertility, exp(rate)", font = 4, col = 1))
contour(exp(rate), se, PARASPACEFLUCT[,,f], drawlabels = T, zlim = c(0,n), labcex = 1.1, lwd = 1.5, col = 4, nlevels = 5, add = T, method = "flattest", vfont = c("sans serif", "plain"))
image(exp(rate), se, PARASPACEFLUCT[,,f], zlim = c(n,1000*n), add = T, col = rev(adjustcolor(gray.colors((n)), alpha.f =0.9)))
image(exp(rate), se, PARASPACEFLUCT[,,f], zlim = c(-1000*n,0), add = T, col = adjustcolor(3, alpha.f =0.2))
#contour(exp(rate), se, HHc[,,f], drawlabels = T, zlim = c(-1000*n,0), col = 3, labcex = 1.5, lwd = 3, nlevels = 10, add = T, method = "flattest", vfont = c("sans serif", "plain"))
text(2.2,0.003, "Region III\n Persistence", cex = 1.20, font = 4)
text(1.5,0.004, "Region II\n Extinction", cex = 1.20, srt = 45, font = 4)
text(1.2,0.008, "Region I\n Extinction", cex = 1.20, font = 4)

points(c(1.40), c(0.005), pch = 16, cex = 3, col = c(1))
points(c( 1.62), c(0.005), pch = 16, cex = 3, col = c(2))
points(c( 1.80), c( 0.005), pch = 16, cex = 3, col = c(3))
text(c(1.40), c(0.0054), "1", cex = 1.5, font = 2)
text(c( 1.62), c(0.0054), "2", cex = 1.5, font = 2)
text(c( 1.8), c( 0.0054), "3", cex = 1.5, srt = 360, font = 2)


points(c(1.3), c(0.007), pch = 16, cex = 2, col = c(1))
points(c( 1.15), c(0.003), pch = 16, cex = 2, col = c(1))
points(c( 2.2), c( 0.0082), pch = 16, cex = 2, col = c(2))
points(2.0,0.0075, cex = 2, pch = 16, col = 2)
points(c(1.3), c(0.0027), pch = 16, cex = 2, col = c(2))

text(c(1.22), c(0.007), "s11", cex = 1.1, font = 2)
text(c( 1.07), c(0.003), "s12", cex = 1.1, font = 2)
text(c( 2.28), c( 0.0082), "s22", cex = 1.1, srt = 360, font = 2)
text(1.92,0.0075, "s21", cex = 1.1, font = 2)
text(c(1.22), c(0.0027), "s23", cex = 1.1, font = 2)


#================== Fitness Fluctuating selection ==================================
#===================================================================================

FitnessFunc = function(tmax, smaxs, smaxt, kapa, percent){
Ti = 1:tmax
s <- smaxs*sin(2*pi*Ti/kapa + pi) # selection coeff for genotype AA
te <- smaxt*sin(2*pi*Ti/kapa)#smaxt*cos(Ti)^2# selection coeff for aa

hh1 = 0.5-percent*sin(2*pi*Ti/kapa)#(smaxs - smaxs*percent/100)*cos(Ti)^10


Fit_AA = (1-te)/(1+smaxt) 
Fit_aa = (1-s)/(1+smaxs)
Fit_Aa =  (1-hh1*te-(1-hh1)*s)/(1+smaxs)     #1 - hh2*Fit_AA - hh1*Fit_aa
return(list(Fit_AA, Fit_aa, Fit_Aa))

}


#parameters
tmax = 100 # Time
smaxs = 0.005 # selection coefficient for a
smaxt = smaxs# selection coefficient for A
kapa = 50 # number of time-steps ina full cycle for fluctuating selection
percent = 0.2

FITNESS = FitnessFunc(tmax, smaxs, smaxt, kapa, percent)

FITNESS_AA = FITNESS[[1]]
FITNESS_aa = FITNESS[[2]]
FITNESS_Aa = FITNESS[[3]]

# ploting fitness vs time
par(mgp=c(2,1,0),mar=c(5,4,4,2)+0.1)
plot(FITNESS_aa, type = 'l', lwd = 3, ylab = list("Fitness", cex = 1.5, font = 4, col = 1), xlab = list("Generation", cex = 1.5, font = 4, col = 1))
lines(FITNESS_AA, lwd = 3, col = 2)
lines(FITNESS_Aa, lwd = 3, col = 4)
lines(rowMeans(cbind(FITNESS_aa, FITNESS_AA, FITNESS_Aa)), lwd = 3, col = 3)
abline(h = 1-smaxs, col = "brown", lty = 2, lwd = 3)
abline(h = 1, col = "blue", lty = 2, lwd = 3)
abline(h = (1+ 1-smaxs + 1-smaxt)/3, col = "green", lty = 2, lwd = 3)
legend("topright", legend = c(expression(w[AA]), expression(w[aa]), expression(w[Aa]), expression(bar(w))), col = c(1,2,4,3), lwd =3, text.font = 4)

#End of this file
```
