## Supplementary material for "Models for Eco-evolutionary Extinction Vortices and their Detection": R scripts for regenerating data used in the manuscript: READMEMEEVD.txt

This folder contains the R scripts for simulating the individual-based model and the and analytic approximation in the manuscript "Models for Eco-evolutionary Extinction Vortices and their Detection". Data from the individual-based model are produced from the ""IndividualBasedModelSubmit.R" script. The data is then analysed using "AnalysisFunctionsSubmit.R" script. For the analytic approximation, the "NumericApproximationSubmit.R" file is used. Further, we provide a short script "ParameterSpaceSubmit.R", used for simulating fitness trajectory examples and dividing our parameter space (Figure 1 and 2 in the manuscript respectively). 

Within the scripts, all the parameters used are described in the comments. The denotation of some parameters in the scripts differ from those used in the manuscript partly because they are generic functions within R. Therefore, we strongly advise to read through the description provided within the script so as to have the right parameter combinations. Also, note that there are other parameters not used in our manuscript and are clearly identified via comments. Note that the provided parameter values in the scripts may only represent one scenario of parameter combinations used in one of the cases in the manuscript. Also, in the respective scripts, we have provided the main code for plotting of the figures in the manuscript at the end of each section or function. The axes limits may have to be manually changed to suit the specific parameter combination(s). 

The individual-based model data was produced on a Linux computing cluster (Ubuntu 16.04.6). Note that some parameter combinations especially involving Case 3 and Control scenario as described in the manuscript when using 100 replicates can take one or more days to finish on a node with 16 cores. To avoid exceeding memory limits in some parameter combinations, we save each case separately as RData files and combine the cases for a given parameter later into a 3-dimensional array as explained in the script comments.   

The produced data was analysed in Linux operating system (Release: 16.04 ), with R version 3.2.3 and has not been tested on any other versions.
